# [1,2,4]Triazolo[3,4-*b*]benzothiazole scaffold as versatile nicotinamide mimic allowing nanomolar inhibition of different PARP enzymes

**DOI:** 10.1101/2022.08.29.505683

**Authors:** Sudarshan Murthy, Maria Giulia Nizi, Mirko M. Maksimainen, Serena Massari, Juho Alaviuhkola, Barbara E. Lippok, Chiara Vagaggini, Sven T. Sowa, Albert Galera-Prat, Yashwanth Ashok, Harikanth Venkannagari, Renata Prunskaite-Hyyryläinen, Elena Dreassi, Bernhard Lüscher, Patricia Korn, Oriana Tabarrini, Lari Lehtiö

**Affiliations:** Biocenter Oulu and Faculty of Biochemistry and Molecular Medicine, University of Oulu, 90220 Oulu, Finland; Department of Pharmaceutical Sciences, University of Perugia, 06123 Perugia, Italy; Institute of Biochemistry and Molecular Biology, RWTH Aachen University, 52074 Aachen, Germany; Department of Biotechnology, Chemistry and Pharmacy, University of Siena, I-53100, Siena, Italy

## Abstract

Here we report [1,2,4]triazolo[3,4-*b*]benzothiazole (TBT) as a new inhibitor scaffold, which competes with nicotinamide in the binding pocket of human poly- and mono-ADP-ribosylating enzymes. The binding mode was studied through analogs and their crystal structures with TNKS2, PARP2, PARP14 and PARP15. Based on the substitution pattern, we were able to identify The 3-amino derivatives **21** (OUL243) and **27** (OUL232), as inhibitors of mono-ARTs PARP7, PARP10, PARP11, PARP12, PARP14 and PARP15 at nM potencies, with compound **27** being the most potent PARP10 inhibitor described to date with an IC_50_ of 7.8 nM and the first PARP12 inhibitor ever reported. On the contrary, hydroxy derivative **16** (OUL245) inhibits poly-ARTs with a selectivity towards PARP2. The scaffold does not possess inherent cell toxicity and the inhibitors can enter cells and engage with the target protein. This, together with favorable ADME properties, demonstrates the potential of the TBT scaffold for future drug development efforts towards selective inhibitors against specific enzymes.

## INTRODUCTION

ADP-ribosylation is a post-translational modification found in bacteria and eukaryotes and it is also associated with viral and bacterial infections. The human diphtheria toxin-like ARTD family contains PARP and tankyrase (TNKS) enzymes that can catalyze both mono-ADP-ribosylation (MAR, mono-ARTs) as well as generate elongated and branched chains of poly-ADP-ribose (PAR, poly-ARTs).^1^ The PARPs and TNKSs form a family of structurally and functionally diverse enzymes, which are involved in the regulation of various key biological and pathological processes such as DNA repair, cell differentiation, gene transcription, and signal transduction pathways.^2–4^ PARPs and TNKSs use nicotinamide adenine dinucleotide, NAD^+^, to transfer an ADP-ribose (ADPr) unit onto target proteins or nucleic acids with a release of nicotinamide. The transfer of ADPr in proteins occurs onto amino acid side chains with a nucleophilic oxygen, nitrogen, or sulfur resulting in O-, N-, or S-glycosidic linkage to the ADP-ribose. This can be further extended to PAR by poly-ARTs PARP1-2 and TNKS1-2.^5,6^ Poly-ARTs contain a triad of amino acids H-Y-E in their active sites. H-Y are important for binding the NAD^+^, while E stabilizes the oxocarbenium ion transition state and enables the elongation of the ADP-ribose chain by activating the ribose 2’-hydroxyl group.^7^ However, the H-Y-E motif is not an absolute indicator determining the PARylation activity as there are two enzymes, PARP3 and PARP4 having the H-Y-E motif but appear to be unable to produce PAR chains.^8,9^

Over the past decades PARPs have emerged as drug targets due to their roles in critical cellular processes.^10^ Especially the discovery of synthetic lethality of PARP1 inhibition in the context of BRCA deficient cancers^11,12^ boosted inhibitor development and led to the first approved drug, olaparib, in 2014. Other PARP1/2 inhibitors, including rucaparib, niraparib and talazoparib have also entered clinical applications for the treatment of ovarian and breast cancers deficient in homologous recombination-mediated DNA double-strand break repair.^13–15^ TNKSs have also emerged as promising drug targets especially due to their role in the Wnt/β-catenin signaling^16–18^ with two compounds, E7449 and STP1002 (structure undisclosed), having proceeded into clinical studies along with other promising compounds, such as OM-153,^19^ which is in advanced pre-clinical testing (Figure 1). The patent literature on poly-ART inhibitors has been expanding and also recently reviewed.^20,21^

**Figure 1.**
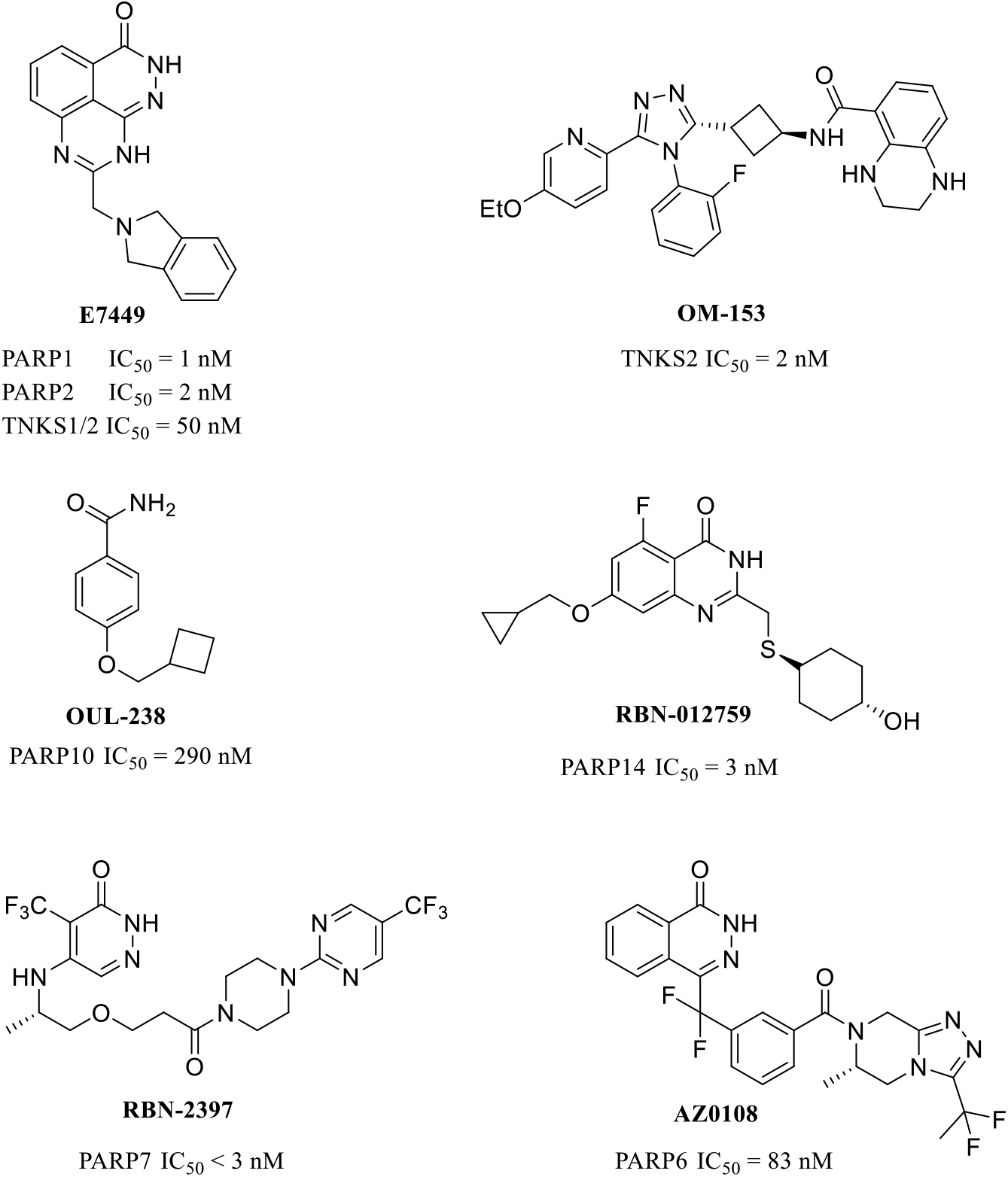
Examples of the most potent PARP inhibitors.

The majority of mono-ARTs (PARP6-16) have a different active site triad, H-Y-Φ, where Φ represents a hydrophobic amino acid, such as isoleucine, leucine, or tyrosine. Especially the lack of glutamate has been linked to the activity being limited to MARylation. Notably, PARP13 is thought to be inactive^22^ and PARP9 is modulating ADP-ribosylation activity of the E3 ubiquitin ligase DTX3L.^23–25^ Although understudied until recently, roles of mono-ARTs in controlling signaling events in cells along with the recent discovery of their implications in many diseases, make them new possible drug targets. This has resulted in the interest in developing small molecule inhibitors, precious research tools, which can be used in parallel with biochemical methods to validate the cellular functions of these enzymes.

PARP10 was the first enzyme of the family described as a mono-ART^26^ and its inhibitor, OUL35 described earlier by us,^27^ could rescue HeLa cells from PARP10-induced cell death and sensitize HeLa cells to DNA damage in agreement with knockdown studies.^28,29^ Also patient cells deficient in PARP10 were shown to be sensitized to DNA damage.^30^ Subsequently, multiple studies have reported additional PARP10 inhibitors,^31–35^ such as **OUL238**^35^ which is one of the most potent compounds (Figure 1).

In addition to PARP10, inhibitors have also been developed for other mono-ARTs. PARP14 mediates gene transcription through its MARylation activity and it has been implied as a possible therapeutic target for example in lymphoma, myeloma, hepatocellular carcinoma and in prostate cancer.^36–38^ At the moment, the most advanced inhibitor for PARP14 is RBN-012759 (Figure 1), disclosed by Ribon Therapeutics with an IC_50_ of 3 nM.^39^ Ribon Therapeutics has also developed RBN-2397, a PARP7 selective inhibitor with < 3 nM IC_50_ also demonstrating antitumor effects in xenografts, currently progressed to phase I clinical trials.^40^ Also AstraZeneca has contributed to the inhibitor development against mono-ART by describing a potent PARP6 inhibitor, AZ0108 (Figure 1), which prevents ADP-ribosylation of Chk1 that subsequently contributes to anti-tumor effects in breast cancer mouse models.^41^ Recently, the first inhibitors for PARP15 have been described although the cellular role of this enzyme is not yet well elucidated.^42,43^ The efforts summarized above were recently reviewed documenting the high and increasing interest in the development of mono-ART inhibitors.^44^

Here we report our contribution in identifying a set of compounds based on a new nicotinamide mimicking chemotype, a [1,2,4]triazolo[3,4-*b*]benzothiazole (TBT), able to inhibit different PARP family enzymes with submicromolar activity depending on the substitution pattern around the central core. The shortlisted compounds were profiled against most of the active human PARP enzymes leading to the most potent inhibitors for PARP10 described to date, with the best compound reaching 7.8 nM IC_50_ in an enzymatic assay. The binding mode of the TBT scaffold was studied through the synthesis of analogs and their complex crystal structures with PARP2, TNKS2, PARP14 and PARP15. We demonstrate that the compounds enter cells and engage with the target proteins with the most potent compound showing 150 nM EC_50_ value and the scaffold does not possess inherent cell toxicity. In addition, *in vitro* ADME studies show that the compounds have a good solubility in the 50-150 µM range, are extremely stable in polar solvents and human plasma, and are not susceptible to first pass metabolism by enzymes of the human microsomes. The TBT scaffold therefore forms a basis for drug development efforts towards multiple enzymes of the family.

## RESULTS

### Biochemical analysis and structural studies of OUL40 (1) and analogs design

We previously screened a compound library from the open chemical repository of the National Cancer Institute, which led to the identification of the potent and selective PARP10 inhibitor OUL35.^27^ From the same screening, also a TBT derivative OUL40 (NSC295701) (**1**) (Figure 2A) emerged, showing an IC_50_ = 3.2 µM against PARP10. This led us to hypothesize that **1** could be a new nicotinamide mimicking compound, with the potential to inhibit multiple PARPs. Indeed, when assayed against a panel of two poly-ARTs, PARP2 and TNKS2, and two additional mono-ARTs, PARP14 and PARP15, IC_50_ values in the low micromolar range (1.2-5.3 µM) were obtained (Table 1). With X-ray crystallography, we confirmed the binding of **1** into the nicotinamide binding pocket of TNKS2, PARP14 and PARP15 active sites. The inhibitor binding mode is highly similar in all three enzymes, where two hydrogen bonds are generated between the N1 and N2 of the triazole ring with glycine and serine residues, respectively. In addition, π-π interactions are present between the inhibitor core and tyrosine residues (Figure 2B-D).

**Figure 2:**
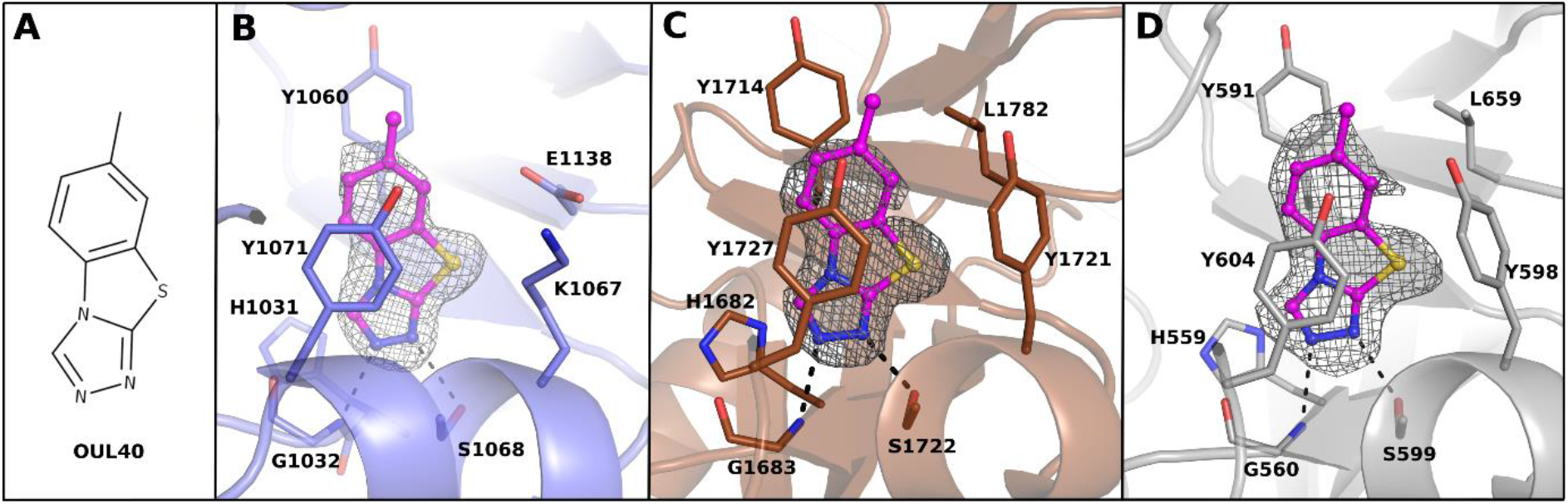
New nicotinamide mimicking compound. **(A)** The structural formula of compound **1. (B)** TNKS2 **(C)** PARP14, and **(D)** PARP15 crystal structures in complex with **1**. TNKS2, PARP14 and PARP15 are colored in blue, brown and grey, respectively. **1** is presented as a ball-and-stick model and colored in magenta. The hydrogen bonds are indicated with black dashes. The ligand-omitted sigma A weighted Fo-Fc electron density maps are colored in grey and contoured at 3.0 σ.

**Table 1.**
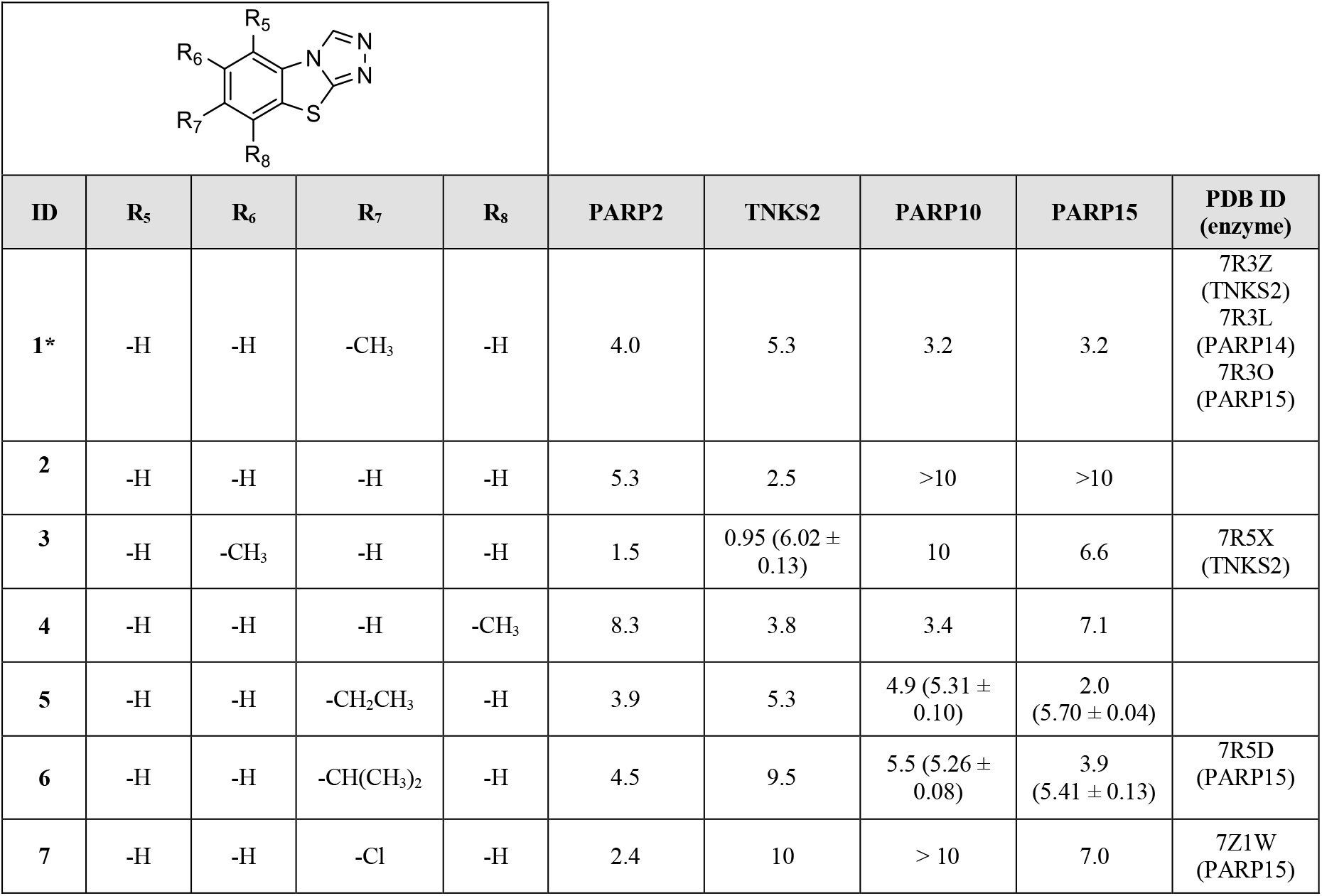
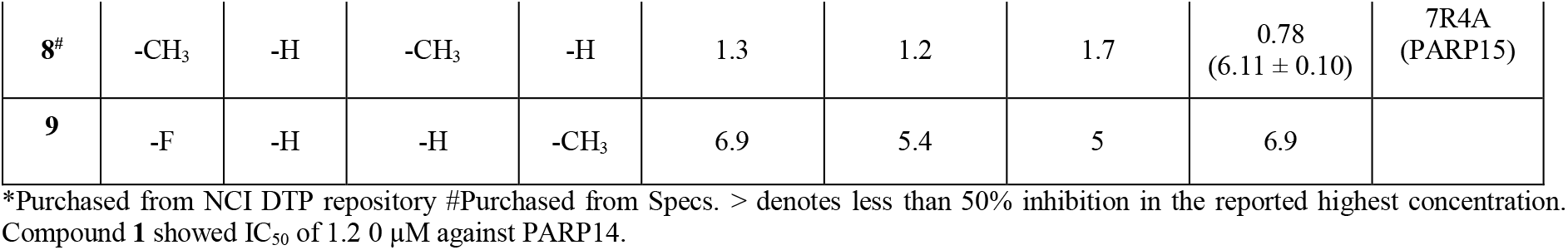
Activity of **1** and the initial analogs. IC_50_ (pIC_50_±SEM) values (µM) and PDB IDs are reported.

|  |  |  |  |  |  |  |  |  |  |
| --- | --- | --- | --- | --- | --- | --- | --- | --- | --- |
| ID | R<sub>5</sub> | R<sub>6</sub> | R<sub>7</sub> | R<sub>8</sub> | PARP2 | TNKS2 | PARP10 | PARP15 | PDB ID (enzyme) |
| **1\*** | -H | -H | -CH<sub>3</sub> | -H | 4.0 | 5.3 | 3.2 | 3.2 | 7R3Z (TNKS2)<br>7R3L (PARP14)<br>7R3O (PARP15) |
| **2** | -H | -H | -H | -H | 5.3 | 2.5 | >10 | >10 |  |
| **3** | -H | -CH<sub>3</sub> | -H | -H | 1.5 | 0.95 (6.02 ± 0.13) | 10 | 6.6 | 7R5X (TNKS2) |
| **4** | -H | -H | -H | -CH<sub>3</sub> | 8.3 | 3.8 | 3.4 | 7.1 |  |
| **5** | -H | -H | -CH<sub>2</sub>CH<sub>3</sub> | -H | 3.9 | 5.3 | 4.9 (5.31 ± 0.10) | 2.0 (5.70 ± 0.04) |  |
| **6** | -H | -H | -CH(CH<sub>3</sub>)<sub>2</sub> | -H | 4.5 | 9.5 | 5.5 (5.26 ± 0.08) | 3.9 (5.41 ± 0.13) | 7R5D (PARP15) |
| **7** | -H | -H | -Cl | -H | 2.4 | 10 | > 10 | 7.0 | 7Z1W (PARP15) |
| **8<sup>#</sup>** | -CH<sub>3</sub> | -H | -CH<sub>3</sub> | -H | 1.3 | 1.2 | 1.7 | 0.78<br>(6.11 ± 0.10) | 7R4A<br>(PARP15) |
| **9** | -F | -H | -H | -CH<sub>3</sub> | 6.9 | 5.4 | 5 | 6.9 |  |

\*Purchased from NCI DTP repository #Purchased from Specs. > denotes less than 50% inhibition in the reported highest concentration. Compound **1** showed IC<sub>50</sub> of 1.20 μM against PARP14.

Our preliminary studies on **1** revealed the potency of the compound scaffold by showing reasonable inhibition even in the absence of the typical benzamide moiety. Importantly, no PARP inhibitor based on this tricyclic scaffold has been reported until now. As it offered many options for substitutions, we were encouraged to study this scaffold in more detail.

At first, we were interested to test whether small modifications of the scaffold would lead to any significant changes in selectivity towards mono- or poly-ARTs, or to specific inhibition of individual ARTs within the respective subfamily. Iterative medicinal chemistry cycles were performed with a first set of compounds that emerged by working on the benzene ring, where the methyl group of **1** was deleted (**2**), moved from C-7 to C-6 or C-8 positions (**3** and **4**), or replaced by bulkier groups as in compounds **5-7**. Disubstituted derivatives **8** and **9** were also contemplated (Table 1). Within further derivatives monomethoxy (**10-13**), dimethoxy (**14** and **15**) monohydroxy (**16** and **17**), and dihydroxy (**18**) groups are decorating the benzene ring (Table 2).

**Table 2.**
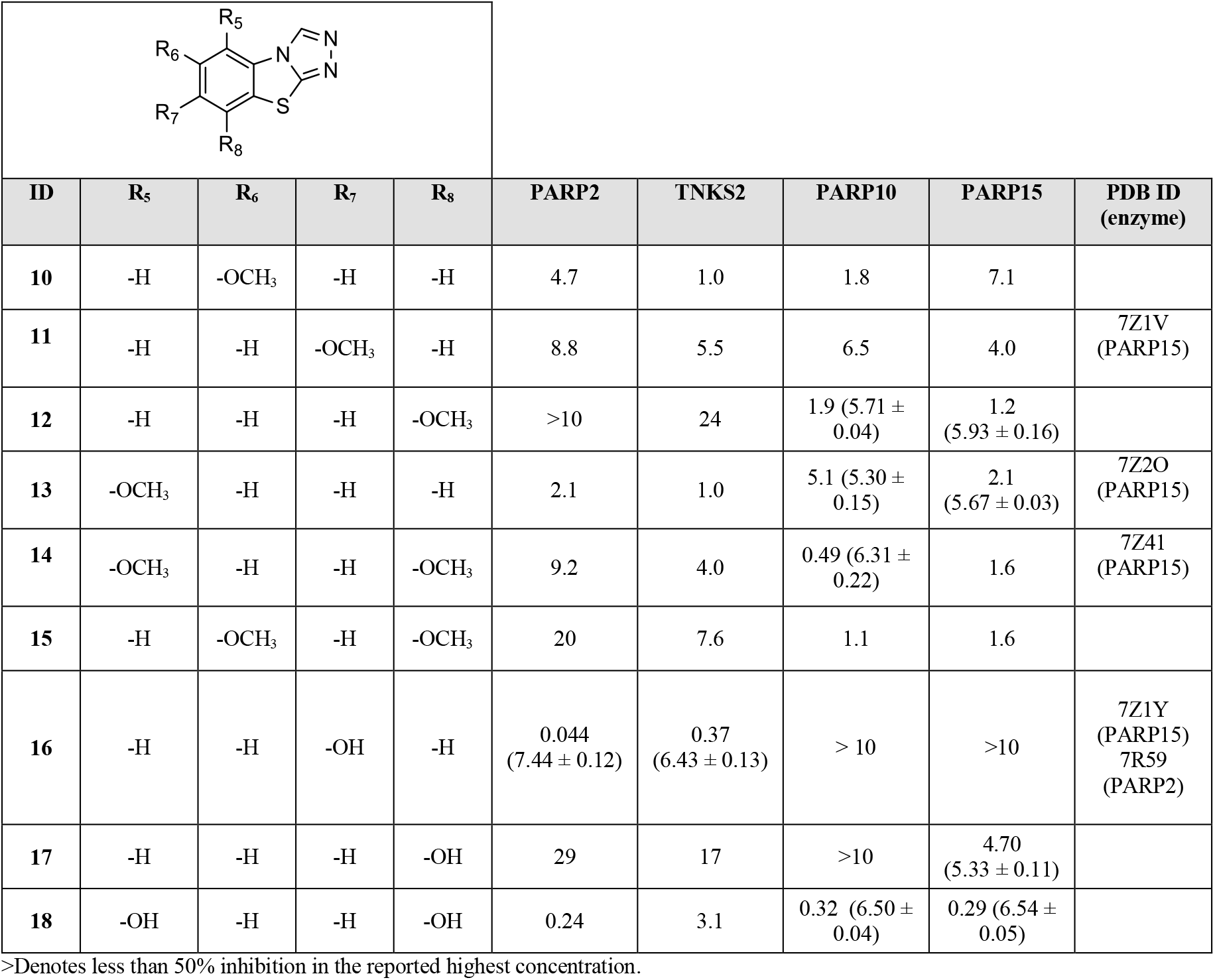
Activity of methoxy and hydroxy substituted analogs. IC_50_ (pIC_50_ ± SEM) values (µM) and PDB IDs are reported.

|  |  |  |  |  | PARP2 | TNKS2 | PARP10 | PARP15 | PDB ID (enzyme) |
| --- | --- | --- | --- | --- | --- | --- | --- | --- | --- |
| ID | R<sub>5</sub> | R<sub>6</sub> | R<sub>7</sub> | R<sub>8</sub> |  |  |  |  |  |
| 10 | -H | -OCH<sub>3</sub> | -H | -H | 4.7 | 1.0 | 1.8 | 7.1 |  |
| 11 | -H | -H | -OCH<sub>3</sub> | -H | 8.8 | 5.5 | 6.5 | 4.0 | 7Z1V (PARP15) |
| 12 | -H | -H | -H | -OCH<sub>3</sub> | >10 | 24 | 1.9 (5.71 ± 0.04) | 1.2 (5.93 ± 0.16) |  |
| 13 | -OCH<sub>3</sub> | -H | -H | -H | 2.1 | 1.0 | 5.1 (5.30 ± 0.15) | 2.1 (5.67 ± 0.03) | 7Z2O (PARP15) |
| 14 | -OCH<sub>3</sub> | -H | -H | -OCH<sub>3</sub> | 9.2 | 4.0 | 0.49 (6.31 ± 0.22) | 1.6 | 7Z41 (PARP15) |
| 15 | -H | -OCH<sub>3</sub> | -H | -OCH<sub>3</sub> | 20 | 7.6 | 1.1 | 1.6 |  |
| 16 | -H | -H | -OH | -H | 0.044 (7.44 ± 0.12) | 0.37 (6.43 ± 0.13) | > 10 | >10 | 7Z1Y (PARP15)<br>7R59 (PARP2) |
| 17 | -H | -H | -H | -OH | 29 | 17 | >10 | 4.70 (5.33 ± 0.11) |  |
| 18 | -OH | -H | -H | -OH | 0.24 | 3.1 | 0.32 (6.50 ± 0.04) | 0.29 (6.54 ± 0.05) |  |

>Denotes less than 50% inhibition in the reported highest concentration.

Subsequent biochemical and structural analyses described later suggested that the C-3 functionalization of the triazole ring could improve selectivity. Indeed, additional compounds were prepared by placing a heteroatom, oxygen, sulfur or nitrogen in this position, which was also derivatized while maintaining a 7-methyl (**19-25**) or a 5,8-dimethoxy (**26-31**) substitution pattern in the benzene ring (Table 3).

**Table 3.**
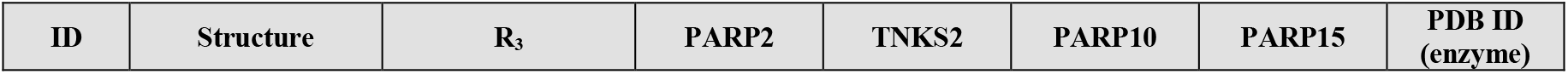
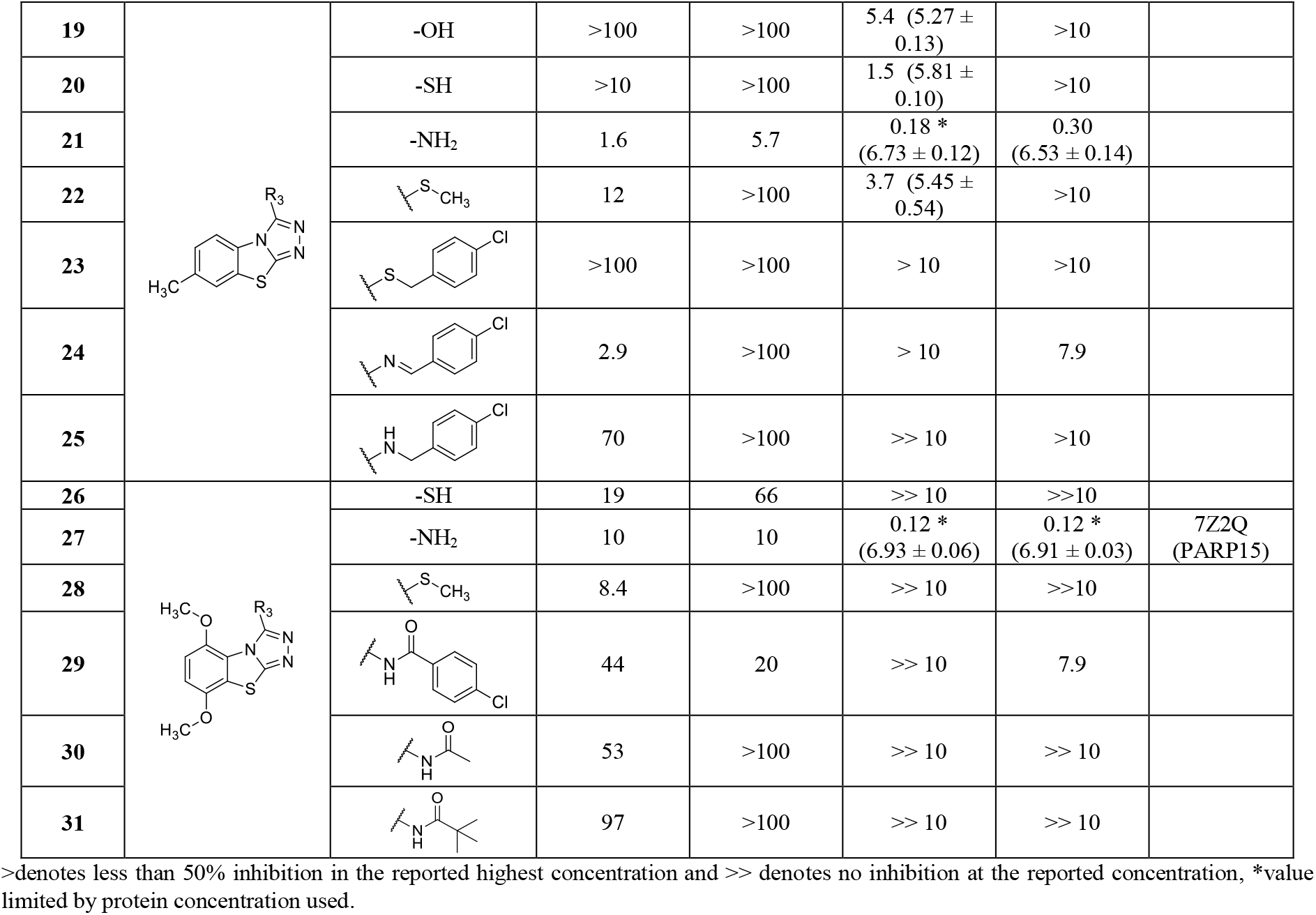
Activity of C-3 substituted analogs. IC_50_ (pIC_50_ ± SEM) values (µM) and PDB IDs are reported.

| ID | Structure | R<sub>3</sub> | PARP2 | TNKS2 | PARP10 | PARP15 | PDB ID (enzyme) |
| --- | --- | --- | --- | --- | --- | --- | --- |
| 19 |  | -OH | >100 | >100 | 5.4 (5.27 ± 0.13) | >10 |  |
| 20 |  | -SH | >10 | >100 | 1.5 (5.81 ± 0.10) | >10 |  |
| 21 |  | -NH<sub>2</sub> | 1.6 | 5.7 | 0.18 *<br>(6.73 ± 0.12) | 0.30<br>(6.53 ± 0.14) |  |
| 22 |  |  | 12 | >100 | 3.7 (5.45 ± 0.54) | >10 |  |
| 23 |  |  | >100 | >100 | > 10 | >10 |  |
| 24 |  |  | 2.9 | >100 | > 10 | 7.9 |  |
| 25 |  |  | 70 | >100 | >> 10 | >10 |  |
| 26 |  | -SH | 19 | 66 | >> 10 | >>10 |  |
| 27 |  | -NH<sub>2</sub> | 10 | 10 | 0.12 *<br>(6.93 ± 0.06) | 0.12 *<br>(6.91 ± 0.03) | 7Z2Q<br>(PARP15) |
| 28 |  |  | 8.4 | >100 | >> 10 | >>10 |  |
| 29 |  |  | 44 | 20 | >> 10 | 7.9 |  |
| 30 |  |  | 53 | >100 | >> 10 | >> 10 |  |
| 31 |  |  | 97 | >100 | >> 10 | >> 10 |  |

>denotes less than 50% inhibition in the reported highest concentration and >> denotes no inhibition at the reported concentration, \*value limited by protein concentration used.

### Chemistry

All the compounds, with the exception of derivatives **1** and **8** that are commercially available, were synthesized as shown in Schemes 1 and 2. In particular, as depicted in Scheme 1, TBT target compounds variously functionalized on the benzene ring were prepared from the key 2-hydrazinobenzothiazole intermediates **33, 64-73**, and **80-82**, obtained through three different synthetic pathways.

**Scheme 1a.**
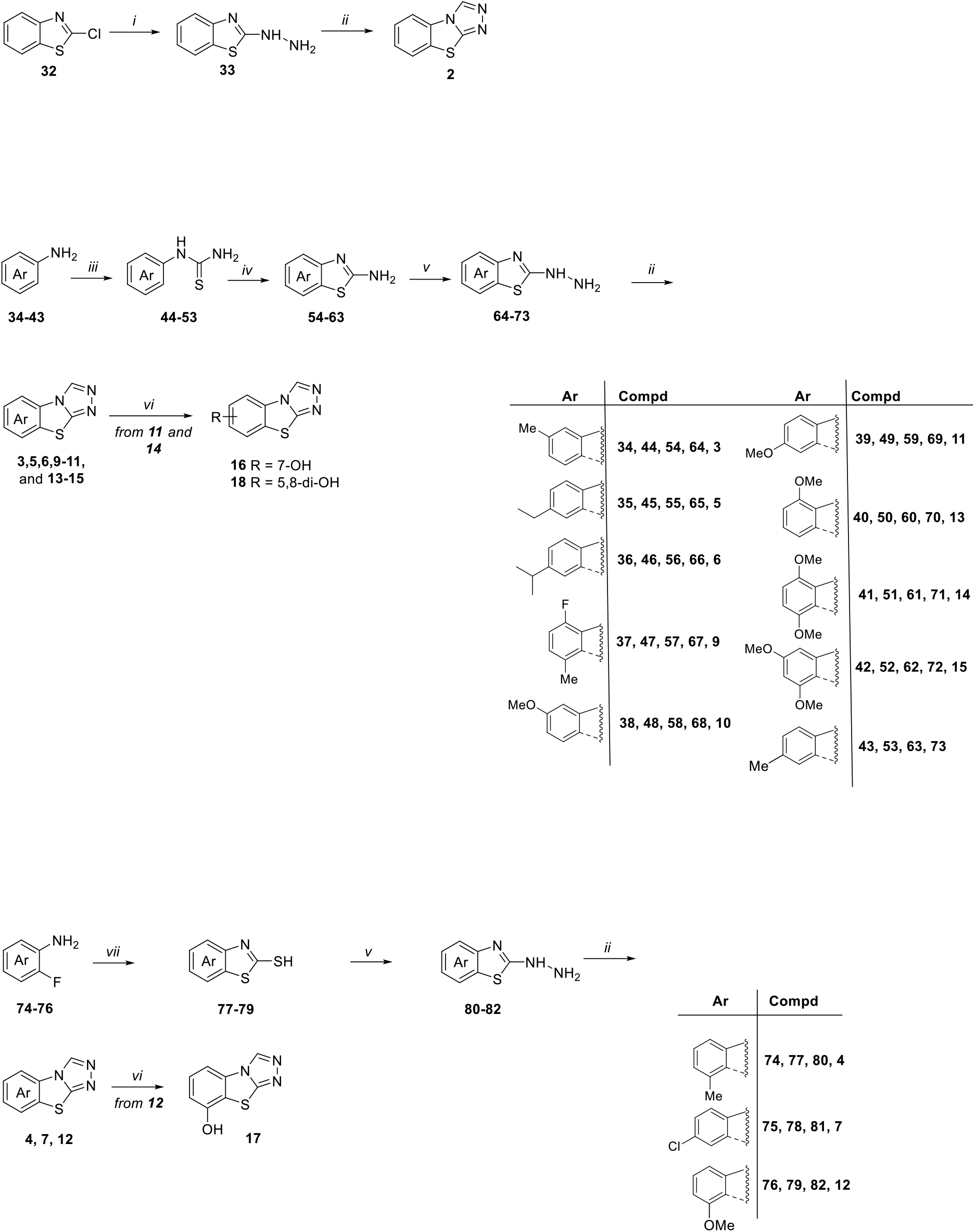
^*a*^ *Reagents and conditions: i*) hydrazine hydrate, EtOH, 80 °C, overnight; *ii)* formic acid, reflux, 7-48 h; *iii*) NH_4_SCN, H_2_O, 12 N HCl, reflux, 6-48 h; *iv*) Br_2_, CHCl_3_, r.t., 2-8 h; *v*) hydrazine hydrate, CH_3_COOH, ethylene glycol, 125 °C, 7-48 h; *vi*) BBr_3_, dry CH_2_Cl_2_, r.t., 3 h; *vii*) potassium ethyl xanthogenate, dry DMF, 110 °C; 3 h.

**Scheme 2a.**
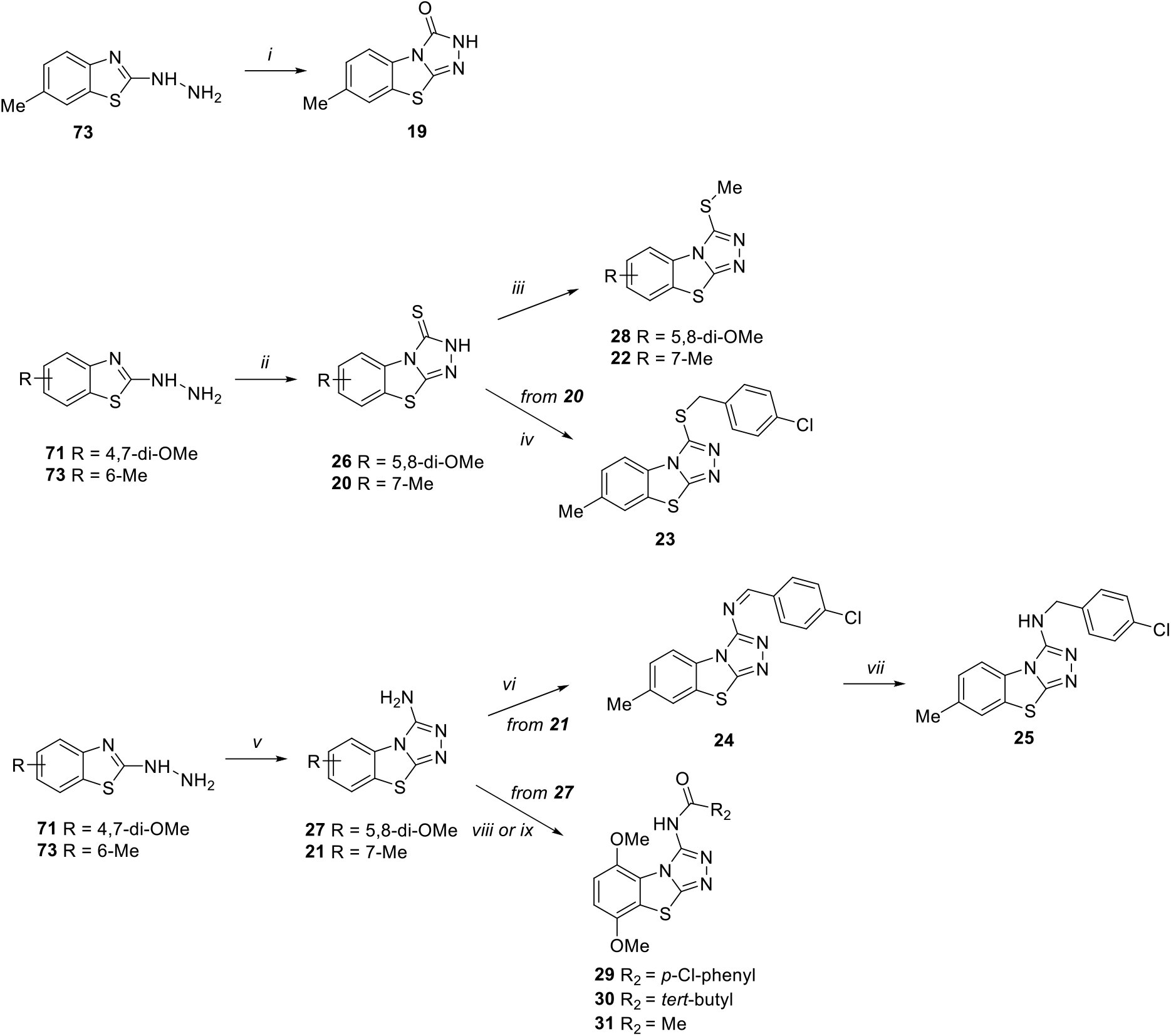
^*a*^ *Reagents and conditions:* i) urea, neat, fusion, 3 h; *ii*) CS_2_, KOH, EtOH, reflux, 2h; *iii*) MeI, K_2_CO_3_, dry DMF, 80 °C, 2 h; *iv*) *p-*chlorobenzyl chloride, EtOH, reflux, 4 h; *v*) CNBr, MeOH, reflux, 3 h; *vi*) *p*-chlorobenzaldehyde, *p*-TsOH, dry benzene, reflux, 16 h; *vii*) NaBH_4_, absolute EtOH, r.t., overnight; *viii*) *p-*chlorobenzyl chloride, Et_3_N, dry DMF, 80 °C, 2 h; *ix)* trimethylacetyl chloride or acetyl chloride Et_3_N, dry toluene, 110 °C, 12 h.

The unsubstituted hydrazinobenzothiazole **33** was obtained starting from 2-chlorobenzothiazole **32** by reaction with hydrazine hydrate in EtOH with a conversion of 98%. Most of the 2-hydrazinobenzothiazoles (**64-73**) were instead synthesized starting from the properly substituted anilines **34-43**, which were converted into the corresponding arylthiourea derivatives **44-53** by reaction with NH_4_SCN in acidic solution at reflux. The successive oxidative cyclization of **44-53** using Br_2_ gave the corresponding 2-aminobenzothiazoles **54-63**, which were then treated with hydrazine hydrate to give **64-73**. On the other hand, 2-hydrazinobenzothiazoles **80-82** were prepared from 2-mercaptobenzothiazoles **77-79**, in turn obtained through a double nucleophilic substitution of properly substituted 2-fluoroanilines **74-76** with potassium ethyl xanthogenate in dry DMF.

The successive reaction of the 2-hydrazinobenzothiazoles with refluxed formic acid in excess led to the synthesis of tricyclic compounds: **2** starting from **33**; **3, 5, 6, 9-11** and **13-15** from **64-72**; while **4, 7, 12** from **80-82**.

The methoxy derivatives **11, 12**, and **14** were further elaborated into the corresponding hydroxyl derivatives **16-18** by using BBr_3_.

As reported in Scheme 2, TBT variously functionalized at the C-3 position **19-31**, were synthesized from 4,7-dimethoxy and 6-methyl hydrazine intermediates **71** and**73**. By treating **73** with urea in neat condition and at the fusion temperature (133 °C), benzothiazol-3-one **19** was obtained.

Starting from both **71** and **73** and using CS_2_ in EtOH, benzothiazole-3-thiones **26** and **20** were obtained, respectively. S-alkylation of **26** and **20** with MeI, in the presence of K_2_CO_3_ in dry DMF gave the corresponding derivatives **28** and **22**. Compound **20** was also S-alkylated by reaction with *p-*chlorobenzyl chloride in EtOH to give compound **23**.

Finally, the reaction of **71** and **73** with CNBr furnished 3-aminobenzothiazole derivatives **27** and **21**. 3-Amino-7-methyl derivative **21** was condensed with *p*-chlorobenzaldehyde to give imine derivative **24**, which was then reduced to amine derivative **25** with NaBH_4_. Amidation of derivative **27** with *p*-chlorobenzoyl chloride in the presence of Et_3_N in dry DMF gave compound **29**, while the reaction of **27** with trimethylacetyl chloride or acetyl chloride in the presence of Et_3_N in dry toluene yielded compounds **30** and **31**, respectively.

### OUL40 (1) analogs: biochemical analysis and structural studies

All the synthesized TBTs were initially tested against representative members of the PARP family: two poly-ARTs, PARP2 and TNKS2, and two mono-ARTs, PARP10 and PARP15. The latter were selected based on availability of a robust cell-based readout for PARP10 engagement and for a similarly robust crystal system of PARP15 to study compound binding modes experimentally. In addition, all the analogs were also routinely tested for the toxicity using a colorimetric WST-1 assay, which only identified the isopropyl derivative **6** and dihydroxy derivative **18** as being toxic in a dose-dependent manner (Figure S1).

We first tested the effect of small alkyl groups and halogens on the benzene ring (Table 1). The only commercially available compound **8** had an additional methyl substituent and it showed improved potency against all the tested PARPs. The removal of the methyl group on the other hand reduced the potency of **2** especially against PARP10 and PARP15 (IC_50_ >10 µM) indicating that an electron donating hydrophobic substituent would be important for potency towards mono-ARTs (Table 1). Shifting of the methyl group to other positions did not have major effects on the potency (**3, 4** and **9**) and all the compounds maintained µM potencies for the tested enzymes. Compound **3** having the methyl in the C-6 position, however, showed higher potency against PARP2 and TNKS2. The TNKS2 crystal structure in complex with **3** revealed that the 6-methyl pushed Tyr1050 to different conformation and provided the additional interaction explaining the poly-ART selectivity (Figure S2A-B). When C-7 methyl found in **1** was extended to a larger alkyl (**5** and **6**), no improvements in inhibition potency was observed, while the C-7 chlorine derivative **7** maintained a micromolar activity only against PARP2 and PARP15. The minor modifications of the C-6 substituent did not result in significant structural changes as observed from the PARP15 crystal structures in complex with **6, 7** and **8**, which showed highly similar binding modes to **1** (Figure S2C-E).

Next, we tested effects of hydroxy and methoxy groups placed in various positions of the benzene ring of the TBT scaffold (Table 2). Interestingly, the presence of a C-7 hydroxy group made **16** very potent and specific for poly-ARTs with nearly ten-fold selectivity for PARP2 over TNKS2 (IC_50_ of 44 nM versus 370 nM). Based on the comparison of complex structures of PARP2 and PARP15 (Figure 3), **16** has a similar binding mode as **1**, but the catalytic residue of PARP2 (Glu558), not present in PARP15, interacts with the hydroxyl group of **16** (Figure 3A). Hydroxyl group also interacts with Met456 backbone amide via a water molecule. In contrast, in PARP15 the hydroxyl group interacts with the carbonyl of Ala583 causing a compound orientation that brings the sulfur atom of **16** in a close contact with the sidechain of Tyr598, which has therefore changed its conformation compared to the binding mode of **1** (Figure 2D and 3B). In addition, the ligand-omitted Fo-Fc electron density map is not well-defined indicating flexibility in the binding mode of **16** (Figure 3B).

**Figure 3:**
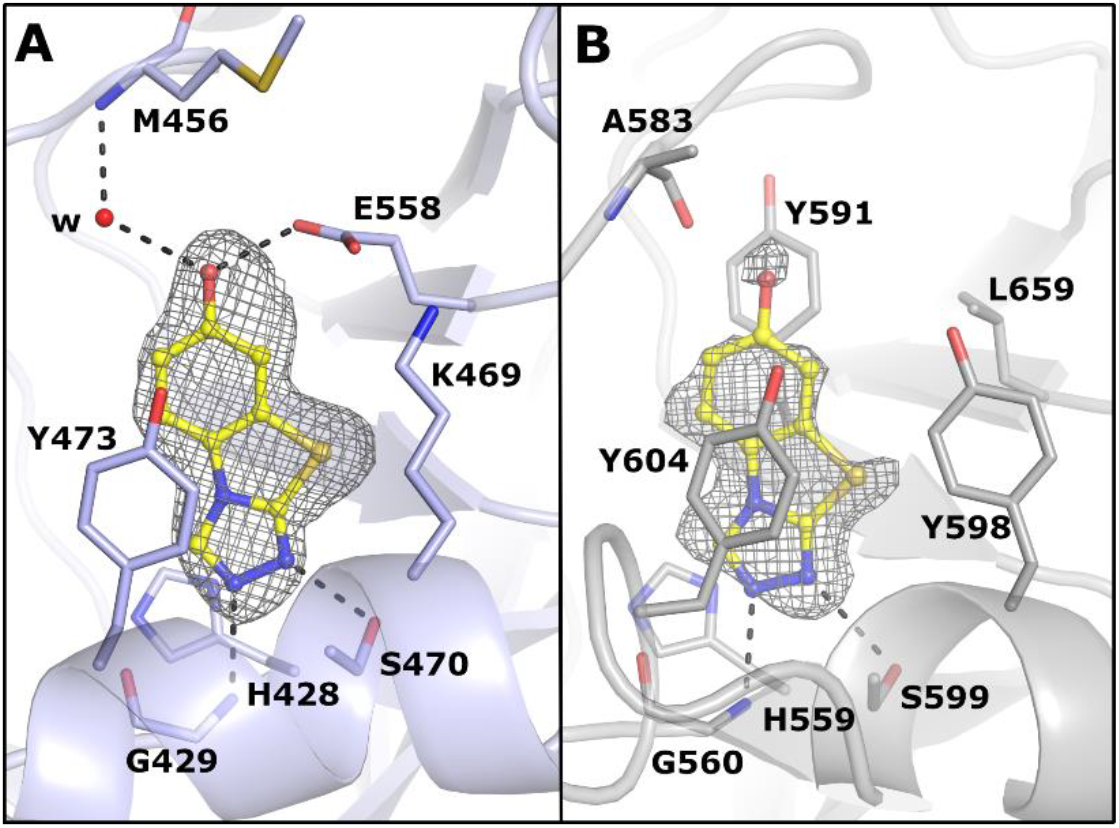
Binding mode of **16** between poly-ART and mono-ART showed by the **(A)** PARP2 and **(B)** PARP15 complex crystal structures. The ligand is presented as a ball-and-stick model and colored in yellow. The hydrogen bonds are indicated with black dashes. The ligand-omitted sigma A weighted Fo-Fc electron density maps are colored in grey and contoured at 3.0 σ.

By shifting the hydroxy group from C-7 to C-8 position, a very different profile was shown by compound **17** that maintained a weak activity only against PARP15. In contrast to the hydroxy of **16**, the replacement of the methyl group of **1** with a methoxy gave compound **11** endowed with a similar profile and a similar binding mode (Figure S2F). Dimethoxy derivative **14** showed submicromolar potency against PARP10 (IC_50_ = 490 nM) and the corresponding *di*-hydroxy analogue **18** expanded the nanomolar potency also against PARP15 and PARP2. This did not provide us the poly-ART vs. mono-ART selectivity, but the presence of an 8-methoxy group made **12** selective against mono-ARTs PARP10 and PARP15. The PARP15 crystal structure in complex with the selective PARP10 inhibitor **14** (Figure 4A) revealed plasticity in the compound orientation compared to **1**. The small rotation was observed by comparing the crystal structures (Figure 4B), which also revealed conformational changes in the side chains of Leu659 and Tyr598. An even more dramatic change was observed in the complex structure with 5-methoxy derivative **13**, which showed a 180° horizontal flip of the compound in comparison to 5,8-dimethoxy **14** (Figure 4C and 4A).

**Figure 4:**
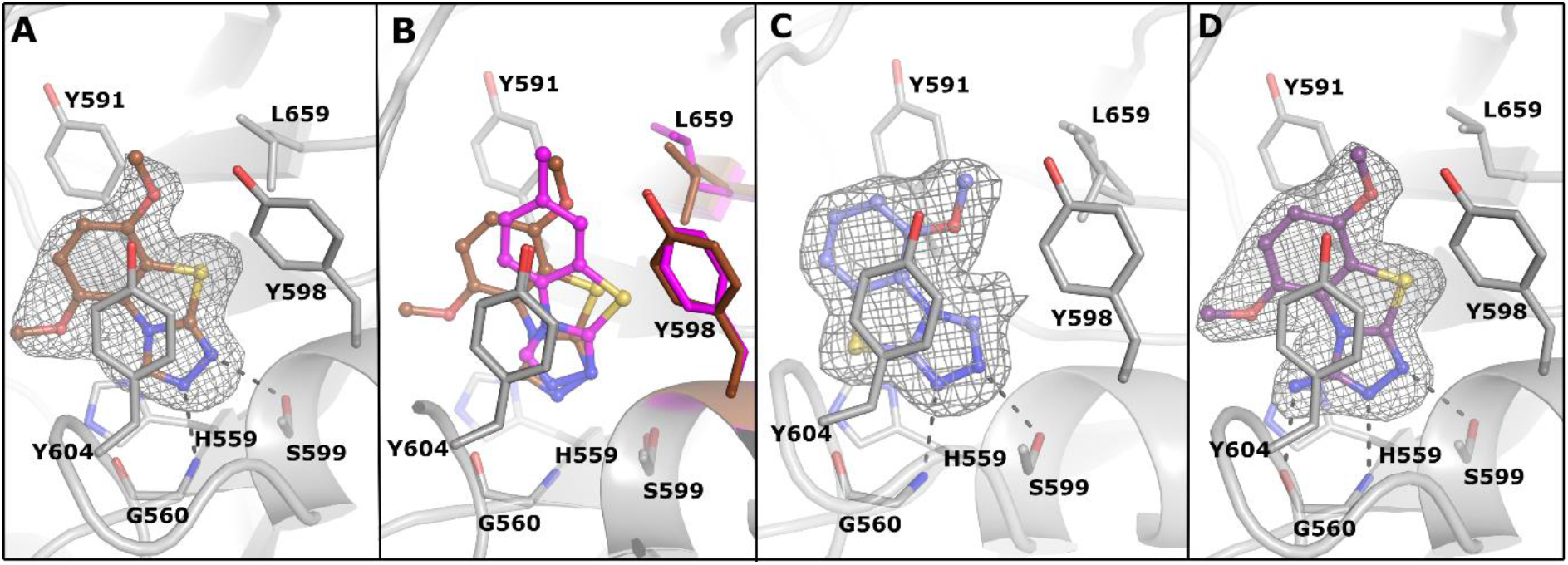
Compound rotation in the PARP active site. **(A)** PARP15 crystal structure in complex with **14**. The ligand is presented as a ball-and-stick model and colored in brown. **(B)** Superimposition of the PARP15 complex structures of **1** (magenta) and **14** (brown). Residues having conformational changes are colored with respective colors regarding the ligands. **(C)** PARP15 crystal structure in complex with **13** and **(D)** with **27**. The ligands are presented as a ball-and-stick model and colored in light blue and purple, respectively. The hydrogen bonds are indicated with black dashes. The ligand-omitted sigma A weighted Fo-Fc electron density maps are colored in grey and contoured at 3.0 σ.

We hypothesized that the plasticity would allow compounds such as **13** and **14** to inhibit multiple PARPs as the compound activities were still at a micromolar level against PARP2 and TNKS2 (Table 2). Therefore, we decided to add an anchor point to the C-3 position in order to fix the compound orientation in the binding pocket. A similar strategy was previously successfully used in the development of TNKS inhibitors.^45^ We tested multiple substituents at the C-3 position while preserving the C-7 methyl of compound **1** or the 5,8-dimethoxy of compound **14**. Compounds having an oxygen (**19**) or small sulfur groups (**20** and **22**) integrated to the scaffold **1** emerged as selective against PARP10 with micromolar activities. A more interesting compound was achieved when using an amino group as C-3 substituent that resulted in **21** showing nanomolar activity against PARP10 (IC_50_ = 180 nM) and PARP15 (IC_50_ = 300 nM) with a clear selectivity towards the mono-ARTs over the poly-ARTs PARP2 and TNKS2, which were inhibited only with micromolar potencies (IC_50_ = 1.6 µM and 5.7 µM, respectively).

To potentially improve the selectivity, we extended the thiol and amino groups with a longer substituent but, independently of the heteroatom, this caused a loss of activity (**23** – **25)** However, the imine derivative **24**, showed some activity against PARP2 (IC_50_ = 2.9 µM) and PARP15 (IC_50_ = 7.9 µM) when compared to more flexible compound **25** (Table 3). Regarding the C-3 substituted dimethoxy analogs, the presence of a thiol group determined a loss of activity for compound **26**, while **28** having a thiomethyl group recovered a modest selectivity against PARP2 (IC_50_ = 8.4 µM). The presence of a 3-amino group emerged as particularly suitable to improve the potency against PARP10 and PARP15,with IC_50_ ranging from 120 to 300 nM, with dimethoxy derivative **27** that also stood out as selective for MARylating enzymes (Table 3). The PARP15 crystal structure in complex with **27** (Figure 4D) showed a highly similar binding mode to the analogue **14** (Figure 3A). However, the amino group of **27** creates a hydrogen bond with Gly560 which together with the activity profiles (**1** vs. **21** and **14** vs. **27**) indicate that the anchor in C-3 is crucial for gaining selectivity against mono-ARTs. To improve even more the selectivity, we extended the anchor, this time by preparing the amide derivatives **29-31**, but in the presence of both either longer or shorter substituents only very weak activity was observed against some enzymes (Table 3).

### PARP profiling and biological evaluation

The 7-hydroxy derivative **16** (**OUL-245**), and the 3-amino derivatives **21** (**OUL-243**) and **27** (**OUL-232**) emerged as the most interesting compounds of the work as they were both potent and showed selectivity towards either mono- or poly-ARTs. We therefore decided to profile them against a large panel of enzymatically active PARPs (Table 4).

**Table 4.**
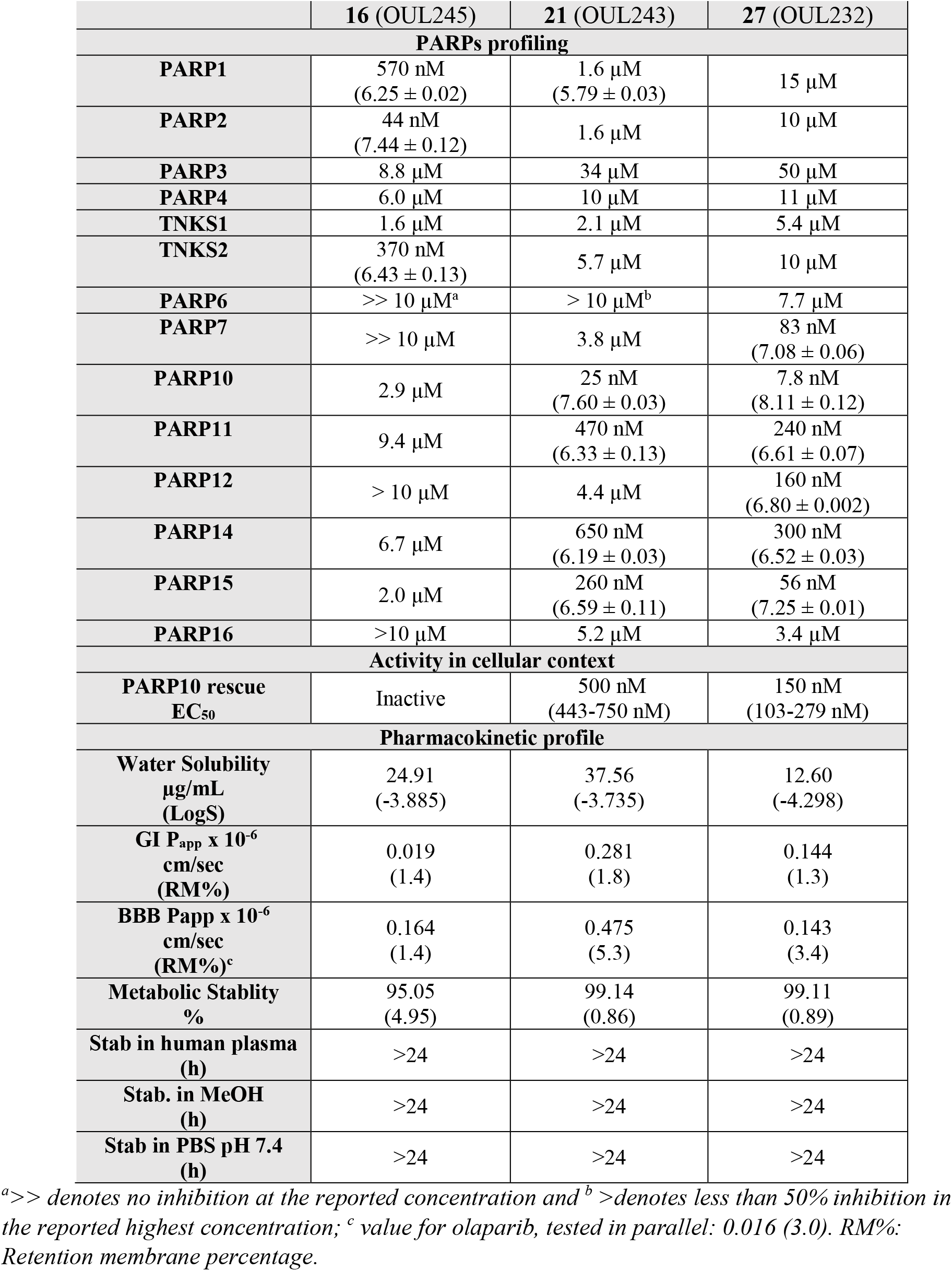
Profile of the selected compounds against the PARP enzymes, IC_50_ (pIC_50_±SEM, n=3), where mono-ARTs PARP7-PARP16 are measured using a proximity enhanced assay,^46^ potency of the compounds in rescuing cells from PARP10 overexpression along with 95% confidence interval for the EC_50_, and ADME profiling

|  | 16 (OUL245) | 21 (OUL243) | 27 (OUL232) |
| --- | --- | --- | --- |
| **PARPs profiling** |  |  |  |
| **PARP1** | 570 nM<br>(6.25 ± 0.02) | 1.6 μM<br>(5.79 ± 0.03) | 15 μM |
| **PARP2** | 44 nM<br>(7.44 ± 0.12) | 1.6 μM | 10 μM |
| **PARP3** | 8.8 μM | 34 μM | 50 μM |
| **PARP4** | 6.0 μM | 10 μM | 11 μM |
| **TNKS1** | 1.6 μM | 2.1 μM | 5.4 μM |
| **TNKS2** | 370 nM<br>(6.43 ± 0.13) | 5.7 μM | 10 μM |
| **PARP6** | >> 10 μM<sup>a</sup> | > 10 μM<sup>b</sup> | 7.7 μM |
| **PARP7** | >> 10 μM | 3.8 μM | 83 nM<br>(7.08 ± 0.06) |
| **PARP10** | 2.9 μM | 25 nM<br>(7.60 ± 0.03) | 7.8 nM<br>(8.11 ± 0.12) |
| **PARP11** | 9.4 μM | 470 nM<br>(6.33 ± 0.13) | 240 nM<br>(6.61 ± 0.07) |
| **PARP12** | > 10 μM | 4.4 μM | 160 nM<br>(6.80 ± 0.002) |
| **PARP14** | 6.7 μM | 650 nM<br>(6.19 ± 0.03) | 300 nM<br>(6.52 ± 0.03) |
| **PARP15** | 2.0 μM | 260 nM<br>(6.59 ± 0.11) | 56 nM<br>(7.25 ± 0.01) |
| **PARP16** | >10 μM | 5.2 μM | 3.4 μM |
| **Activity in cellular context** |  |  |  |
| **PARP10 rescue**<br>**EC<sub>50</sub>** | Inactive | 500 nM<br>(443-750 nM) | 150 nM<br>(103-279 nM) |
| **Pharmacokinetic profile** |  |  |  |
| **Water Solubility**<br>**μg/mL**<br>**(LogS)** | 24.91<br>(-3.885) | 37.56<br>(-3.735) | 12.60<br>(-4.298) |
| **GI P<sub>app</sub> x 10<sup>-6</sup>**<br>**cm/sec**<br>**(RM%)** | 0.019<br>(1.4) | 0.281<br>(1.8) | 0.144<br>(1.3) |
| **BBB Papp x 10<sup>-6</sup>**<br>**cm/sec**<br>**(RM%)<sup>c</sup>** | 0.164<br>(1.4) | 0.475<br>(5.3) | 0.143<br>(3.4) |
| **Metabolic Stability**<br>**%** | 95.05<br>(4.95) | 99.14<br>(0.86) | 99.11<br>(0.89) |
| **Stab in human plasma**<br>**(h)** | >24 | >24 | >24 |
| **Stab. in MeOH**<br>**(h)** | >24 | >24 | >24 |
| **Stab in PBS pH 7.4**<br>**(h)** | >24 | >24 | >24 |

<sup>a</sup>>> denotes no inhibition at the reported concentration and <sup>b</sup>>denotes less than 50% inhibition in the reported highest concentration; <sup>c</sup> value for olaparib, tested in parallel: 0.016 (3.0). RM%: Retention membrane percentage.

It should be noted that compounds **21** and **27** had reached the sensitivity limit of the mono-ART assay and that the values reported in Table 3 were artificially high due to the enzyme concentrations needed for a robust conversion of NAD^+^. We therefore had to improve the assay method and developed a homogeneous proximity enhanced assay for mono-ARTs.^46^ This new assay was used here to test the selected compounds against PARP7-PARP16 as it allowed us to measure robust IC_50_ values for the discovered potent inhibitors while using less enzyme (Figure S3).

As emerged from Table 4, both the 3-amino derivatives **21** and **27** were potent inhibitors of multiple human mono-ARTs. While **21** still inhibited multiple poly-ARTs at low µM concentration, **27** was overall more selective for mono-ARTs in agreement with our initial assessment. Compound **27** showed the highest potency as it inhibited PARP10 with an IC_50_ of 7.8 nM making it the best PARP10 inhibitor described to date. Additionally, **27** inhibited PARP7, PARP11, PARP12, PARP14 and PARP15 at low nanomolar potencies. Notably, no inhibitors of PARP12 have been described earlier, and just a few PARP15 inhibitors have been discovered recently.^35,43^

Compound **16** was confirmed as a weak inhibitor of mono-ARTs while showing potent inhibition of poly-ARTs PARP1-2 and TNKS2. Interestingly, compound **16** shows selectivity towards PARP2 (IC_50_ = 44 nM) even over highly similar PARP1 (13-fold), the same behavior was observed when comparing its activity on TNKS2 that was 4-fold higher than on TNKS1. In addition, **16** showed µM IC_50_ values for the other active site glutamate containing mono-ARTs PARP3-4. This is consistent with the crystal structures where the hydroxyl of **16** forms a hydrogen bond with the glutamate (Figure 3B).

### Cell assay for PARP10 target engagement

Taken together, our *in vitro* experiments identified a potent scaffold that when suitably functionalized gave derivatives that inhibit multiple PARPs. To complement and strengthen our data, we aimed at demonstrating the effectiveness of these compounds in a cell model. We tested **16, 21** and **27** for their capability of rescuing cells from PARP10-induced cell death using a colony formation assay. In line with our results of the WST-1 assay, none of the compounds showed toxicity in this assay using control cells expressing a catalytically inactive PARP10 mutant (PARP0-GW) (Figure 5A).

**Figure 5.**
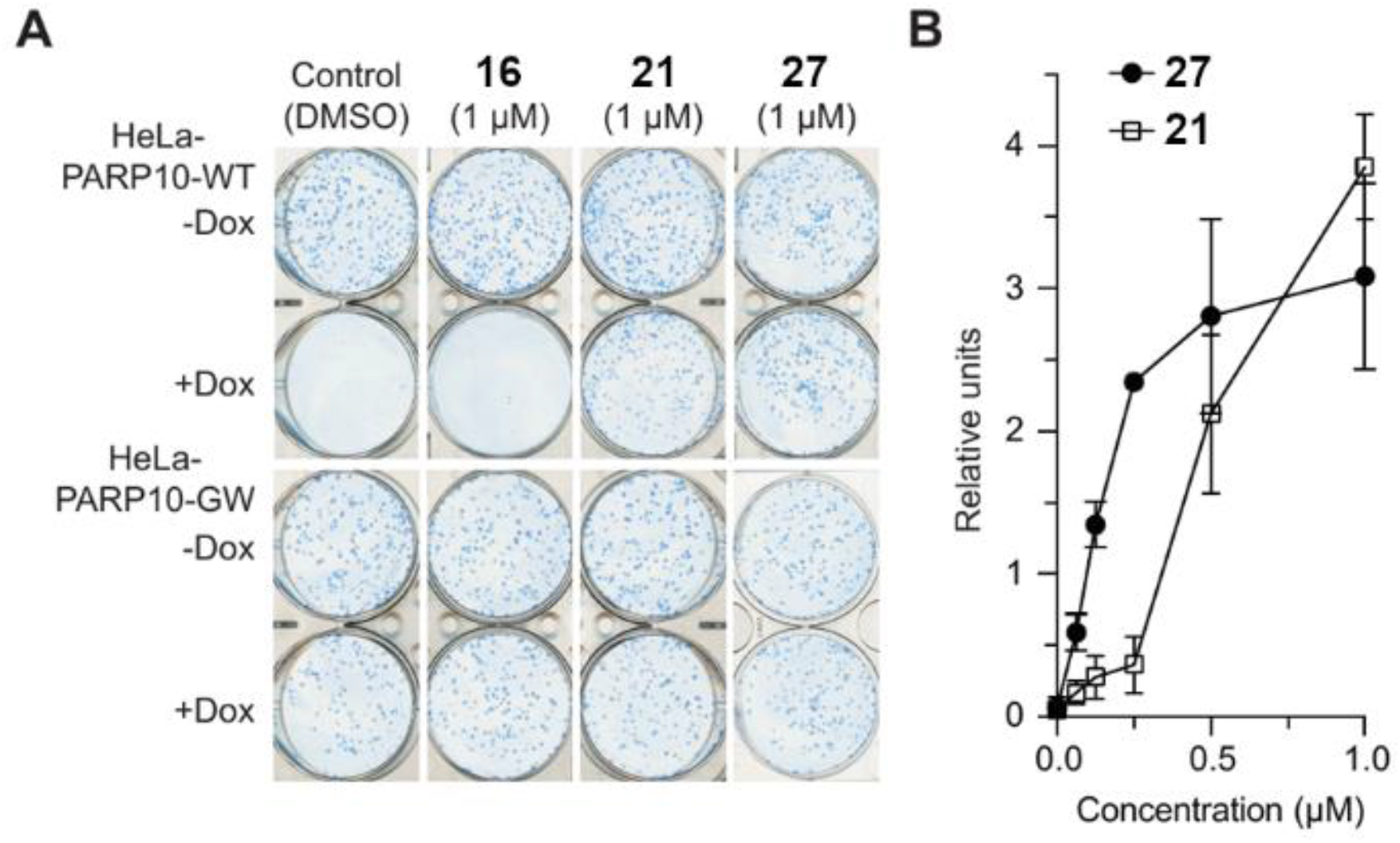
Cell assay for PARP10 inhibition. **(A)** Compounds **21** and **27** effectively rescue the PARP10 overexpressing cells from ADP-ribosylation dependent cell death whereas **16** does not show this effect consistent with its lower potency. The compounds are not toxic (-Dox) and do not affect the cells expressing catalytically inactive PARP-10-GW mutant. Cell colonies were grown for 10-12 days, stained with methylene blue. **(B)** Quantifications of titration experiments that were measured using ImageJ. Mean and standard deviation for 3 experiments are shown.

When wild type PARP10 overexpression is induced by doxycyclin (Dox) it leads to cell death observed from the lack of colonies (Figure 5A). At 1 µM, compounds **21** and **27** efficiently rescued cells from the PARP10-induced cell death, while **16** did not show any effect at this concentration (Figure 5A). The results are in good agreement with the enzymatic IC_50_ values (Table 4) and demonstrate the usability of the compounds based on TBT scaffold in inhibiting PARPs in cellular contexts. The titration experiments with **21** and **27** revealed that they are indeed the most potent PARP10 inhibitors described so far also in cell assays (Figure 5B). Especially **27** was effective in rescuing the cells with an EC_50_ of 150 nM (Table 4), 4-fold more potent than the compounds disclosed earlier,^35^ and again the ranking is in good agreement with the potencies measured in enzymatic assays.

### *In vitro* ADME properties

The colony forming assay had already shown the ability of the TBT compounds to enter the cells where they engaged the target enzymes, but the successive preclinical studies as well as their use in proof-of-concept *in vivo* studies, require a wider physicochemical characterization. Thus, the *in vitro* ADME properties were assessed for the best PARP inhibitors, **16, 21**, and **27** (Table 4). First, we evaluated the thermodynamic water solubility and from the results compounds **16** and **21** present a good solubility of 24.91 µg/mL and 37.56 µg/mL, respectively. This is likely due to the presence of polar -OH and –NH_2_ groups. The two -OMe groups instead imparted lower solubility to compound **27** (12.60 µg/mL) (Table 4).

Then, parallel artificial membrane permeability assay (PAMPA) was performed to predict passive permeability through different biological membranes, such as the gastrointestinal tract (GI) and the blood brain barrier (BBB). The results shown in Table 4 suggest that all three compounds exhibit suboptimal GI permeability and low membrane retention, reflecting their degree of hydrophilicity already observed in the aqueous solubility tests. The permeability was better in BBB for smaller compounds **16** (Papp 0.164) and **21** (Papp 0.475) which is 10-30 times higher than that of a clinical PARP inhibitor olaparib tested in parallel (Papp 0.016). Although improved over the control inhibitor, the permeability should be taken into account in the successive optimization steps.^47^

All the compounds showed excellent phase I metabolic stability in human liver microsomes, with compounds **21** and **27** showing >99% of unchanged compound, while **16** exhibited a lower although good stability (>95%). The lower steric bulk on the molecule with only a hydroxyl substituent could have enabled the formation of a metabolite likely due to aromatic oxidation (4.95%). Finally, stability tests were performed in MeOH, PBS buffer and human plasma incubated at 37 °C. All compounds were shown to be stable in these conditions for more than 24 h.

## DISCUSSION AND CONCLUSIONS

Most of the PARP inhibitors, whether inhibiting mono- or poly-ARTs, share a benzamide group as nicotinamide mimic moiety, or such a group rigidified into a cycle.^44,48^ A few other scaffolds are used as benzamide bioisosters of which the triazole is also exploited.^49^ Unfortunately, sharing the same pharmacophoric requirements, the compounds usually lack the desired selectivity profile.^44,50^

In this paper, we initially discovered compound **1**, based on a new nicotinamide mimic scaffold, TBT, which was able to inhibit multiple PARP enzymes at micromolar potency. The TBT is an underexplored scaffold in medicinal chemistry, with few examples of compounds with antifungal,^51^ anticonvulsant^52^ or anti-inflammatory^53^ properties. Most importantly, no PARPi based on this scaffold have been reported until now. By making small changes around the tricyclic scaffold of **1**, we were able to shift the activity from pan to a selective inhibition of either the mono- or the poly-ARTs.

In particular, the benzene ring was decorated with various alkyl substituents, as well as with one or two methoxy or hydroxy groups, or with halogen atoms, while the C-3 position of the triazole ring was functionalized with heteroatoms also derivatized with side chains of different length. The amino group emerged as the most interesting C-3 substituent that when coupled with a 7-methyl or 5,8-dimethoxy groups on the benzene ring, gave compounds **21** and **27**, respectively which had nM potency and selectivity toward mono-ARTs. Of note, compound **27** emerged as the most potent PARP10 inhibitor ever reported to date both in the enzymatic assay (IC_50_ = 7.8 nM) and in inhibition of intracellular PARP10 (EC_50_ = 150 nM). Furthermore, it also potently inhibited PARP15 with IC_50_ of 56 nM and PARP12 at 160 nM becoming the first potent PARP12 inhibitor. 7-Hydroxy derivative **16** is of special interest as a poly-ART inhibitor since it is both potent (IC_50_ = 44 nM) and specific PARP2 inhibitor with 13-fold selectivity over PARP1. Structurally the PARP1 and PARP2 are both able to form the same interactions with the hydroxy moiety (Figure 3A) and therefore there is no apparent reason for the observed selectivity.

Preliminary ADME analysis indicates good aqueous solubility, not limiting passive permeability in GI and BBB, and excellent stability. The effectiveness of the currently approved PARP1 inhibitors has been shown to be significantly reduced by their poor brain availability due to efflux transporters and restricted delivery across BBB. Comparison studies have shown that niraparib has a greater tumor exposure and sustainability in the brain, while olaparib, rucaparib and talazoparib have a more limited BBB penetration.^54–56^ Limited data is available to understand the penetration and residence of PARP inhibitors in a disrupted BBB setting but pharmacokinetic studies have shown that olaparib, despite the low permeability in PAMPA model, is able to penetrate recurrent glioblastoma at levels allowing radiosensitization.^57^

There is still an unmet need to discover PARPi with the appropriate profile to treat brain metastasis, brain cancers or to potentially treat neurodegenerative diseases^58^ and the TBT scaffold may thus be a potential candidate for further development towards these indications. In summary, the nM potencies measured for the TBT analogs both for mono- and poly-ARTs, experimentally determined binding modes, and favorable ADME properties elucidate possibilities on development of PARP specific chemical probes and drug leads based on the compounds disclosed here.

## EXPERIMENTAL SECTION

### Chemistry

All starting materials, reagents, and solvents were purchased from common commercial suppliers and were used as such without further purification. Compounds **2**^59^, **7**^60^, **11**^52^, **16**^52^, **19**^61^, **20**^53^, **33**^62^, **44**^63^, **45**^64^, **46**^65^, **48**^63^, **49**^66^, **50**^67^, **51**^68^, **52**^69^, **53**^68^, **54**^70^, **55**^71^, **56**^71^, **58**^70^, **59**^70^, **60**^72^, **61**^73^, **62**^69^, **63**^70^, **64**^74^, **68**^74^, **69**^74^, **70**^74^, **73**^74^, **77-79**^75^, **80**^76^, **81**^77^, and **82**^69^ were prepared as described in literature. Organic solutions were dried over anhydrous Na_2_SO_4_ and concentrated with rotary evaporator at low pressure. All the reactions were routinely checked by thin-layer chromatography (TLC) on silica gel 60F254 (Merck) and visualized by using UV and iodine. Flash chromatography separations were carried out on Merck silica gel 60 (mesh 230-400) or by using automated Buchi Reveleris X2-UV with column FP Ecoflex Si 12 g. Yields were of purified products and were not optimized. ^1^HNMR spectra were recorded at 400 MHz (Bruker Avance DRX-400), while ^13^CNMR spectra were recorded at 101 MHz (Bruker Avance DRX-400). Chemical shifts are given in ppm (*δ*) relative to TMS. Spectra were acquired at 298 K. Data processing was performed with standard Bruker software XwinNMR, and the spectral data are consistent with the assigned structures. Mating constant (*J*) are reported in Hz. The purity of the tested compounds was evaluated by HPLC analysis using Jasco LC-4000 instrument equipped with a UV-Visible Diode Array Jasco-MD4015 (Jasco Corporation, Tokyo, Japan), and XTerra MS C18 column, 5 µm x 4.6 mm x 150 mm (Waters Corporation, Massachusetts, USA). Chromatograms were analyzed by ChromNAV2.0 Chromatography Data System software. The purity of the compounds, performed at λ 254 nm, at the λ max of each compound and the absolute maximum of absorbance between 200 and 600 nm, was ≥ 95 %. The peak retention time (ret. time) is given in minutes. High resolution mass detection was performed for some representative compounds and it was based on electrospray ionization (ESI) in positive polarity using Agilent 1290 Infinity System equipped with a MS detector Agilent 6540A Accurate Mass Q-TOF.

### 4 Fluoro-7-methyl-1,3-benzothiazol-2-amine (57)

A solution of Br_2_ (0.2 mL, 3.3 mmol) in CHCl_3_ (5 mL) was slowly added to a suspension of **47** (0.6 g, 3.3 mmol) in CHCl_3_ (13 mL), at 0 °C. The reaction mixture was stirred at r.t. for 4 h and, then, a solution of 10 % Na_2_SO_3_ was added to the mixture. CHCl_3_ was removed under reduced pressure and NH_4_OH solution was added until the formation of a precipitate that was filtered, giving compound **57** (0.36 g, 66%). ^1^H NMR (400 MHz, DMSO-*d*_6_) *δ*: 2.46 (3H, s, CH_3_), 6.75-6.78 (1H, m, aromatic CH), 6.92-7.00 (1H, m, aromatic CH), 7.67 (2H, bs, NH_2_).

### 6-Ethyl-2-hydrazino-1,3-benzothiazole (65). General procedure (A) for the synthesis of hydrazinobenzothiazoles)

Hydrazine hydrate (0.30 mL, 5.88 mmol) and CH_3_COOH (0.17 mL, 2.94 mmol) were added to a suspension of **55**^71^ (0.35 g, 1.96 mmol) in ethylene glycol (18 mL), and the reaction mixture was stirred for 12 h at 125 °C. Then, the mixture was then poured into ice/water and a saturated solution of NaHCO_3_ was added until pH = 8, to give a precipitate that was filtered yielding **65** (0.158 g, 42%). ^1^H NMR (400 MHz, DMSO-*d*_6_) *δ*: 1.19 (3H, t, *J* =7.4 Hz, CH_2_*CH*_*3*_), 2.62 (2H, q, *J* = 7.4 Hz, *CH*_*2*_CH_3_), 4.96 (2H, bs, NH_2_), 7.04 (1H, d, *J* = 7.6 Hz, H4), 7.23 (1H, d, *J* = 7.9 Hz, H5), 7.51 (1H, s, H7), 8.86 (1H, bs, NH).

### 6-Ispropyl-2-hydrazino-1,3-benzothiazole (66)

The title compound was prepared according to the general procedure A, starting from **56**^71^ (30 h) in 52 % yield. ^1^H NMR (400 MHz, DMSO-*d*_6_) *δ*: 1.21 (3H, d, *J* =2.1 Hz, CH_3_), 1.22 (3H, d, *J* =2.1 Hz, CH_3_), 2.87-2.91 (1H, m, CH), 4.97 (2H, bs, NH_2_), 7.08 (1H, d, *J* = 8.3 Hz, H4), 7.23 (1H, dd, *J* = 1.9 and 8.2 Hz, H5), 7.55 (1H, s, H7), 8.88 (1H, bs, NH).

### 4-Fluoro-2-hydrazino-7-methyl-1,3-benzothiazole (67)

The title compound was prepared according to the general procedure A, starting from **57** (24 h) in 32% yield. ^1^H NMR (400 MHz, DMSO-*d*_6_) *δ*: 2.26 (3H, s, CH_3_), 5.11 (2H, bs, NH_2_), 6.72-6.76 (1H, m, aromatic H), 6.91-6.96 (1H, m, aromatic H), 9.09 (1H, bs, NH).

### 2-Hydrazino-4,7-dimethoxy-1,3-benzothiazole (71)

The title compound was prepared according to the general procedure A, starting from **61**^73^ (26 h) in 20 % yield. ^1^H NMR (400 MHz, DMSO-*d*_6_) *δ*: 3.73 (3H, s, OCH_3_), 3.77 (3H, s, OCH_3_), 5.00 (2H, bs, NH_2_), 6.48 (1H, d, *J* = 8.6 Hz, H5), 6.70 (1H, d, *J* = 8.6 Hz, H6), 8.86 (1H, bs, NH).

### 2-Hydrazino-5,7-dimethoxy-1,3-benzothiazole (72)

The title compound was prepared according to the general procedure A, starting from **62**^69^ (20 h) in 44% yield. ^1^H NMR (400 MHz, DMSO-*d*_6_) *δ*: 3.75 (3H, s, OCH_3_), 3.87 (3H, s, OCH_3_), 5.00 (2H, bs, NH_2_), 6.27 (1H, s, H5), 6.56 (1H, s, H6), 8.98 (1H, bs, NH).

### 6 Methyl[1,2,4]triazolo[3,4-*b*][1,3]benzothiazole (3). General procedure (B) for the synthesis of [1,2,4]triazolo[3,4-*b*][1,3]benzothiazoles

A solution of **64**^74^ (0.14 g, 0.78 mmol) in formic acid (5 mL) was refluxed for 9 h. The reaction mixture was then poured in ice/water and the pH was neutralized using a saturated solution of NaHCO_3_. The reaction mixture was extracted with EtOAc (x3) and the organic layers were washed with brine, dried over Na_2_SO_4_ and evaporated to dryness under reduced pressure to give a solid that was purified by crystallization using cyclohexane/EtOAc (2:1) yielding **3** (0.010 g, 10%). ^1^H NMR (400 MHz, DMSO-*d*_6_) *δ*: 2.41 (3H, s, CH_3_), 7.28 (1H, d, *J* = 8.2 Hz, H7), 7.87 (1H, d, *J* = 8.2 Hz, H8), 92 (1H, s, H5), 9.55 (1H, s, H3). ^13^C NMR (101 MHz, DMSO-*d*_6_) *δ*: 21.34, 115.50, 125.48, 127.99, 128.67, 129.35, 137.00, 137.41, 155.12. HPLC: CH_3_CN/H_2_O + 0.1% FA (70:30), ret. time: 2.05 min, peak area: 99.21%.

### 7 Methyl[1,2,4]triazolo[3,4-*b*][1,3]benzothiazole (4)

The title compound was prepared according to the general procedure B starting from **80**^76^ (12 h) in 30% yield as a pink solid, after purification by flash chromatography eluting with CHCl_3_:MeOH (95:5). ^1^H NMR (400 MHz, DMSO-*d*_6_) *δ*: 2.42 (3H, s, CH_3_), 7.27-7.31 (1H, m, H7), 7.48 (1H, t, *J* = 7.7 Hz, H6), 7.91-7.93 (1H, m, H5), 9.65 (1H, s, H3). ^13^C NMR (101 MHz, DMSO-*d*_6_) *δ*: 19.70, 112.85, 127.55, 127.61, 129.18, 131.54, 134.71, 137.44, 154.07. HPLC: CH_3_CN/H_2_O + 0.1% FA (70:30), ret. time: 2.06 min, peak area: 99.9%.

### 7-Ethyl[1,2,4]triazolo[3,4-*b*][1,3]benzothiazole (5)

The title compound was prepared according to the general procedure B starting from **65** (24 h) in 50% yield as a yellow solid, after purification by flash chromatography eluting with CHCl_3_:MeOH (99:1). ^1^H NMR (400 MHz, DMSO-*d*_6_) *δ*: 1.23 (3H, t, *J* = 7.5 Hz, CH_2_*CH*_*3*_), 2.73 (2H, q, *J* = 7.6 Hz, *CH*_*2*_CH_3_), 7.44 (1H, dd, *J* = 1.0 and 7.3 Hz, H6), 7.89 (1H, s, H8), 8.02 (1H, d, *J* = 8.3 Hz, H5), 9.60 (1H, s, H3). ^13^C NMR (101 MHz, DMSO-*d*_6_) *δ*: 16.18, 28.61, 115.13, 124.77, 127.33, 127.62, 132.13, 137.18, 143.33, 154.88. HPLC: CH_3_CN/H_2_O (65:35), ret. time: 2.81 min, peak area: 98.49%.

### 7-Isopropyl[1,2,4]triazolo[3,4-*b*][1,3]benzothiazole (6)

The title compound was prepared according to the general procedure B starting from **66** (24 h) in 58% yield as a white solid, after purification by flash chromatography eluting with CHCl_3_:MeOH (99:1) and successive treatment with cyclohexane. ^1^H NMR (400 MHz, DMSO-*d*_6_) *δ*: 1.26 (6H, d, *J* = 6.9 Hz, CH_3_ x 2), 3.01-3.04 (1H, m, CH), 7.47-7.49 (1H, m, H6), 7.95 (1H, s, H8), 8.03 (1H, d, *J* = 8.3 Hz, H5), 9.60 (1H, s, H3). ^13^C NMR (101 MHz, DMSO-*d*_6_) *δ*: 24.44, 34.07, 115.15, 123.44, 126.05, 127.70, 132.16, 137.19, 147.99, 154.93. HPLC: CH_3_CN/H_2_O (65:35), ret. time: 3.50 min, peak area: 98.13%.

### 5-Fluoro-8-methyl[1,2,4]triazolo[3,4-*b*][1,3]benzothiazole (9)

The title compound was prepared according to the general procedure B starting from **67** (10 h) in 13% yield as a yellowish, after purification by flash column chromatography eluting with CH_2_Cl_2_:MeOH (98:2). ^1^H NMR (400 MHz, DMSO-*d*_6_) *δ*: 2.37 (3H, s, CH_3_), 7.32 (1H, m, H7), 7.43 (1H, t, *J* = 8.3 Hz, H6), 9.43 (1H, s, H3). ^13^C NMR (101 MHz, DMSO-*d*_6_) *δ*: 18.91, 114.06 (d, *J* = 16.5 Hz), 117.63 (d, *J* = 16.2 Hz), 127.89 (d, *J* = 6.5 Hz), 130.21 (d, *J* = 4.0 Hz), 133.54 (d, *J* = 2.5 Hz), 138.15, 148.63 (d, *J* = 245 Hz), 154.06. HPLC: CH_3_CN/H_2_O + 0.1% FA (70:30), ret. time: 2.12 min, peak area: 99.9%.

### 6-Methoxy[1,2,4]triazolo[3,4-*b*][1,3]benzothiazole (10)

The title compound was prepared according to general procedure B starting from **68**^74^ (24 h) in 15% yield as a white solid, after purification by flash chromatography eluting with CH_2_Cl_2_:MeOH (98:2). ^1^H NMR (400 MHz, DMSO-*d*_6_) *δ*: 3.93 (3H, s, OCH_3_), 8.01 (2H, s, aromatic H), 8.30 (1H, s, aromatic H), 9.56 (1H, s, H3). ^13^C NMR (101 MHz, DMSO-*d*_6_) *δ*: 57.45, 100.22, 108.92, 123.75, 129.21, 129.73, 137.09, 155.21, 155.82. HPLC: CH_3_CN/H_2_O + 0.1% FA (70:30), ret. time: 2.19 min, peak area: 99.72%.

### 8-Methoxy[1,2,4]triazolo[3,4-*b*][1,3]benzothiazole (12)

The title compound was prepared according to the general procedure B starting from **82**^69^ (6 h) in 31% yield as a pink solid, after purification by flash chromatography eluting with CH_2_Cl_2_:MeOH (98:2). ^1^H NMR (400 MHz, DMSO-*d*_6_) *δ*: 3.95 (3H, s, OCH_3_), 7.12 (1H, d, *J* = 8.2 Hz, H7), 7.49-7.55 (1H, m, H6), 7.67 (1H, d, *J* = 8.2 Hz, H5). ^13^C NMR (101 MHz, DMSO-*d*_6_) *δ*: 56.87, 107.90, 108.81, 118.83, 129.02, 130.20, 137.42, 154.79. HPLC: CH_3_CN/H_2_O + 0.1% FA (70:30), ret. time: 1.70 min, peak area: 98.77%.

### 5-Methoxy[1,2,4]triazolo[3,4-*b*][1,3]benzothiazole (13)

The title compound was prepared according to the general procedure B starting from **70**^74^ (6 h) in 23% yield as a yellow solid, after purification by flash chromatography eluting with CH_2_Cl_2_:MeOH (98:2). ^1^H NMR (400 MHz, DMSO-*d*_6_) *δ*: 4.01 (1H, s, OCH_3_), 7.22 (1H, d, *J* = 6.7 Hz, H6), 7.42 (1H, t, *J* = 8.1 Hz, H7), 7.55 (1H, d, *J* = 8.2 Hz, H8), 9.34 (1H, s, H3). ^13^C NMR (101 MHz, DMSO-*d*_6_) *δ*: 56.98, 109.94, 117.20, 119.23, 127.71, 132.88, 138.28, 148.28, 154.57. HPLC: CH_3_CN/H_2_O + 0.1 % FA (70:30), ret. time: 1.70 min, peak area: 99.47%.

### 5,8-Dimethoxy[1,2,4]triazolo[3,4-*b*][1,3]benzothiazole (14)

The title compound was prepared according to the general procedure B starting from **71** (12 h) in 25% yield as a pink solid, after purification by crystallization using cyclohexane/EtOAc (2:1). ^1^H NMR (400 MHz, DMSO-*d*_6_) *δ*: 3.89 (3H, s, OCH_3_), 3.95 (3H, s, OCH_3_), 7.04 (1H, d, *J* = 9.0 Hz, H6), 7.15 (1H, d, *J* = 9.0 Hz, H7), 9.29 (1H, s, H3). ^13^C NMR (101 MHz, DMSO-*d*_6_) *δ*: 56.85, 57.12, 108.40, 110.55, 119.58, 119.97, 138.20, 142.52, 148.15, 154.51. HRMS: *m/z* calcd for C_27_H_28_N_3_O_4_S 258.0313 [M + Na^+^], found 258.0310. HPLC: CH_3_CN/H_2_O + 0.1% FA (70:30), ret. time: 2.15 min, peak area: 99.53%.

### 6,8-Dimethoxy[1,2,4]triazolo[3,4-*b*][1,3]benzothiazole (15)

The title compound was prepared according to the general procedure B starting from **72** (15 h) in 10% yield as a white solid, after purification by flash chromatography eluting with CH_2_Cl_2_:MeOH (98:2). ^1^H NMR (400 MHz, DMSO-*d*_6_) *δ*: 3.88 (1H, s, OCH_3_), 3.96 (1H, s, OCH_3_), 6.76 (1H, s, H7), 7.46 (1H, s, H5), 9.58 (1H, s, H3). ^13^C NMR (101 MHz, DMSO-*d*_6_) *δ*: 56.72, 57.07, 93.28, 97.86, 110.21, 130.48, 137.27, 155.35, 155.42, 161.11. HPLC: CH_3_CN/H_2_O (70:30), ret. time: 1.78 min, peak area: 97.88%.

### [1,2,4]Triazolo[3,4-*b*][1,3]benzothiazol-8-ol (17)

A 1M solution of BBr_3_ in CH_2_Cl_2_ (1.22 mL, 1.22 mmol) was added dropwise to a solution of **12** (0.05 g, 0.24 mmol) in dry CH_2_Cl_2_ (3 mL) at 0 °C and under nitrogen atmosphere. The reaction was stirred at rt overnight and, then, MeOH was added. The mixture was evaporated under reduced pressure to give a residue that was poured into ice/water, added of 6N HCl until pH 4, and extracted with EtOAc. The organic layers were washed with brine, dried over Na_2_SO_4_ and evaporated under reduced pressure, obtaining a solid that was purified by flash chromatography eluting with CH_2_Cl_2_:MeOH (95:5), yielding **17** as a brown solid (22%). ^1^H NMR (400 MHz, DMSO-*d*_6_) *δ*: 6.96 (1H, d, *J* = 8.2 Hz, H7), 7.43 (1H, t, *J* = 8.1 Hz, H6), 7.59 (1H, d, *J* = 8.1 Hz, H5), 9.62 (1H, s, H3), 11.14 (1H, s, OH). ^13^C NMR (101 MHz, DMSO-*d*_6_) *δ*: 106.30, 112.68, 117.96, 128.84, 130.64, 137.40, 153.73, 154.89. HPLC: CH_3_CN + 0.1% FA/H_2_O + 0.1% FA (50:50), ret. time: 2.00 min, peak area: 99.12%.

### [1,2,4]Triazolo[3,4-*b*][1,3]benzothiazol-5,8-diol (18)

The title compound was synthesized following the same procedure as used for the synthesis of compound 17 starting from **14** in 10% yield as a purple solid, after purification with flash column chromatography eluting with CH_2_Cl_2_:MeOH (95:5)^1^H NMR (400 MHz, DMSO-*d*_6_) *δ*: 6.79 (1H, d, *J* = 8.7 Hz, H7), 6.89 (1H, d, *J* = 8.8 Hz, H6), 9.24 (1H, s, H3), 10.29 (1H, s, OH), 10.43 (1H, s, OH). ^13^C NMR (101 MHz, DMSO-*d*_6_) *δ*: 112.72, 114.60, 118.87, 119.17, 138.04, 139.41, 145.59, 154.59. HPLC: CH_3_CN + 0.1% FA/H_2_O + 0.1% FA (50:50), ret. time: 1.89 min, peak area: 99.93%.

### 7-Methyl[1,2,4]triazolo[3,4-*b*][1,3]benzothiazol-3-amine (21)

CNBr (0.47 g, 4.44 mmol) was added to a solution of **73**^74^ (0.53 g, 2.96 mmol) in MeOH (10 mL) and the reaction mixture was refluxed for 3.5 h. Then, it was poured in ice/water, saturated solution of NaHCO_3_ was added until pH 8 and the obtained precipitate was filtered and purified by flash chromatography eluting with CHCl_3_:MeOH (95:5), obtaining **21** as a brown solid (0.06 g, 10%). ^1^H NMR (400 MHz, DMSO-*d*_6_) *δ*: 2.39 (3H, s, CH_3_), 6.41 (2H, bs, NH_2_), 7.30 (1H, dd, *J* = 0.8 and 8.3 Hz, H6), 7.71 (1H, s, H8), 7.90 (1H, d, *J* = 8.3 Hz, H5). ^13^C NMR (101 MHz, DMSO-*d*_6_) *δ*: 21.31, 113.76, 125.43, 127.63, 127.84, 131.43, 135.63, 149.11, 150.90. HRMS: *m/z* calcd for C_9_H_8_N_4_S 205.0550 [M + H^+^], found 205.0544. HPLC: CH_3_CN/H_2_O (70:30), ret. time: 1.61 min, peak area: 98.97%.

### 7-Methyl-3-(methylthio)[1,2,4]triazolo[3,4-*b*][1,3]benzothiazole (22)

K_2_CO_3_ (0.21 g, 1.5 mmol) and MeI (0.1 mL, 1 mmol) were added under nitrogen atmosphere to a solution of **20**^53^ (0.11 g, 0.5 mmol) in dry DMF (6 mL). The reaction mixture was stirred at 80 °C for 1.5 h and then poured in ice/water. The mixture was acidified with 2N HCl until pH 5 and the obtained precipitate was filtered and purified by crystallization using cyclohexane: EtOAc (2:1), obtaining **22** as a brown solid (0.03 g, 22%). ^1^H NMR (400 MHz, DMSO-*d*_6_) *δ*: 2.39 (3H, s, CH_3_), 2.70 (3H, s, SCH_3_), 7.37 (1H, d, *J* = 8.2 Hz, H5), 7.83 (1H, s, H8), 7.86 (1H, d, *J* = 8.3 Hz, H6). ^13^C NMR (101 MHz, DMSO-*d*_6_) *δ*: 16.08, 21.26, 113.95, 125.79, 127.62, 128.25, 131.87, 136.63, 144.35, 156.11. HPLC: CH_3_CN/H_2_O + 0.1% FA (70:30), ret. time: 2.15 min, peak area: 99.88%.

### 3-[(4-Chlorobenzyl)thio]-7-methyl[1,2,4]triazolo[3,4-*b*][1,3]benzothiazole (23)

A suspension of **20**^53^ (0.12 g, 0.54 mmol) and KOH (0.03 g, 0.54 mmol) in absolute EtOH (8 mL) was refluxed for 30 minutes under nitrogen atmosphere. Then, *p*-chlorobenzyl chloride (0.09 g, 0.54 mmol) was added and the reaction mixture was stirred at reflux. After 4 h, EtOH was removed under reduced pressure to give a residue that was added of water and extracted with EtOAc (x3). The organic layers were washed with brine, dried over Na_2_SO_4_ and evaporated to dryness under reduced pressure to give a solid that was purified by crystallization using cyclohexane: EtOAc (3:1), to give **23** as a white solid (0.05 g, 27%). ^1^H NMR (400 MHz, DMSO-*d*_6_) *δ*: 2.36 (1H, s, CH_3_), 4.39 (2H, s, Bz CH_2_), 7.22-7.31 (5H, m, aromatic H), 7.80 (1H, s, H8), 7.84 (1H, d, *J* = 8.3 Hz, H6). ^13^C NMR (101 MHz, DMSO-*d*_6_) *δ*: 21.25, 37.72, 114.10, 125.61, 127.57, 128.02, 128.73, 131.19, 131.72, 132.53, 136.43, 136.62, 142.34, 156.55. HPLC: CH_3_CN/H_2_O + 0.1 % FA (70:30), ret. time: 3.32 min, peak area: 99.52%.

### *N*-[(1Z)-(4-chlorophenyl)methylene]-7-methyl[1,2,4]triazolo[3,4-*b*][1,3]benzothiazol-3-amine (24)

*p*-TsOH (10 mg, 10 % w/w) and *p*-chlorobenzaldehyde (0.07 g, 0.5 mmol) were added to a solution of **21** (0.10 g, 0.5 mmol) in dry benzene (20 mL). The reaction mixture was stirred at reflux by using a Dean-Stark apparatus for 16 h and then poured in ice/water and saturated solution of NaHCO_3_ was added until pH 8. The mixture was extracted with CH_2_Cl_2_ (x3) and the organic layers were washed with brine, dried over Na_2_SO_4_ and evaporated to dryness under reduced pressure to give an oil that was purified by flash chromatography eluting with CH_2_Cl_2_:MeOH (95:5) and then treated with Et_2_O to give **24** as a brown solid (0.02 g, 10%). ^1^H NMR (400 MHz, DMSO-*d*_6_) *δ*: 2.51 (1H, s, CH_3_), 7.43 (1H, d, *J* = 8.3 Hz, H5), 7.68 (2H, d, *J* = 8.5 Hz, H3’ and H5’), 7.85 (1H, s, H8), 8.14 (1H, d, *J* = 8.2 Hz, H6), 8.21 (2H, d, *J* = 8.5 Hz, H2’ and H6’), 9.41 (1H, s, CH). ^13^C NMR (101 MHz, DMSO-*d*_6_) *δ*: 21.55, 115.30, 125.72, 127.98, 128.94, 129.90, 132.03, 132.13, 134.24, 136.83, 138.51, 153.43, 155.14, 163.73. HRMS: *m/z* calcd for C_16_H_11_ClN_4_S 327.04770 [M + H^+^], found 327.0472. HPLC: CH_3_CN/H_2_O + 0.1 % FA (60:40), ret. time: 5.80 min, peak area: 95.64%.

### *N*-(4-chlorobenzyl)-7-methyl[1,2,4]triazolo[3,4-*b*][1,3]benzothiazol-3-amine (25)

NaBH_4_ (0.026 g, 0.69 mmol) was added to a suspension of **24** (0.15 g, 0.46 mmol) in EtOH (10 mL), at 0 °C and under nitrogen atmosphere. The reaction mixture was stirred overnight at r.t. and, then, it was poured in ice/water, saturated solution of NaHCO_3_ was added until pH 8 furnishing a precipitate that was filtered and purified by flash chromatography eluting with CH_2_Cl_2_:MeOH (97:3), to give **25** as a green solid (0.03 g, 17%). ^1^H NMR (400 MHz, DMSO-*d*_6_) *δ*: 2.41 (3H, s, CH_3_), 4.54 (2H, d, *J* = 5.6 Hz, Bz CH_2_), 7.23 (1H, bs, NH), 7.34 (1H, d, *J* = 8.3 Hz, H5), 7.39 (2H, d, *J* = 8.3 Hz, H3’ and H5’), 7.49 (2H, d, *J* = 8.3 Hz, H2’ and H6’), 7.72 (1H, s, H8), 7.97 (1H, d, *J* = 8.3 Hz, H6). ^13^C NMR (101 MHz, DMSO-*d*_6_) *δ*: 21.13, 46.17, 113.61, 125.33, 127.45, 127.57, 128.47, 129.81, 131.24, 131.70, 135.59, 139.03, 149.76, 150.64. HPLC: CH_3_CN/H_2_O + 0.1 % FA (60:40), ret. time: 2.64 min, peak area: 99.11%.

### 5,8-Dimethoxy[1,2,4]triazolo[3,4-*b*][1,3]benzothiazole-3(2*H*)-thione (26)

KOH (0.09 g, 1.7 mmol) dissolved in few drops of H_2_O was added to a suspension of **71** (0.38 g, 1.70 mmol) in EtOH (7 mL) and, then, CS_2_ (0.5 mL, 8.5 mmol) was added. The reaction mixture was refluxed for 2 h and, then, EtOH was removed under reduced pressure and 2N HCl was added. The obtained precipitate was filtered, purified by flash chromatography eluting with cyclohexane:EtOAc (80:20) and treated with EtOH, to give **26** as a purple solid (0.08 g, 18 %). ^1^H NMR (400 MHz, DMSO-*d*_6_) *δ*: 3.85 (3H, s, OCH_3_), 3.93 (3H, s, OCH_3_), 7.13 (1H, d, *J* = 9.0 Hz, H6), 7.20 (1H, d, *J* = 9.0 Hz, H7), 13.91 (1H, s, NH). ^13^C NMR (101 MHz, DMSO-*d*_6_) *δ*: 57.03, 57.82, 109.76, 114.61, 118.81, 122.65, 143.91, 148.03, 151.75, 163.65. HPLC: CH_3_CN/H_2_O (65:35), ret. time: 1.95 min, peak area: 96.90%.

### 5,8-Dimethoxy[1,2,4]triazolo[3,4-*b*][1,3]benzothiazol-3-amine (27)

The title compound was synthesized following the same procedure as used for the synthesis of compound **21** starting from **71** in 12% yield as a pink solid, after purification by flash chromatography eluting with CHCl_3_: MeOH 95:5 and successive treatment by EtOH. ^1^H NMR (400 MHz, DMSO-*d*_6_) *δ*: 3.90 (3H, s, OCH_3_), 3.99 (3H, s, OCH_3_), 6.46 (2H, bs, NH_2_), 7.02 (1H, d, *J* = 9.0 Hz, H6), 7.16 (1H, d, *J* = 9.0 Hz, H7). ^13^C NMR (101 MHz, DMSO-*d*_6_) *δ*: 57.00, 57.78, 108.36, 111.50, 120.03, 120.15, 141.48, 148.45, 148.55, 150.67. HRMS: *m/z* calcd for C_10_H_10_N_4_O_2_S 251.0590 [M + H^+^], found 251.0584. HPLC: CH_3_CN/H_2_O (70:30), ret. time: 1.64 min, peak area: 97.78%.

### 5,8-Dimethoxy-3-(methylthio)-2,3-dihydro[1,2,4]triazolo[3,4-*b*][1,3]benzothiazole (28)

The title compound was synthesized following the same procedure as used for the synthesis of compound **22** starting from **26** in 15% yield as a pink solid, after purification by flash chromatography eluting with cyclohexane:EtOAc (from 100:0 to 50:50). ^1^H NMR (400 MHz, DMSO-d6) *δ*: 2.61 (3H, s, SCH_3_), 3.87 (3H, s, OCH_3_), 3.89 (3H, s, OCH_3_), 7.02 (1H, d, *J* = 8.9 Hz, H6), 7.11 (1H, d, *J* = 8.9 Hz, H7). ^13^C NMR (101 MHz, DMSO-*d6*) *δ*: 15.79, 56.95, 57.07, 109.05, 111.58, 120.07, 120.71, 142.13, 146.93, 148.30, 156.33. HPLC: CH_3_CN/H_2_O (60:40), ret. time: 2.23 min, peak area: 95.80%.

### 4-Chloro-*N*-(5,8-dimethoxy-2,3-dihydro[1,2,4]triazolo[3,4-*b*][1,3]benzothiazol-3-yl)benzamide (29)

Et_3_N (0.16 mL, 1.2 mmol) and *p*-chlorobenzoyl chloride (0.13 mL, 1.04 mmol) were added to a solution of **27** (0.22 g, 0.8 mmol) in dry DMF (6 mL) under nitrogen atmosphere. The reaction mixture was stirred for 2 h at 80 °C and, then, it was poured in ice/water obtaining a precipitate that was filtered and purified by flash chromatography eluting with CHCl_3_:MeOH (95:5), to give **29** as a purple solid (0.03 g, 10%). ^1^H NMR (400 MHz, DMSO-*d*_6_) *δ*: 3.39 (3H, s, OCH_3_), 3.95 (3H, s, OCH_3_), 7.09-7.15 (2H, m, H6 and H7), 7.71 (2H, d, *J* = 6.7 Hz, H3’ and H5’), 8.11 (2H, d, *J* = 6.7 Hz, H2’ and H6’). ^13^C NMR (101 MHz, DMSO-*d*_6_) *δ*: 57.02, 57.14, 109.15, 111.33, 120.10, 120.38, 129.30, 130.30, 13.38, 137.90, 142.56, 142.92, 148.01, 154.93, 166.58. HRMS: *m/z* calcd for C_17_H_13_ClN_4_O_3_S 389.048 [M + H^+^], found 389.047. HPLC: CH_3_CN/H_2_O + 0.1% FA (60:40), ret. time: 2.17 min, peak area: 97.06%.

### N-(5,8-dimethoxy[1,2,4]triazolo[3,4-*b*][1,3]benzothiazol-3-yl)-2,2-dimethylpropanamide (30)

The title compound was synthesized following the same procedure as used for the synthesis of compound **29**, using trimethylacetyl chloride (0.11 mL, 0.88 mmol) and dry toluene as solvent at 110 °C, overnight. After purification by flash chromatography eluting with CHCl_3_:MeOH (97:3), compound **30** was obtained as a grey solid (0.06 g, 20%). ^1^H NMR (400 MHz, DMSO-*d*_6_) *δ*: 1.29 (9H, s, CH_3_) 3.89 (3H, s, OCH_3_), 3.95 (3H, s, OCH_3_), 7.13 (1H, d, *J* =9.1 Hz, H7), 7.22 (1H, d, *J* =9.1 Hz, H6), 10.14 (1H, s, NH). ^13^C NMR (101 MHz, DMSO-*d*_6_) *δ*: 27.68, 57.18, 57.99, 109.25, 112.11, 120.40, 120.55, 143.45, 143.49, 148.22, 154.51, 156.13, 179.03. HPLC: CH_3_CN/H_2_O + 0.1 % FA (70:30), ret. time: 1.77 min, peak area: 99.76%.

### N-(5,8-dimethoxy[1,2,4]triazolo[3,4-*b*][1,3]benzothiazol-3-yl)acetamide (31)

The title compound was synthesized following the same procedure as used for the synthesis of compound **29**, using acetyl chloride (0.63 mL, 0.88 mmol) and dry toluene as solvent at 110 °C, overnight. The reaction mixture was extracted with EtOAc, the organic layers were washed with brine, dried over Na_2_SO_4_ and evaporated under reduced pressure, obtaining an oil that was purified by flash chromatography eluting with CHCl_3_:MeOH (97:3), to give **31** as a pink solid (0.05 g, 20%). ^1^H NMR (400 MHz, DMSO-*d*_6_) *δ*: 2.19 (3H, s, CH_3_), 3.87 (3H, s, OCH_3_), 3.90 (3H, s, OCH_3_), 6.81 (1H, d, *J* = 8.7 Hz, H7), 6.92 (1H, d, *J* = 8.6 Hz, H6), 12.49 (1H, s, NH). ^13^C NMR (101 MHz, DMSO-*d*_6_) *δ*: 23.22, 56.45, 56.88, 104.96, 108.99, 121.28, 140.12, 146.97, 148.23, 157.34, 169.78. HPLC: CH_3_CN/H_2_O + 0.1% FA (70:30), ret. time: 1.80 min, peak area: 95.05%.

#### Protein production

The proteins used in tables 1-3 were produced as described earlier. Dockerin constructs to allow enhanced activity assays for mono-ARTs (Table 4) were produced in *E. coli* with an N-terminal MBP and a C-terminal dockerin domain from *Hungateiclostridium thermocellum* Cel48s as described recently^46^.

Human PARP6 (Uniprot #Q2NL67-1) was cloned into a modified pFASTBac1 vector (Addgene #30116) with N-terminal 6X His-maltose binding protein (MBP) tag with a TEV protease site by SLIC cloning. Construct was sequence verified by dideoxy sequencing. Protein was produced as described before.^25^ Sf21 cells were transfected with bacmids using Fugene6 (Promega #E2693). V_0_ virus containing media was harvested after 7 days. These viruses were amplified to increase titre (V_1_). For protein production, Sf21 cells at a density of 1×10^6^ cells per ml and a pre-determined volume of V_1_ virus that induces growth arrest at this density was used. Cells were harvested after 72 hours of growth arrest and frozen at -20°C with lysis buffer (50 mM Hepes pH 7.4, 0.5 M NaCl, 10% glycerol, 0.5 mM TCEP and 10 mM imidazole) until needed.

Pellet was thawed and 0.2 mM Pefabloc was added to the suspension. The mixture was sonicated and then centrifuged at 16,000 rpm to separate soluble proteins from cellular debris. Supernatant was centrifuged again to remove any carry over debris. Supernatant was bound to HiTrap™ IMAC columns (Cytiva) and then washed with 4 column volume (CV) of lysis buffer and then wash buffer (50 mM Hepes pH 7.4, 0.5 M NaCl, 10% glycerol, 0.5 mM TCEP and 25 mM imidazole). Proteins were eluted in (50 mM Hepes pH 7.4, 0.5 M NaCl, 10% glycerol, 0.5 mM TCEP and 350 mM imidazole). Eluted proteins were loaded into MBP-trap columns and washed with size exclusion (SEC) buffer (30 mM Hepes pH 7.5, 0.35 M NaCl, 10% glycerol, 0.5 mM TCEP) and eluted in SEC buffer supplemented with 10 mM Maltose. TEV protease (1:30) molar ratio was used to cleave tags.^78^ A reverse IMAC step was used to separate tags and TEV protease from cleaved PARP6. A final SEC was performed, and fractions were pooled, concentrated and flash frozen. Identity of the purified protein was confirmed using MALDI-TOF analysis.

#### Activity assay

Inhibition experiments were performed using a homogenous assay measuring NAD+ consumption.^79–81^ Reactions were carried out in quadruplicates and IC_50_ curves were fitted using sigmoidal dose response curve (four variables) in GraphPad Prism version 8.02. For the compounds showing < 1 µM potency the experiment was repeated three times and pIC_50_±SEM was calculated. Assay conditions for PARP2, TNKS2, PARP10 and PARP15 (Tables 1-3) were are recently reported.^43^ The conditions for the proximity enhanced mono-ART essays (Table 4) were also as reported recently.^46^ PARP6 (400 nM) inhibition was measured using the standard buffer (50 mM sodium phosphate pH 7.0) for the enhanced activity assay using 500 nM NAD^+^ and 18 hour incubation in RT.

#### Crystallization and structure refinement

The crystallizations were carried out using a sitting drop vapour diffusion method at +20°C for PARP15, and at +4°C for PARP2 and TNKS2. A hanging drop vapour diffusion method at +20°C was used for PARP14. The compounds were dissolved in DMSO at 10 mM concentration and were used to obtain protein-inhibitor complex structures either by co-crystallization or soaking as stated specifically for each protein below. The protein and precipitant solutions were mixed at 2:1-1:2 ratios with the Mosquito crystallization robot (SPT Labtech) resulting in 160 - 500 nl droplets. The crystallization experiments were monitored using the RI54 imagers (Formulatrix) through the IceBear software.^82^

The inhibitors except **27** were co-crystallized with PARP15 as previously reported.^43^ A 10 mg/ml of PARP15 was mixed with the compound solution to reach approximately 700 µM concentration. 0.2 M NH_4_Cl pH 7.5, 16 – 20% (w/v) PEG 3350 was used precipitant solution. The crystals were cryoprotected with a solution containing 0.2 M NH_4_Cl and 30% (v/v) MPD. **27** was soaked to a PARP15 crystal with the cryoprotectant solution containing 1 mM **27** and incubated for 15 min at +20°C prior to cryo-freezing with liquid nitrogen.

Prior to crystallization, the TNKS2 ART domain (5.3 mg/ml) was mixed with 1:100 chymotrypsin and incubated for 2 hours at room temperature. The protein was then mixed with precipitant solution containing 100 mM Tris (pH 8.5), 200 mM lithium sulfate and 20-24% (w/v) PEG3350. Crystals formed within 2-3 days. TNKS2 crystals were soaked for 8-24 hours with 1 or 5 diluted in precipitant solution to a final concentration of approximately 1 mM compound in the crystallization droplets. The crystals were cryoprotected using precipitant solution containing 20% (v/v) glycerol.

A 15 mg/ml PARP14 was mixed with **1** to reach approximately 860 µM inhibitor concentration. 0.17 M NH_4_SO_4_, 15% (v/v) glycerol, 27% (v/v) PEG 4000 was used as a precipitant solution. Crystals were obtained in 2 days and were cryo-frozen with liquid nitrogen.

Compound **16** complex crystal structure with PARP2 was obtained using dry compound co-crystallization. 20 nl of 10 mM compound 10 were transferred to the crystallization plate and allowed to dry at 37°C before proceeding. 30 mg/ml PARP2 was then mixed with precipitant solution containing 100 mM Tris pH 9.5 and 20% PEG 3350. Crystals were cryoprotected with a solution containing 100 mM Tris pH9.0, 200 mM NaCl, 25 % PEG3350, 22 % glycerol and 100 µM compound **16**.

All datasets collected from PARP2, TNKS2, PARP14 and PARP15 crystals were processed with XDS^83^. Phases were solved by using molecular replacement with the programs MOLREP^84^ in CCP4i2^85^ or with Phaser^86^. The existing models having PDB ids 4TVJ^87^, 5OWS^88^, 3GOY^50^ and 3BLJ^22^ were used as search models for PARP2, TNKS2, PARP14 and PARP15, respectively. The models were built by using the Coot program^89^ and refined with Refmac5^90^ in CCP4i2. Data collections and refinement statistics are shown in Table S1.

#### Cell viability assay

Cell viability was assessed by colorimetric WST-1 (Cellpro-Roche, Sigma-Aldrich) assay following manufacturer’s instructions. Shortly, HEK293T cells were seeded at the density of 2,5×10^4^ cells per well in 96-well plate in 100 μl of Dulbecco’s modified Eagle’s media (DMEM, Biowest) supplemented with 10% fetal bovine serum (Biowest) and 1% of penicillin and streptomycin. Cells were allowed to growth for 18 h before adding compounds at the indicated concentrations (100 μM, 50 μM, and 10 μM). Also, DMSO and 10 mM hydroxyurea (Sigma) was used as an internal control for induced cell toxicity. Cells were grown for additional 24 h. After which WST-1 reagent was pipetted followed by 2 h incubation and absorbance measuring by Tecan Infinite M1000 or Tecan Spark (Tecan) plate reader. Assay was performed in triplicates, and repeated at least three times. Data was normalized to DMSO control.

#### PARP10 rescue assay

HeLa Flp-In T-REx-PARP10 and -PARP10-G888W cells were grown in DMEM medium supplemented with 10% heat-inactivated fetal calf serum at 37 °C in 5% CO_2_. For colony formation assays, 500 HeLa cells were seeded in 6-well culture plates. Once the cells adhered, protein expression was induced by adding 500 ng/mL doxycycline. Different concentrations of the indicated compounds were added to the cell culture medium as indicated in the figure. The cells were grown for 10-12 days and then stained using methylene blue. The number of colonies was assessed using ImageJ. EC_50_ curves were fitted using three variables in GraphPad Prism version 8.02.

#### *In vitro* ADME studies

All solvents and reagents were from Sigma-Aldrich Srl (Milan, Italy). Dodecane was purchased from Fluka (Milan, Italy). Pooled male donors 20 mg/mL HLM were from Merk-Millipore (Burlington, MA, USA). Milli-Q quality water (Millipore, Milford, MA, USA) was used. Hydrophobic filter plates (MultiScreen-IP, clear plates, 0.45 mm diameter pore size), 96-well microplates, and 96-well UV-transparent microplates were obtained from Merk-Millipore (Burlington, MA, USA).

#### UV/LC-MS methods

UV/LC-MS LC analyses for ADME studies were performed by UV/LC-MS with Agilent 1260 Infinity HPLC-DAD interfaced with an Agilent MSD 6130 (Agilent Technologies, Palo Alto, CA). Chromatographic separation was obtained using a Phenomenex Kinetex C18-100 Å column (150 × 4.6 mm) with 5 µm particle size and gradient elution with a binary solution; (eluent A: H2O, eluent B: ACN, both eluents were acidified with formic acid 0.1% v/v) at room temperature. The analysis started with 5% of B (from t= 0 to t= 1 min), then B was increased to 95% (from t=1 to t=10 min), then kept at 95% (from t= 10 to t=19 min) and finally return to 5% of eluent A in 1.0 min. The flow rate was 0.6 mL/min and injection volumes were 10 µL.

#### Water solubility

Each solid compound (1 mg) was added to 1 mL of distilled water. Each sample was mixed at room temperature in a shaker water bath 24h. The resulting suspension was filtered through a 0.45 µm nylon filter (Acrodisc) and the solubilized compound was quantified in triplicate using UV/LC-MS method reported above, by comparison with the appropriate calibration curve that was obtained from samples of the compound dissolved in methanol at different concentrations.^91^

#### Parallel artificial membrane permeability assay (PAMPA)

Each ‘donor solution’ was prepared from a solution of the appropriate compound (DMSO, 1 mM) diluted with phosphate buffer (pH 7.4, 0.025 M) up to a final concentration of 500 µM. Filters were coated with 10 µL of 1% dodecane solution of phosphatidylcholine or 5 µL of brain polar lipid solution (20 mg/mL 16% CHCl_3_, 84% dodecane) prepared from CHCl_3_ solution 10% w/v, for intestinal permeability and BBB permeability, respectively. Donor solution (150 µL) was added to each well of the filter plate and to each well of the acceptor plate were added 300 µL of solution (50% DMSO in phosphate buffer). The sandwich plate was assembled and incubated for 5 h at room temperature. After the incubation time, the plates were separated, and samples were taken from both the donor and acceptor wells and the amount of compound was measured by UV/LC-MS. All compounds were tested in three independent experiments. Permeability (P_app_) was calculated according to the following equation obtained from literature ^92,93^ with some modification in order to obtain permeability values in cm/s:

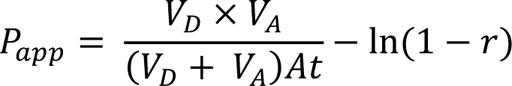

where V_A_ is the volume in the acceptor well, V_D_ is the volume in the donor well (cm^3^), A is the “effective area” of the membrane (cm^2^), t is the incubation time (s) and r the ratio between drug concentration in the acceptor and equilibrium concentration of the drug in the total volume (V_D_+V_A_). Drug concentration is estimated by using the peak area integration. Membrane retentions (%) were calculated according to the following equation:

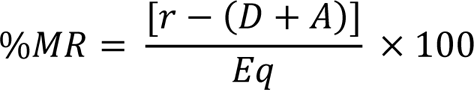

where r is the ratio between drug concentration in the acceptor and equilibrium concentration, D, A, and Eq represented drug concentration in the donor, acceptor, and equilibrium solution, respectively.

#### Metabolic Stability in HLM (Human liver microsomes)

Each compound in DMSO solution was incubated at 37°C for 1 hour in phosphate buffer (25 mM pH 7.4), human liver microsomal protein (0.2 mg/mL) and in presence of an NADPH regenerating system (NADPH 0.2 mM, NADPH^+^ 1 mM, D-glucose-6-phosphate 4 mM, 4 unit/mL glucose-6-phosphate dehydrogenase) in MgCl_2_ 48 mM at a final volume of 500 µL. The reaction was stopped by cooling in ice and quenched adding 1.0 mL of acetonitrile. The reaction mixtures were then centrifuged (4000 rpm for 10 min), the supernatant was taken, dried under nitrogen flow, suspended in 100 µL of methanol and the parent drug and metabolites were subsequently determined by UV/LC-MS. The percentage of not metabolized compound was calculated by comparison with reference solutions. For each compound, the determination was performed in three independent experiments.

#### Stability tests

For the stability measurements in polar solvents each compound was dissolved at room temperature in MeOH or PBS (0.025 M, pH 7.4) up to a final concentration of 500 µM. Aliquot samples (20 µL) were taken at fixed time points (0.0, 4.0, 8.0, 24.0 h) and were analysed by UV/LC-MS. For each compound, the determination was performed in three independent experiments.

To test the stability in human plasma the incubation mixture (total volume of 2.0 mL) was constituted by following components: pooled human plasma (1.0 mL, 55.7 mg protein/mL)^94^, HEPES buffer (0.9 mL, 25 mM, 140 mM NaCl pH 7.4) and 0.1 mL of each compound in DMSO (2.0 mM). The solution was mixed in a test tube that was incubated at 37 °C. At set time points (0.0, 0.08, 0.25, 0.50, 1.0, 2.0, 4.0, 8.0 and 24.0 h), samples of 50 µL were taken, mixed with 450 µL of cold acetonitrile and centrifuged at 5000 rpm for 15 min.^95^ The supernatant was removed and analysed by UV/LC-MS. For each compound, the determination was performed in three independent experiments.

## DATA AVAILABILITY

Atomic coordinates and structure factors have been deposited to the Protein Data Bank under accession numbers also mentioned in the tables 7R3Z, 7R3L, 7R3O, 7R4A, 7R5X, 7R5D, 7Z1W, 7Z1Y, 7R59, 7Z1V, 7Z41, 7Z2O and 7Z2Q. Raw diffraction images are available at IDA (https://doi.org/10.23729/0b11fe27-a545-48b0-a953-292d1e1e1d38).

## ACKNOWLEDGEMENT

We thank Chiara Bosetti for measuring some dose responses with the TNKS2 enzyme. The use of the facilities and expertise of the Biocenter Oulu Structural Biology core facility (a member of Biocenter Finland, Instruct-ERIC Centre Finland and FINStruct), Proteomics and Protein Analysis core facility (a member of Biocenter Finland) and Biocenter Oulu sequencing center are gratefully acknowledged. We thank the staff members of DLS, ESRF and MAXIV.

## ABBREVIATIONS USED

ADPr: ADP-ribose
BBB: blood brain barrier
GI: gastrointestinal
MAR: mono-ADP-ribosylation
PAMPA: parallel artificial membrane permeability assay
PAR: poly-ADP-ribosylation
TBT: [1,2,4]triazolo[3,4-*b*]benzothiazole
TNKS: tankyrase

