## Supplementary information for "[1,2,4]Triazolo[3,4-*b*]benzothiazole scaffold as versatile nicotinamide mimic allowing nanomolar inhibition of different PARP enzymes"

**CONTENT**

**Table S1**. Crystallography data processing and refinement statistics.

**Figure S1.** WST-1 cell toxicity analysis of compounds.

**Figure S2.** Binding modes of OUL40 analogs to TNKS2 and PARP15.

**Figure S3.** Examples of $IC_{50}$ measurements.

**Synthesis methods and characterization data**

**Example of HPLC analysis of the target compounds**

**Example of $^{1}$HNMR and $^{13}$CNMR spectra of the target compounds**

**Table S2.** SMILES and biochemical data of compounds 1-31.

**Table 1. Data collection and refinement statistics.**

| | TNKS2-**1** (PDB 7R3Z) | PARP14-**1** (PDB 7R3L) | PARP15-**1** (PDB 7R3O) | PARP15-**8** (PDB 7R4A) |
|---|---|---|---|---|
| **Data collection** | | | | |
| Beamline | DLS, I04 | ESRF, ID30A-1 | DLS, I04 | MAX IV, BioMax |
| Wavelength (Å) | 0.97949 | 0.96600 | 0.97950 | 0.979957 |
| Space group | $C222_1$ | C2 | $P2_12_12_1$ | $P2_12_12_1$ |
| Cell dimensions | | | | |
| *a*, *b*, *c* (Å) | 91.96, 97.49, 117.90 | 81.88, 83.75, 80.03 | 45.24, 68.54, 158.57 | 45.46, 68.74, 159.23 |
| α, β, γ (°) | 90, 90, 90 | 90, 115.7, 90 | 90, 90, 90 | 90, 90, 90 |
| Resolution (Å) | 50 – 2.25 | 53 – 1.999 | 50 – 2.20 | 50 – 1.90 |
| Outer shell (Å) | 2.31 – 2.25 | 2.033 – 1.999 | 2.26 – 2.20 | 1.95 – 1.90 |
| No. unique reflections | 25467 (1844) | 32725 (1587) | 25834 (1893) | 40198 (2893) |
| $R_{merge}$ | 0.123 (1.51) | 0.088 (0.534) | 0.185 (0.942) | 0.145 (1.25) |
| Mean $I/\sigma I$ | 10.7 (1.96) | 10.3 (2.30) | 7.80 (1.94) | 8.1 (1.80) |
| CC ½ (%) | 99.6 (74.5) | 99.7 (84.2) | 99.4 (79.1) | 99.6 (46.2) |
| Completeness (%) | 99.9 (99.9) | 99.2 (99.2) | 99.9 (100) | 99.9 (99.9) |
| Redundancy | 6.5 (6.8) | 3.1 (3.1) | 6.6 (6.7) | 6.4 (6.6) |
| **Refinement** | | | | |
| $R_{work}$ / $R_{free}$ | 0.197 / 0.245 | 0.202 / 0.237 | 0.205 / 0.242 | 0.233 / 0.272 |
| No. atoms | | | | |
| Protein | 3352 | 2683 | 3193 | 3183 |
| Ligand/ion | 54 | 41 | 17 | 18 |
| Water | 78 | 145 | 88 | 150 |
| *B*-factors | | | | |
| Protein | 46.1 | 24.7 | 34.2 | 33.5 |
| Ligand/ion | 49.04 | 33.2 | 42.6 | 35.0 |
| Water | 39.3 | 25.7 | 26.0 | 36.6 |
| R.m.s. deviations | | | | |
| Bond lengths (Å) | 0.0065 | 0.0073 | 0.0080 | 0.0088 |
| Bond angles (°) | 1.448 | 1.418 | 1.515 | 1.528 |
| Ramachandran plot (%) | | | | |
| Favored | 97.8 | 98.7 | 97.2 | 97.9 |
| Allowed | 2.2 | 1.3 | 2.8 | 2.1 |
| Outliers | 0 | 0 | 0 | 0 |

Values in parentheses are for highest-resolution shell.

**Table 1. Data collection and refinement statistics. Contd.**

| | TNKS2-**3** (PDB 7R5X) | PARP15-**6** (PDB 7R5D) | PARP15-**7** (PDB 7Z1W) | PARP15-**16** (PDB 7Z1Y) |
|---|---|---|---|---|
| **Data collection** | | | | |
| Beamline | DLS I03 | ESRF, ID30A-1 | ESRF, ID30A-1 | ESRF, ID30A-1 |
| Wavelength (Å) | 0.92023 | 0.96546 | 0.96546 | 0.96546 |
| Space group | $C222_1$ | $P2_12_12_1$ | $P2_12_12_1$ | $P2_12_12_1$ |
| Cell dimensions | | | | |
| *a*, *b*, *c* (Å) | 91.94, 96.99, 119.44 | 45.37, 68.80, 160.94 | 45.33, 68.47, 159.91 | 45.23, 68.33, 158.65 |
| $\alpha$, $\beta$, $\gamma$ (°) | 90, 90, 90 | 90, 90, 90 | 90, 90, 90 | 90, 90, 90 |
| Resolution (Å) | 48.49 - 2.0 | 50 – 2.15 | 50 – 1.90 | 50 – 1.75 |
| Outer shell (Å) | 2.072 - 2.0 | 2.21 – 2.15 | 1.95 – 1.90 | 1.80 – 1.75 |
| No. unique reflections | 36379 (3591) | 27222 (1843) | 39946 (2903) | 50369 (3652) |
| $R_{merge}$ | 0.0953 (0.722) | 0.057 (0.292) | 0.112 (0.855) | 0.096 (0.968) |
| Mean $I/\sigma I$ | 15.2 (3.46) | 14.5 (3.73) | 9.0 (1.81) | 12.6 (2.25) |
| CC ½ (%) | 99.8 (87.1) | 99.7 (93.5) | 99.6 (70.6) | 99.8 (74.7) |
| Completeness (%) | 99.93 (100.0) | 96.4 (91.1) | 99.5 (99.9) | 99.6 (99.2) |
| Redundancy | 7.9 (8.4) | 3.7 (2.8) | 4.9 (4.7) | 6.3 (5.9) |
| **Refinement** | | | | |
| $R_{work}$ / $R_{free}$ | 0.189 / 0.223 | 0.203 / 0.249 | 0.180 / 0.212 | 0.226 / 0263 |
| No. atoms | | | | |
| Protein | 3408 | 3182 | 3208 | 3202 |
| Ligand/ion | 54 | 19 | 17 | 17 |
| Water | 169 | 135 | 202 | 121 |
| *B*-factors | | | | |
| Protein | 37.9 | 31.9 | 29.4 | 27.9 |
| Ligand/ion | 42.5 | 51.7 | 44.1 | 47.4 |
| Water | 37.8 | 31.2 | 33.1 | 29.8 |
| R.m.s. deviations | | | | |
| Bond lengths (Å) | 0.013 | 0.0072 | 0.0076 | 0.0088 |
| Bond angles (°) | 1.7 | 1.50 | 1.46 | 1.58 |
| Ramachandran plot | | | | |
| Favored | 98.5 | 97.2 | 98.5 | 97.2 |
| Allowed | 1.5 | 2.5 | 1.5 | 2.8 |
| Outliers | 0 | 0.3 | 0 | 0 |

Values in parentheses are for highest-resolution shell.

**Table 1. Data collection and refinement statistics. Contd.**

| | PARP2-**16** (PDB 7R59) | PARP15-**11** (PDB 7Z1V) | PARP15-**14** (PDB 7Z41) | PARP15-**13** (PDB 7Z2O) | PARP15-**27** (PDB 7Z2Q) |
|---|---|---|---|---|---|
| **Data collection** | | | | | |
| Beamline | DLS, I03 | ESRF, ID30A-1 | DLS, I04 | DLS, I03 | ESRF, ID30A-1 |
| Wavelength | 0.97625 | 0.96546 | 0.97950 | 0.9763 | 0.96546 |
| Space group | $P2_12_12_1$ | $P2_12_12_1$ | $P2_12_12_1$ | $P2_12_12_1$ | $P2_12_12_1$ |
| Cell dimensions | | | | | |
| *a*, *b*, *c* (Å) | 58.38, 67.82, 86.69 | 45.14, 68.38, 159.16 | 45.28, 68.61, 158.84 | 45.31, 68.80, 160.21 | 45.24, 68.67, 159.88 |
| $\alpha$, $\beta$, $\gamma$ (°) | 90, 90, 90 | 90, 90, 90 | 90, 90, 90 | 90, 90, 90 | 90, 90, 90 |
| Resolution (Å) | 50-2.00 | 50 – 1.50 | 50 – 2.10 | 50 – 1.50 | 50 – 2.00 |
| Outer shell (Å) | 2.05-2.00 | 1.54 – 1.50 | 2.10 – 2.15 | 1.54 – 1.50 | 2.05 – 2.00 |
| No. unique reflections | 23900 (1743) | 79300 (5774) | 29760 (2184) | 81152 (5903) | 34370 (2523) |
| $R_{merge}$ | 0.235 (1.405) | 0.082 (0.798) | 0.181 (0.714) | 0.050 (1.279) | 0.148 (0.771) |
| Mean $I/\sigma I$ | 9.47 (1.90) | 9.6 (1.79) | 6.7 (1.7) | 26.1 (2.09) | 6.3 (1.84) |
| CC ½ (%) | 99.6 (72.6) | 99.7 (69.1) | 99.4 (78.5) | 100 (81.0) | 98.9 (70.2) |
| Completeness (%) | 100 (100) | 99.3 (99.6) | 99.9 (99.9) | 100 (100) | 99.4 (99.8) |
| Redundancy | 13.4 (13.9) | 4.2 (4.2) | 6.6 (6.3) | 13.1 (13.1) | 4.0 (4.2) |
| **Refinement** | | | | | |
| $R_{work}$ / $R_{free}$ | 0.187 / 0.238 | 0.143 / 0.189 | 0.192 / 0.232 | 0.148 / 0.187 | 0.193 / 0.216 |
| No. atoms | | | | | |
| Protein | 2764 | 3253 | 3203 | 3262 | 3202 |
| Ligand/ion | 19 | 22 | 20 | 18 | 21 |
| Water | 199 | 318 | 202 | 290 | 196 |
| *B*-factors | | | | | |
| Protein | 27.3 | 21.4 | 33.1 | 28.6 | 28.1 |
| Ligand/ion | 23.9 | 25.4 | 38.6 | 32.8 | 29.0 |
| Water | 28.9 | 31.2 | 34.6 | 36.1 | 30.9 |
| R.m.s. deviations | | | | | |
| Bond lengths (Å) | 0.0086 | 0.0097 | 0.0070 | 0.0087 | 0.0062 |
| Bond angles (°) | 1.494 | 1.497 | 1.481 | 1.472 | 1.419 |
| Ramachandran plot | | | | | |
| Favored | 97.1 | 98.5 | 97.7 | 98.5 | 97.7 |
| Allowed | 2.6 | 1.5 | 2.3 | 1.5 | 2.0 |
| Outliers | 0.3 | 0 | 0 | 0 | 0.3 |

Values in parentheses are for highest-resolution shell.

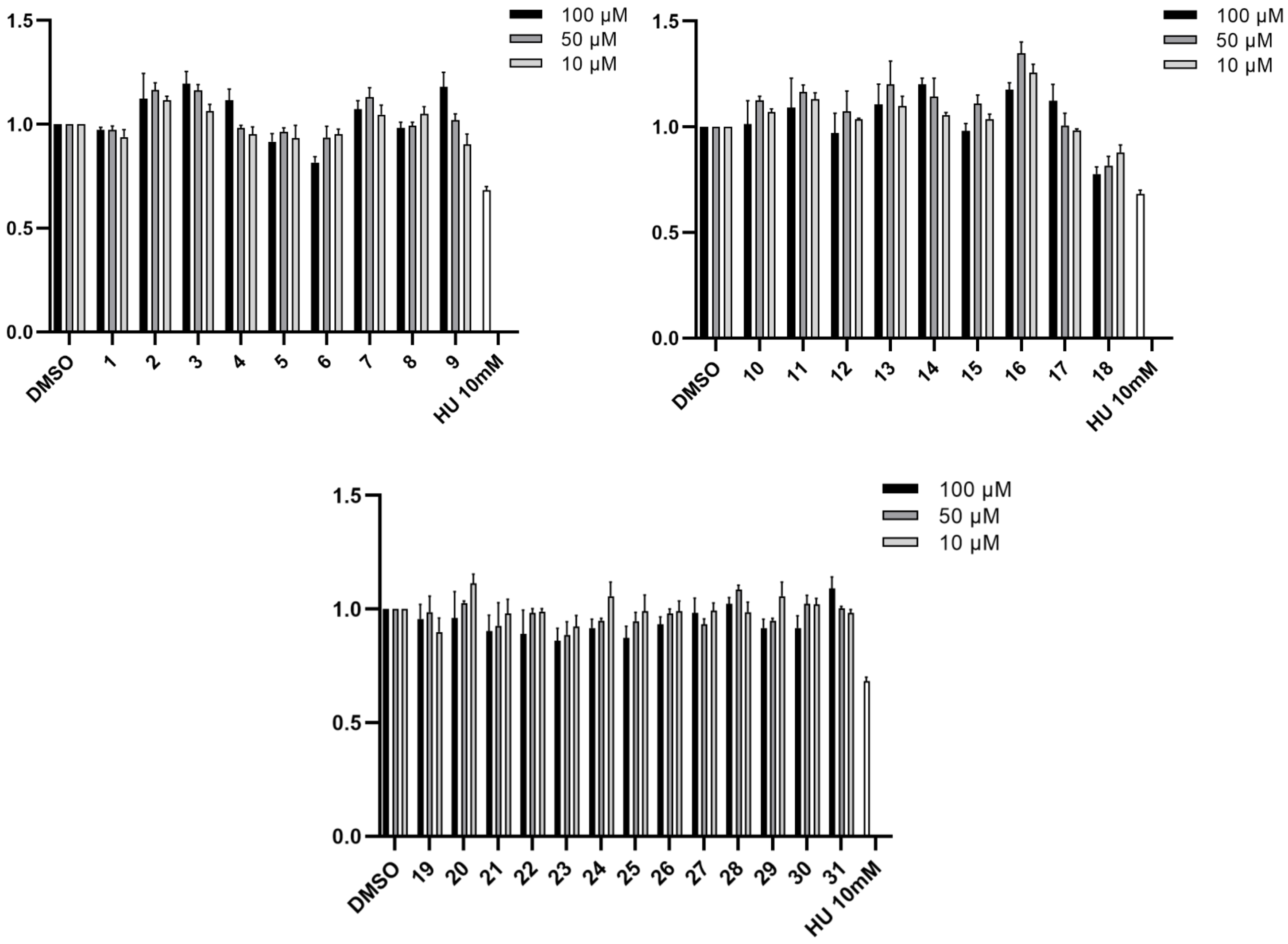


**Figure S1.** Graph represents cell viability used to determine compound **1-31** toxicity in HEK293T cells. Hydroxyurea (HU, 10 mM) was used as toxicity reference. Data was normalized to DMSO. The bars represent the means ± SEM of three experiment repeats.

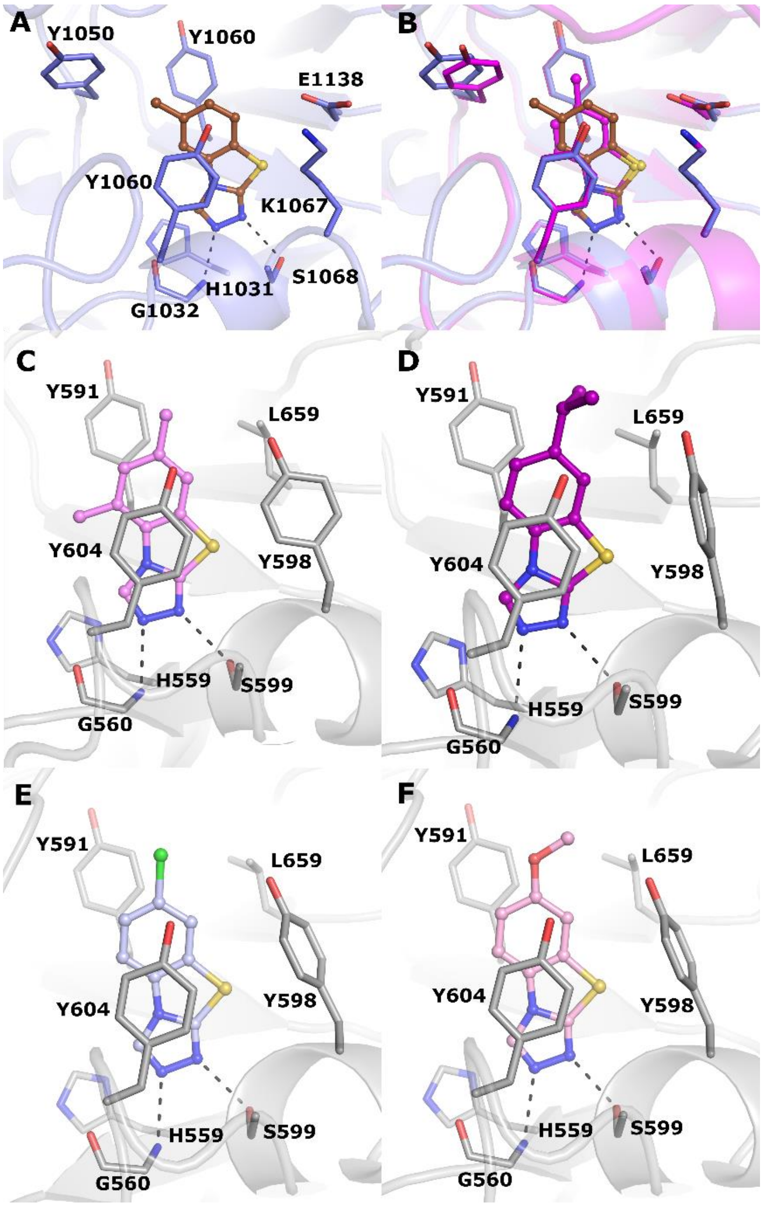


**Figure S2.** Binding modes of OUL40 analogs to TNKS2 and PARP15. **(A)** TNKS2 crystal structure in complex with **3**. The ligand is colored in brown **(B)** Superimposition of the TNKS2 complex structures of **3** and **1**. The complex structure of **1** is colored in magenta. **(C)** PARP15 crystal structure in complex with **8**. The ligand is colored in violet (**D**) PARP15 crystal structure in complex with **6**. The ligand is colored in purple (**E**) PARP15 crystal structure in complex with **7**. The ligand is colored in light blue (**F**) PARP15 crystal structure in complex with **11**. The ligand is colored in pink. Hydrogen bonds are indicated in black dash lines.

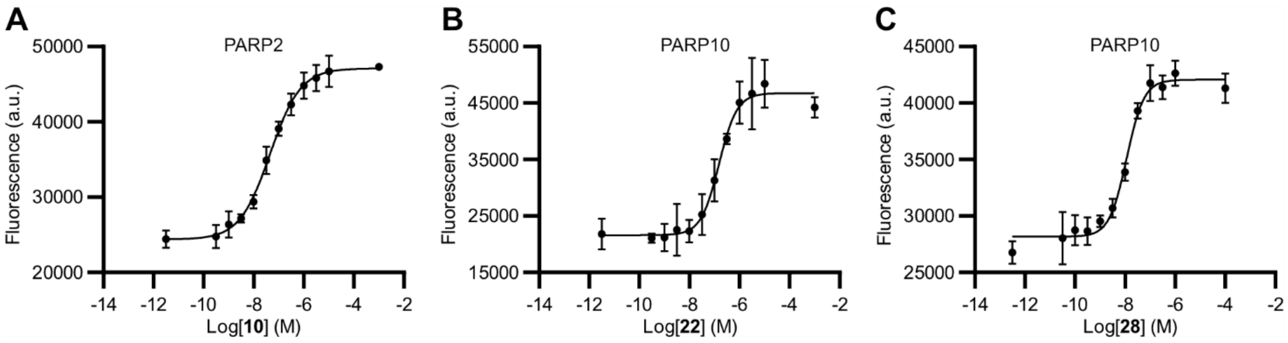


**Figure S3.** $IC_{50}$ curve examples of the best compounds **16**, **21** and **27**. **(A)** $IC_{50}$ curve of **16** measured with PARP2. **(B)** $IC_{50}$ curve of **21** measured with PARP10. **(C)** $IC_{50}$ curve of **27** measured with PARP10. Data shown are mean from n=4 measurements ± standard deviation. Controls were placed two logarithmic units above the highest or below the lowest compound concentration measured.

### Examples of HPLC analysis of target compounds

**6**

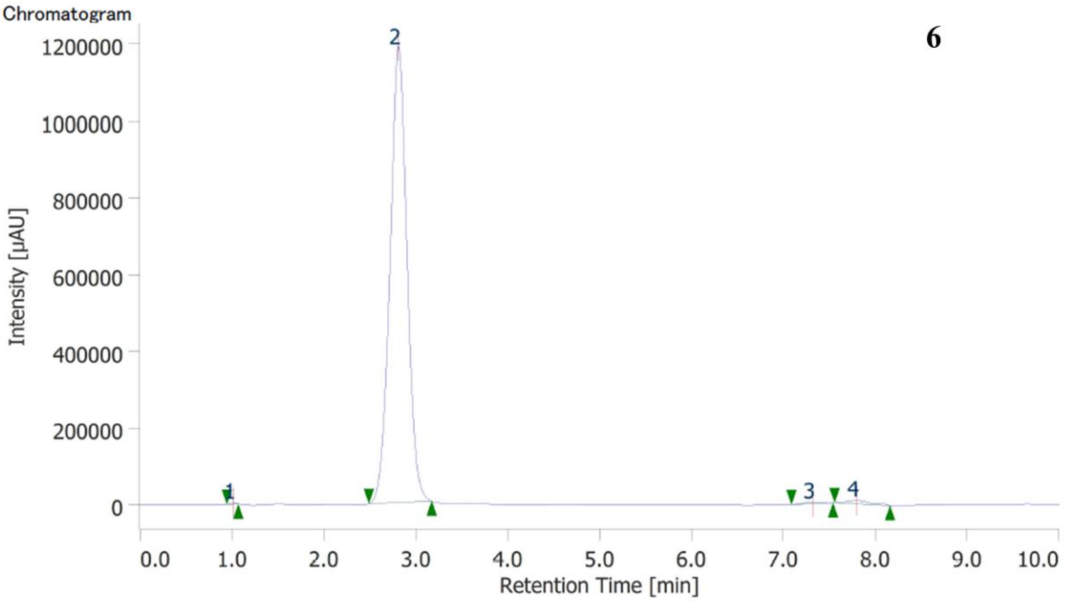


Peak Information

| # | Peak Name | CH | tR [min] | Area [μV·sec] | Height [μV] | Area% | Height% | Quantity | Resolution |
|---|---|---|---|---|---|---|---|---|---|
| 1 | Peak-001 | 9 | 1.017 | 13822 | 3229 | 0.089 | 0.268 | N/A | 7.946 |
| 2 | Peak-002 | 9 | 2.810 | 15238258 | 1188136 | 98.491 | 98.723 | N/A | 11.766 |
| 3 | Peak-003 | 9 | 7.317 | 55013 | 3680 | 0.356 | 0.306 | N/A | 1.077 |
| 4 | Peak-004 | 9 | 7.797 | 164654 | 8455 | 1.064 | 0.703 | N/A | N/A |

15

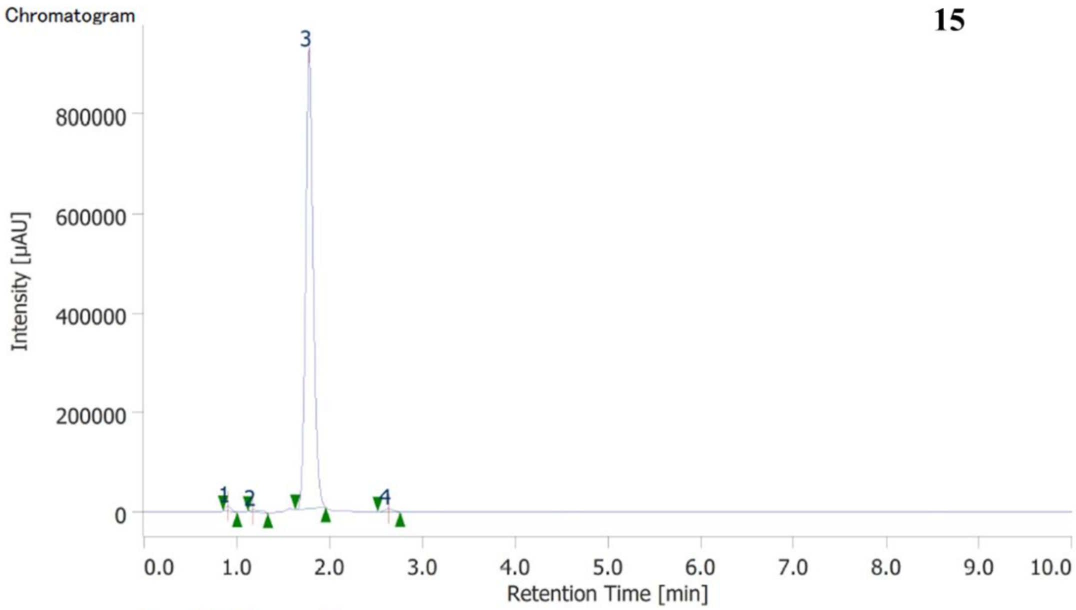


### Peak Information

| # | Peak Name | CH | tR [min] | Area [µV·sec] | Height [µV] | Area% | Height% | Quantity | Resolution |
|---|---|---|---|---|---|---|---|---|---|
| 1 | Unknown | 9 | 0.903 | 41683 | 9834 | 0.792 | 1.043 | N/A | 1.399 |
| 2 | Unknown | 9 | 1.180 | 30953 | 3429 | 0.588 | 0.364 | N/A | 2.902 |
| 3 | Unknown | 9 | 1.783 | 5150668 | 923460 | 97.882 | 97.934 | N/A | 5.508 |
| 4 | Unknown | 9 | 2.637 | 38840 | 6216 | 0.738 | 0.659 | N/A | N/A |

**21**

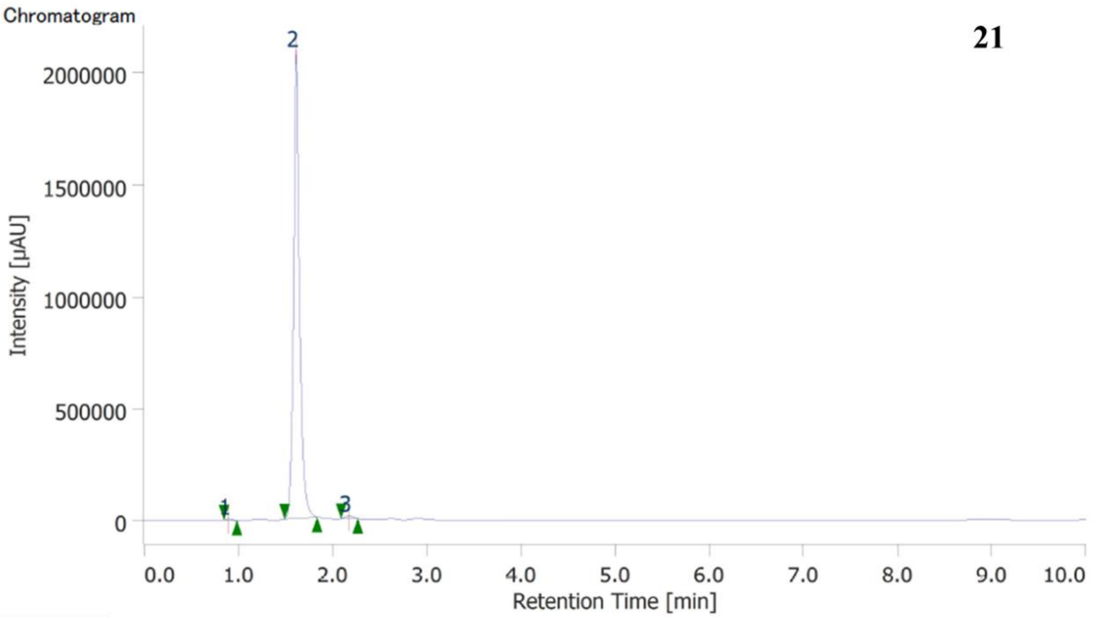


### Peak Information

| # | Peak Name | CH | tR [min] | Area [µV·sec] | Height [µV] | Area% | Height% | Quantity | Resolution |
|---|---|---|---|---|---|---|---|---|---|
| 1 | Unknown | 9 | 0.893 | 23152 | 5750 | 0.251 | 0.272 | N/A | 6.849 |
| 2 | Unknown | 9 | 1.613 | 9125956 | 2096449 | 98.979 | 99.166 | N/A | 4.155 |
| 3 | Unknown | 9 | 2.173 | 70968 | 11885 | 0.770 | 0.562 | N/A | N/A |

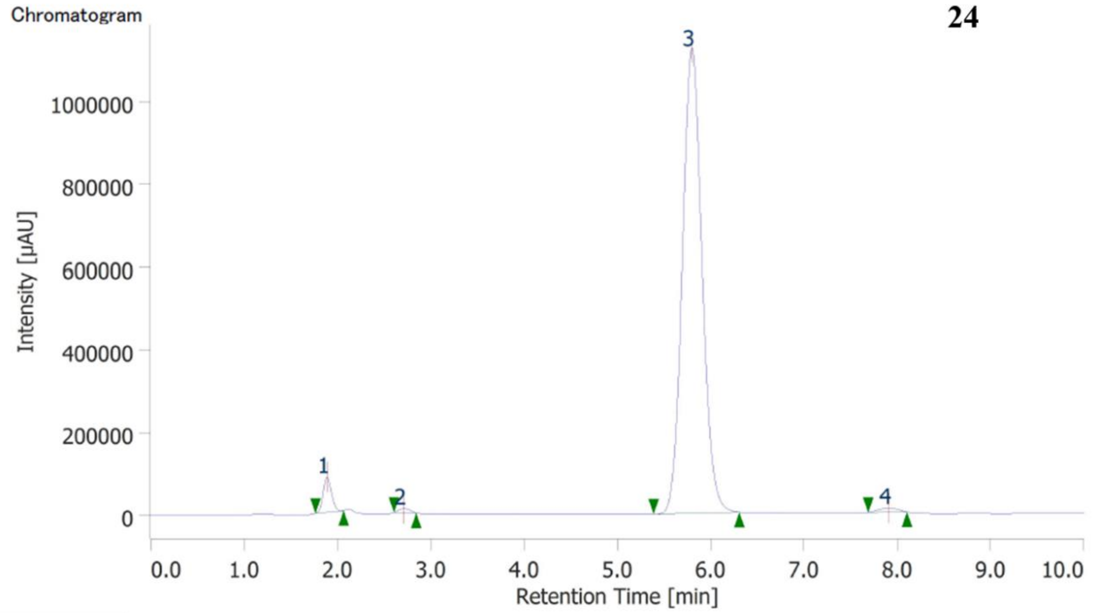


### Peak Information

| # | Peak Name | CH | tR [min] | Area [μV·sec] | Height [μV] | Area% | Height% | Quantity | Resolution |
|---|---|---|---|---|---|---|---|---|---|
| 1 | Unknown | 9 | 1.890 | 526042 | 85238 | 3.057 | 6.925 | N/A | 4.011 |
| 2 | Unknown | 9 | 2.710 | 83441 | 9870 | 0.485 | 0.802 | N/A | 9.883 |
| 3 | Unknown | 9 | 5.800 | 16459689 | 1125846 | 95.644 | 91.471 | N/A | 5.357 |
| 4 | Unknown | 9 | 7.907 | 140164 | 9864 | 0.814 | 0.801 | N/A | N/A |

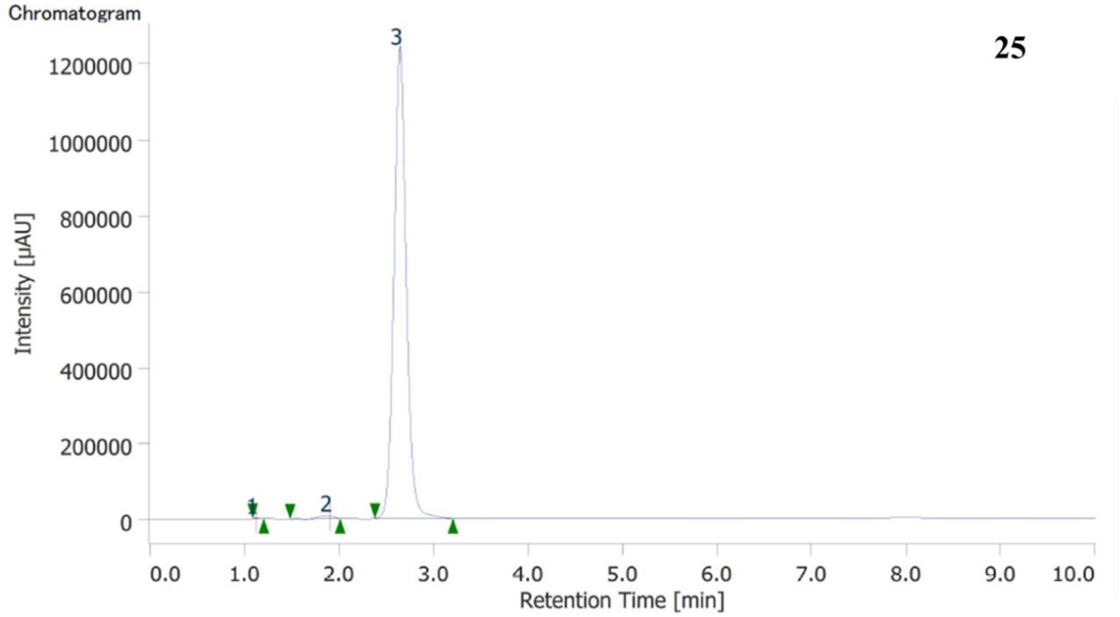


### Peak Information

| # | Peak Name | CH | tR [min] | Area [μV·sec] | Height [μV] | Area% | Height% | Quantity | Resolution |
|---|---|---|---|---|---|---|---|---|---|
| 1 | Unknown | 9 | 1.123 | 6608 | 1983 | 0.059 | 0.159 | N/A | 3.811 |
| 2 | Unknown | 9 | 1.903 | 93277 | 7554 | 0.830 | 0.604 | N/A | 2.689 |
| 3 | Unknown | 9 | 2.647 | 11136319 | 1240213 | 99.111 | 99.237 | N/A | N/A |

27

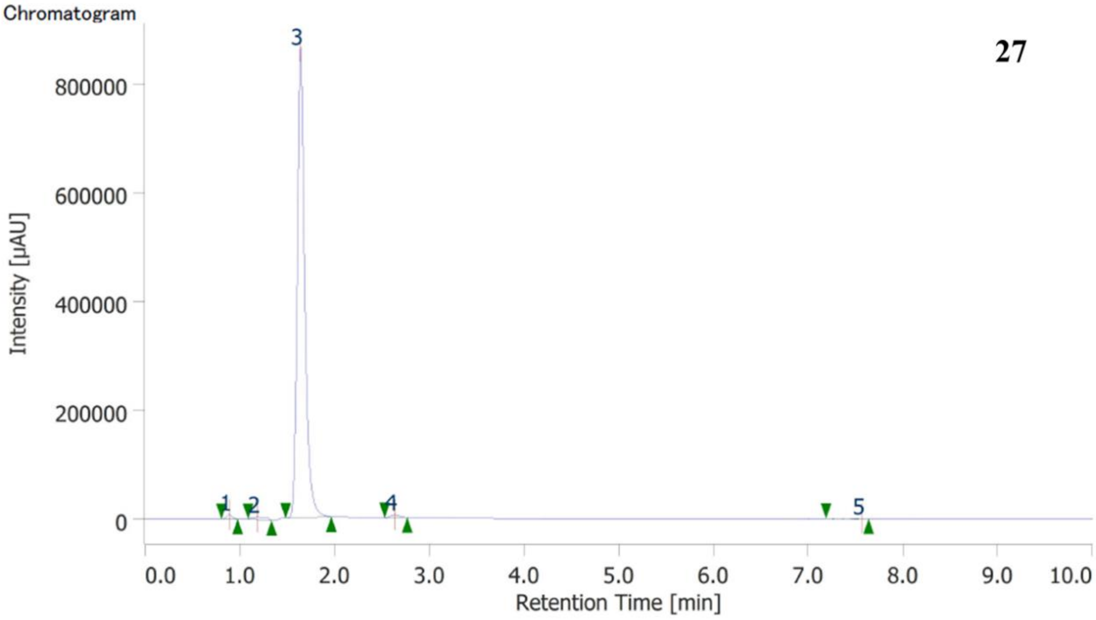


**Peak Information**

| # | Peak Name | CH | tR [min] | Area [µV·sec] | Height [µV] | Area% | Height% | Quantity | Resolution |
|---|---|---|---|---|---|---|---|---|---|
| 1 | Unknown | 9 | 0.890 | 31288 | 7812 | 0.632 | 0.885 | N/A | 1.431 |
| 2 | Unknown | 9 | 1.190 | 46221 | 4465 | 0.933 | 0.506 | N/A | 1.994 |
| 3 | Unknown | 9 | 1.640 | 4843949 | 865625 | 97.788 | 98.047 | N/A | 6.366 |
| 4 | Unknown | 9 | 2.637 | 31988 | 4939 | 0.646 | 0.559 | N/A | 12.250 |
| 5 | Unknown | 9 | 7.567 | 100 | 25 | 0.002 | 0.003 | N/A | N/A |

**31**

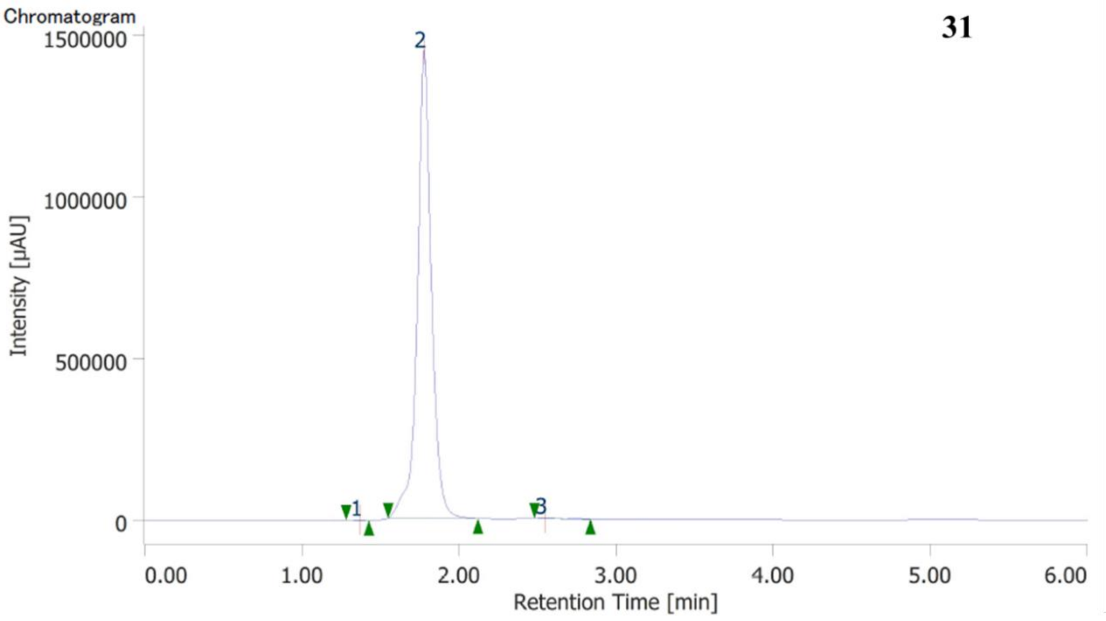


Peak Information

| # | Peak Name | CH | tR [min] | Area [µV·sec] | Height [µV] | Area% | Height% | Quantity | Resolution |
|---|---|---|---|---|---|---|---|---|---|
| 1 | Unknown | 9 | 1.370 | 6517 | 1460 | 0.072 | 0.100 | N/A | 3.008 |
| 2 | Unknown | 9 | 1.777 | 9056710 | 1450053 | 99.764 | 99.758 | N/A | 5.186 |
| 3 | Unknown | 9 | 2.547 | 14870 | 2063 | 0.164 | 0.142 | N/A | N/A |

**Example of $^{1}$HNMR and $^{13}$CNMR spectra of target compounds**

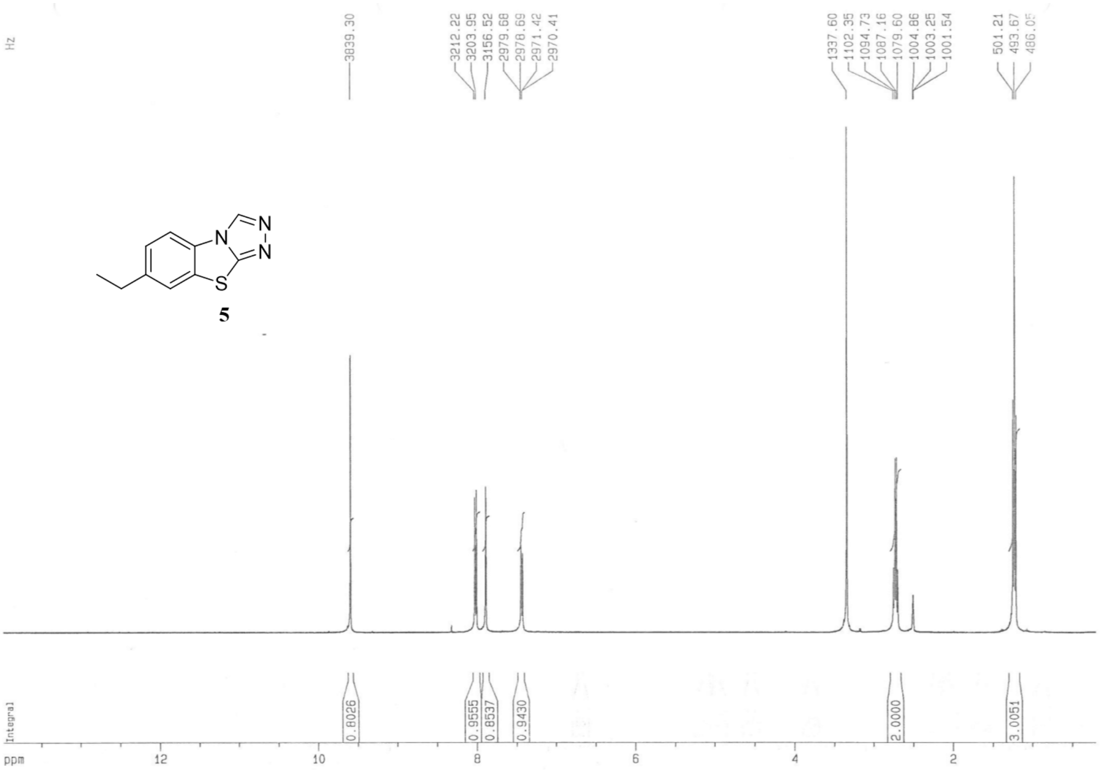

**5**

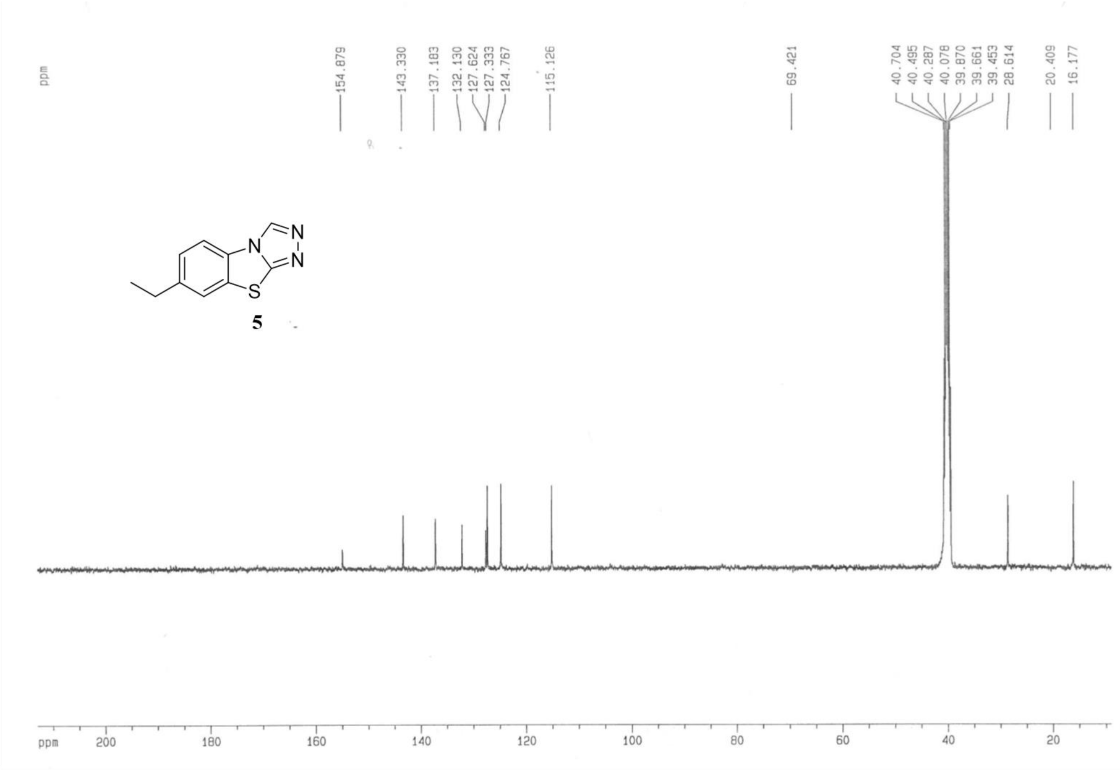
ppm
154.879
143.330
137.183
132.130
127.624
127.333
124.767
115.126
69.421
40.704
40.495
40.287
40.078
39.870
39.661
39.453
28.614
20.409
16.177
N
N
N
S
5
ppm 200 180 160 140 120 100 80 60 40 20

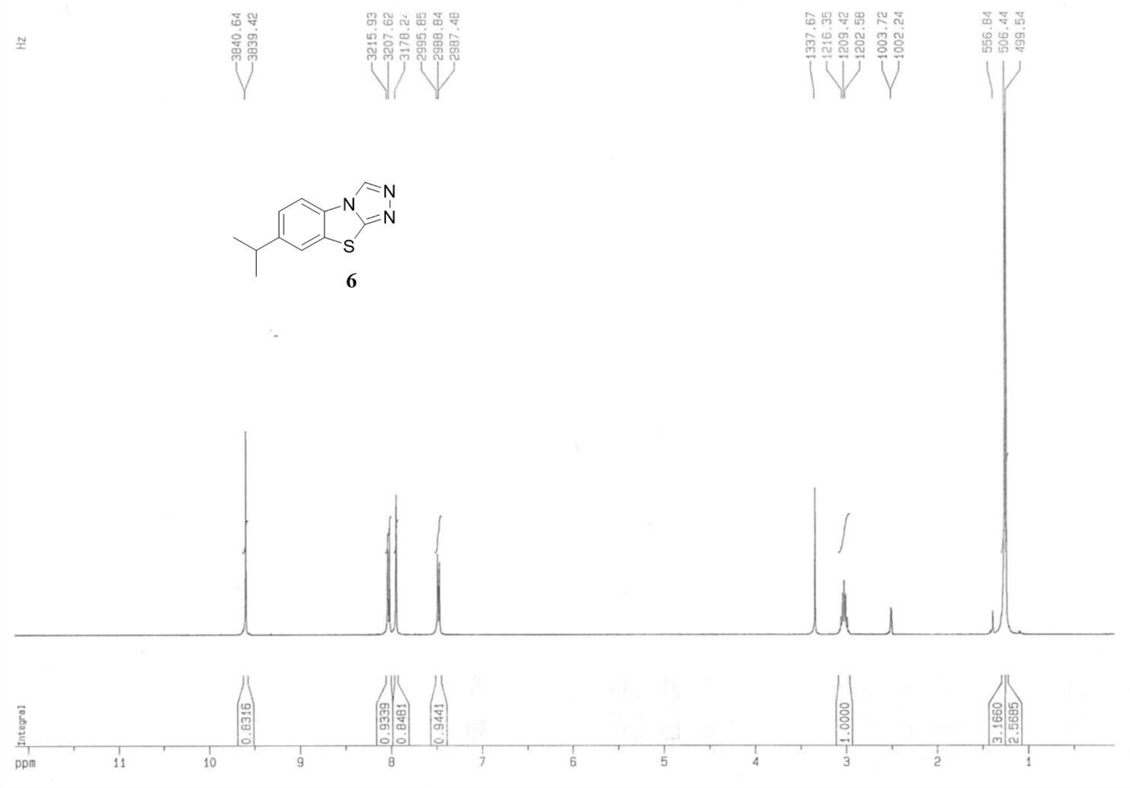
Hz
3840.64
3839.42
3215.93
3207.62
2995.85
2988.84
2987.48
1337.67
1216.35
1209.42
1202.58
1003.72
1002.24
556.84
506.44
499.54
N
N
N
S
6
Integral
0.8316
0.9339
0.8481
0.9441
1.0000
3.1660
2.5685
ppm 11 10 9 8 7 6 5 4 3 2 1

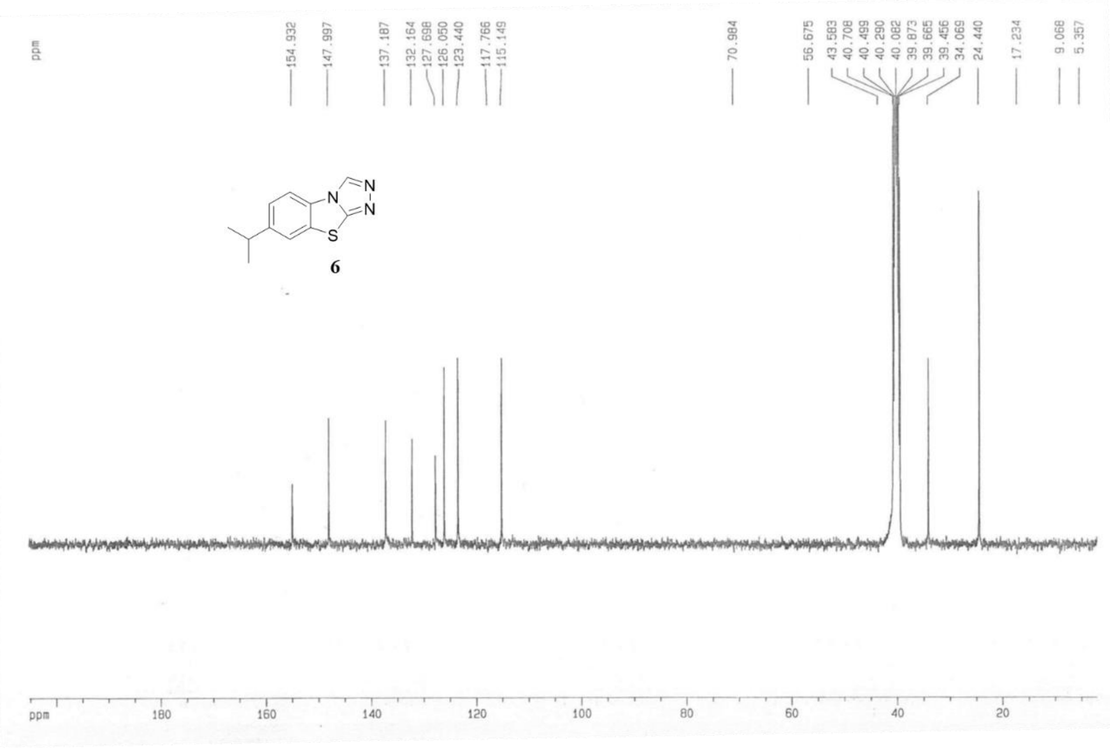
ppm
154.932
147.997
137.187
132.164
127.698
126.050
123.440
117.766
115.149
70.984
56.675
43.583
40.708
40.499
40.290
40.082
39.873
39.665
39.456
34.069
24.440
17.234
9.068
5.357
N
N
N
S
6
ppm 180 160 140 120 100 80 60 40 20

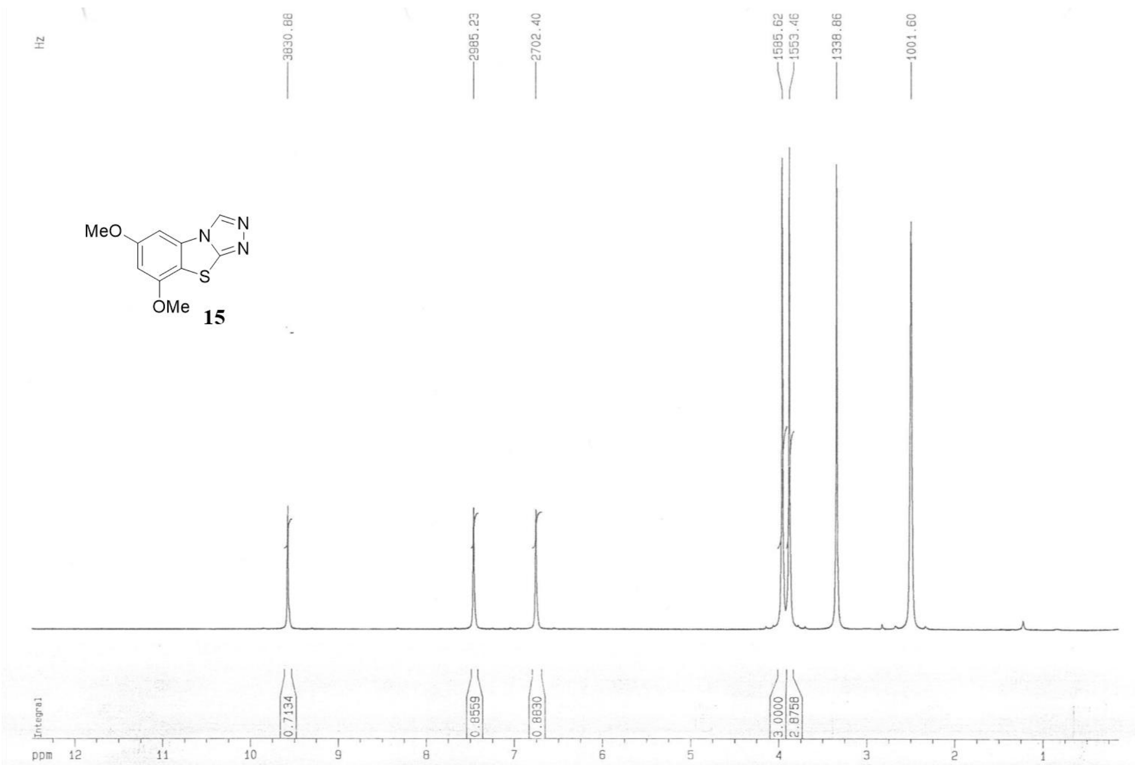
Hz
3830.88
2985.23
2702.40
1585.62
1553.46
1338.86
1001.60
MeO
N
N
N
S
OMe
15
Integral
0.7134
0.8559
0.8830
3.0000
2.8758
ppm 12 11 10 9 8 7 6 5 4 3 2 1

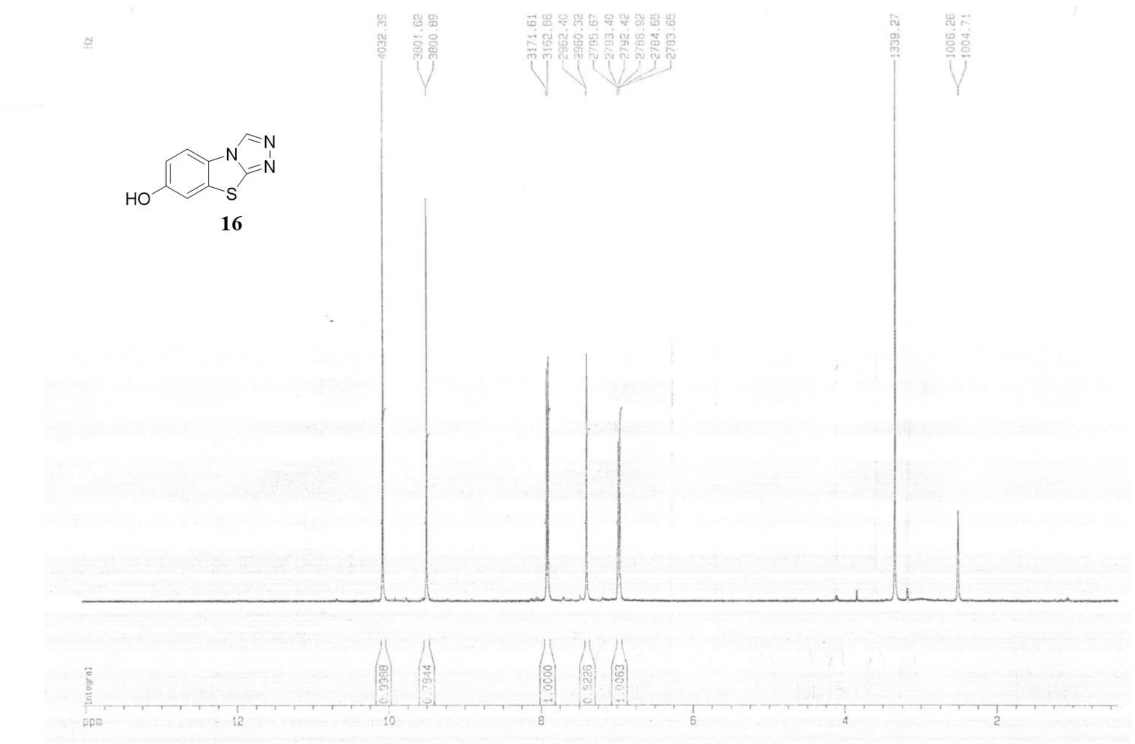
Hz
4032.39
3801.62
3800.89
3171.61
3162.86
2962.40
2960.32
2795.67
2793.40
2792.42
2786.92
2784.65
2783.65
1339.27
1006.26
1004.71
HO
N
N
N
S
16
Integral
0.9398
0.7944
1.0000
0.9226
1.0063
ppm
12
10
8
6
4
2

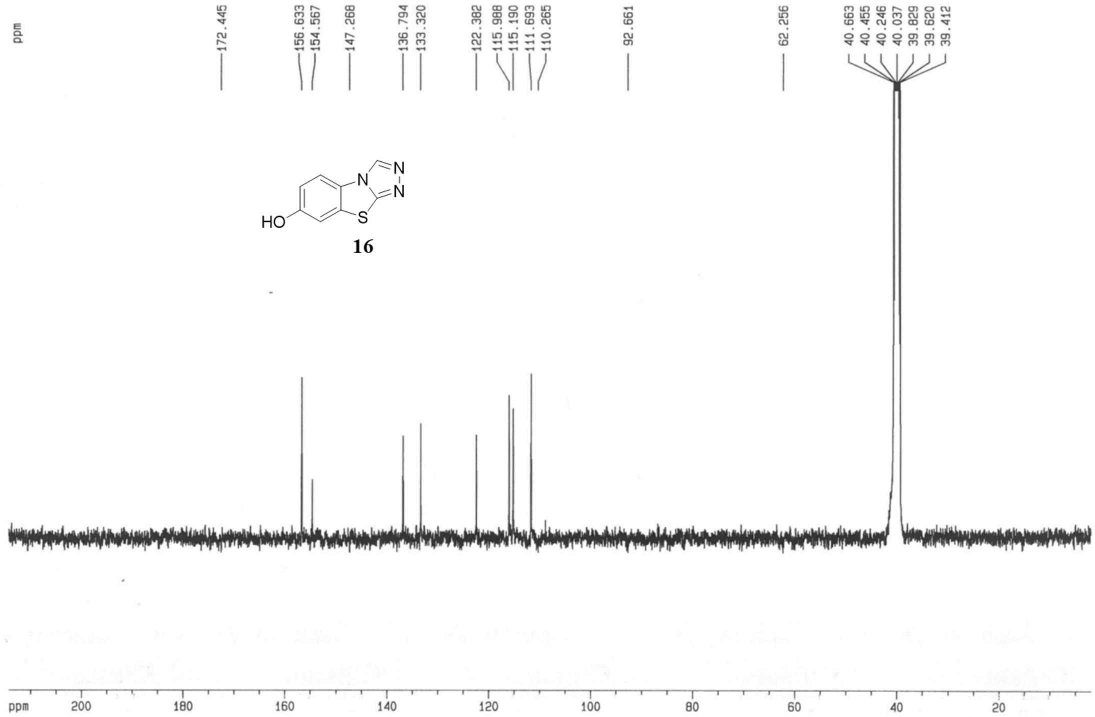
ppm
172.445
156.633
154.567
147.268
136.794
133.320
122.382
115.988
115.190
111.693
110.265
92.661
62.256
40.663
40.455
40.246
40.037
39.829
39.620
39.412
HO
N
N
N
S
16
ppm
200
180
160
140
120
100
80
60
40
20

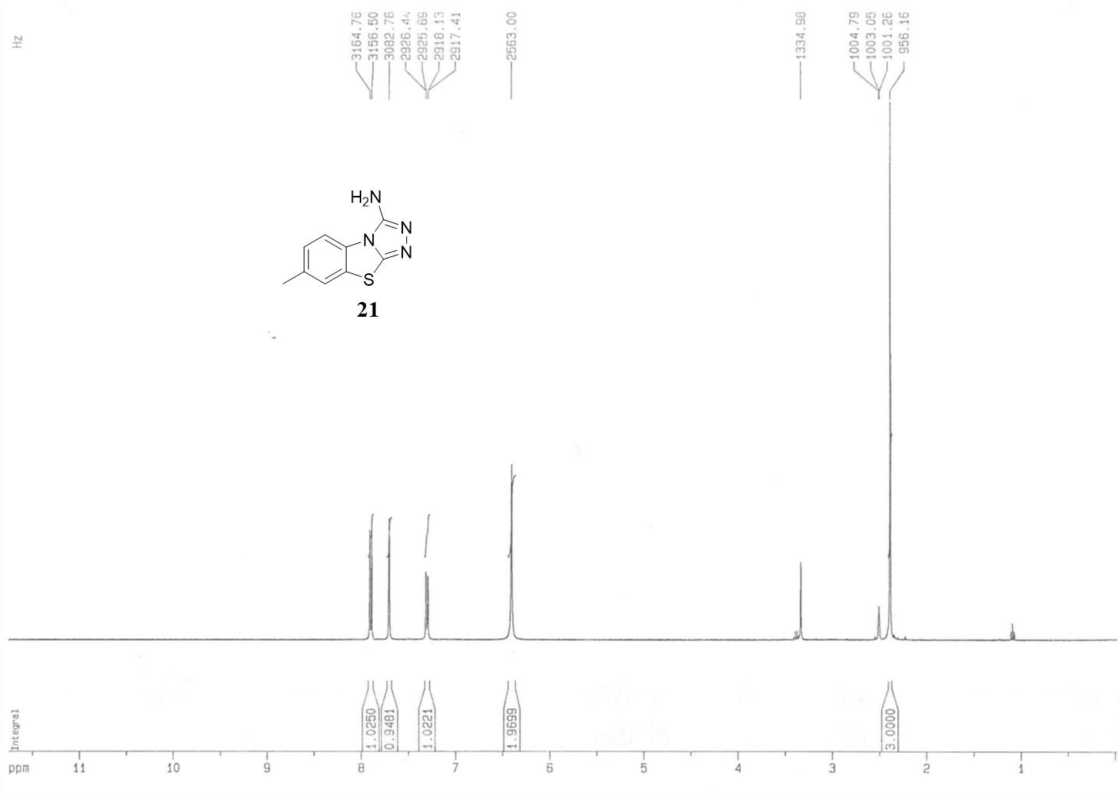

Hz
3164.76
3156.50
3082.76
2926.44
2925.69
2918.13
2917.41
2563.00
1334.98
1004.79
1003.05
1001.26
956.16
$H_2N$
N
N
N
S
21
Integral
1.0250
0.9481
1.0221
1.9699
3.0000
ppm
11
10
9
8
7
6
5
4
3
2
1


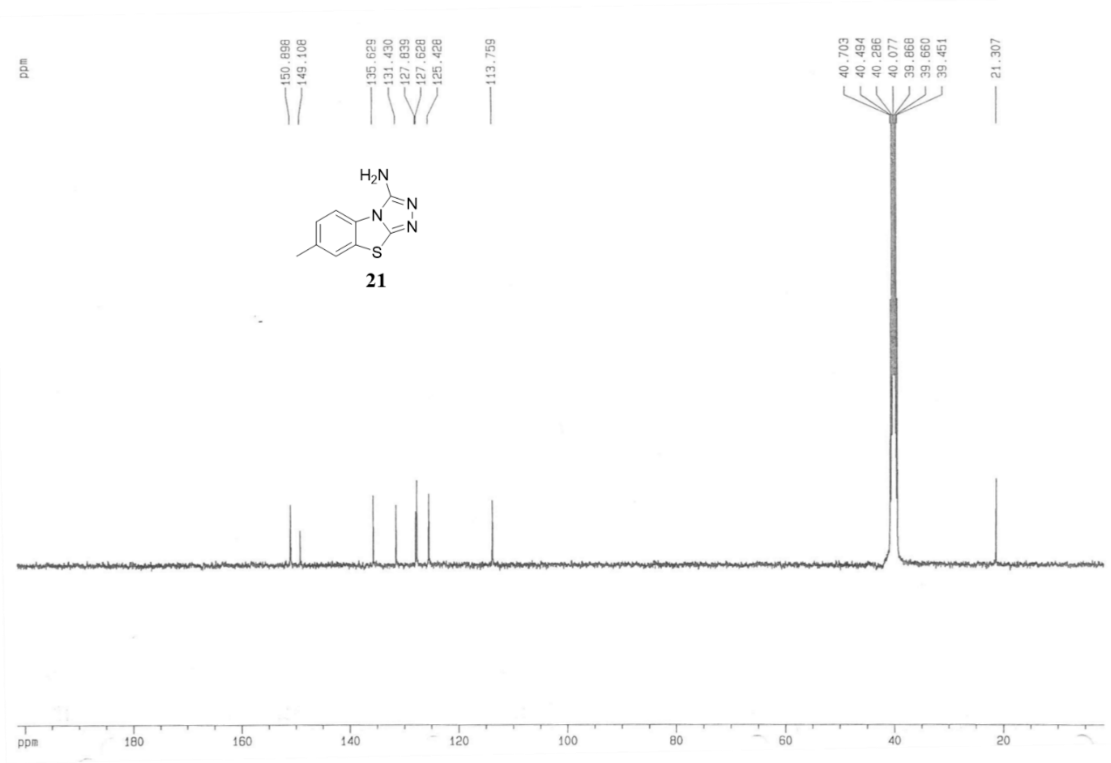

ppm
150.898
149.108
135.629
131.430
127.839
127.628
125.428
113.759
40.703
40.494
40.286
40.077
39.868
39.660
39.451
21.307
$H_2N$
N
N
N
S
21
ppm
180
160
140
120
100
80
60
40
20

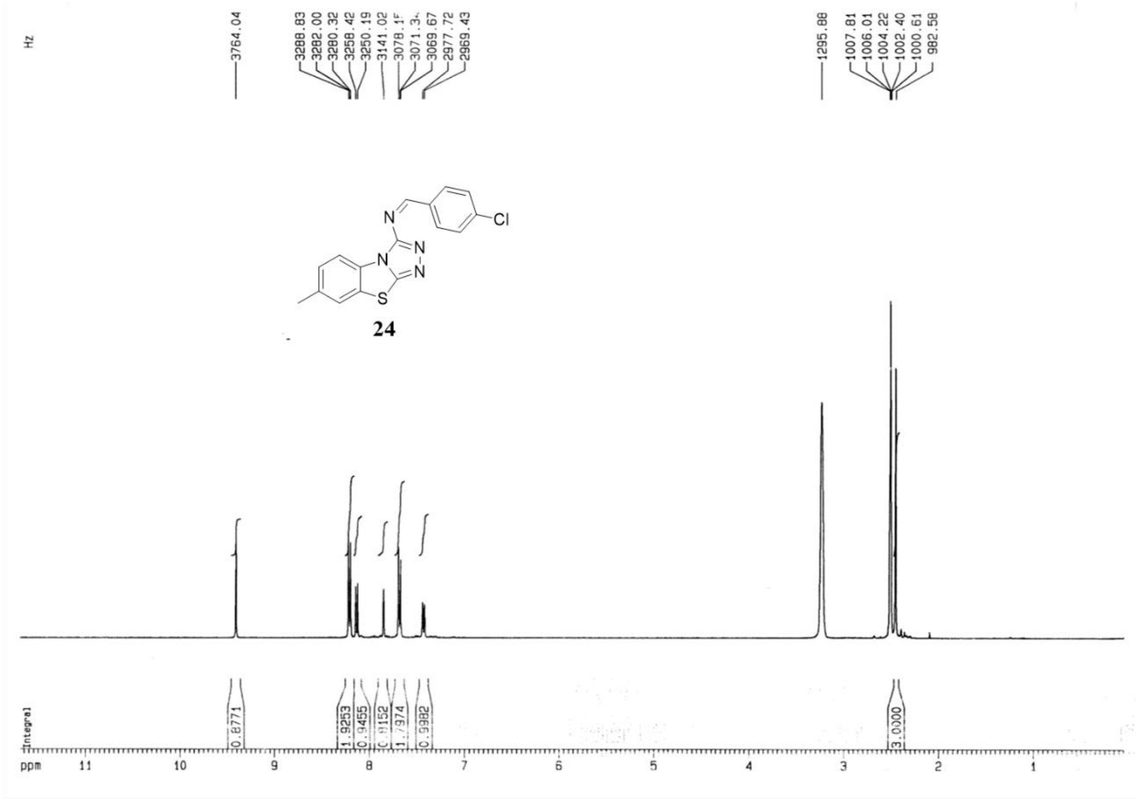
Hz
3764.04
3288.83
3282.00
3280.32
3258.42
3250.19
3141.02
3069.67
2977.72
2969.43
1295.88
1007.81
1006.01
1004.22
1002.40
1000.61
982.58
N
N
N
N
S
Cl
24
Integral
0.8771
1.9253
0.9455
0.9152
1.7974
0.9982
3.0000
ppm 11 10 9 8 7 6 5 4 3 2 1

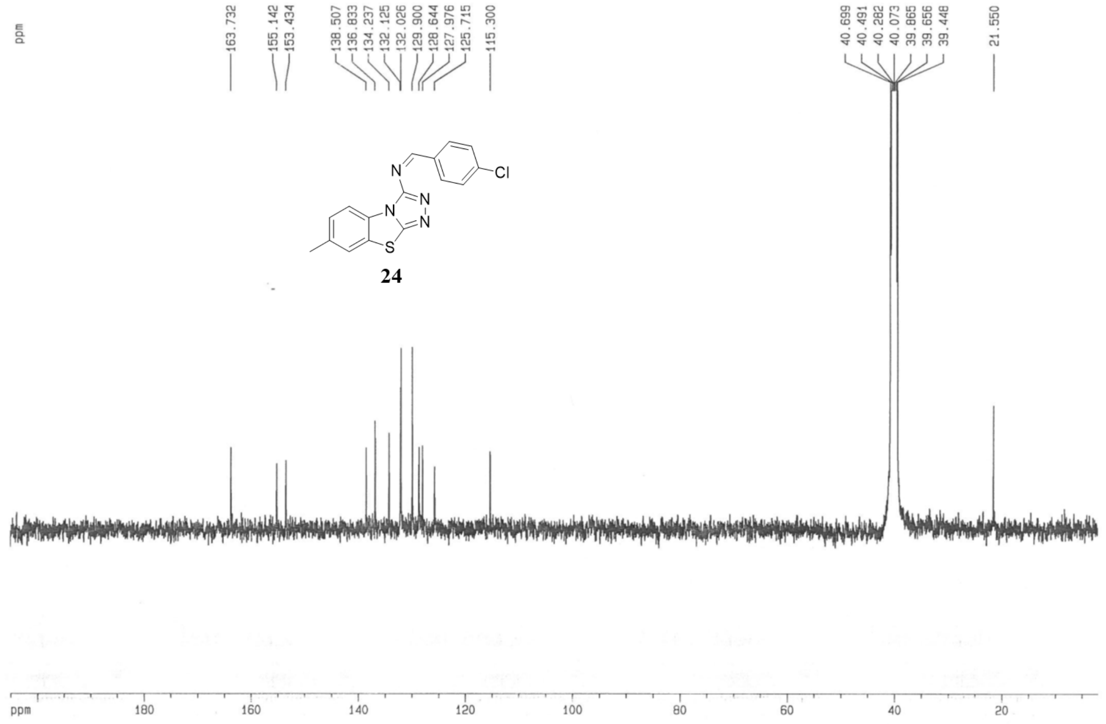
ppm
163.732
155.142
153.434
138.507
136.833
134.237
132.125
132.026
129.900
128.644
127.976
125.715
115.300
40.699
40.491
40.282
40.073
39.865
39.656
39.448
21.550
N
N
N
N
S
Cl
24
ppm 180 160 140 120 100 80 60 40 20

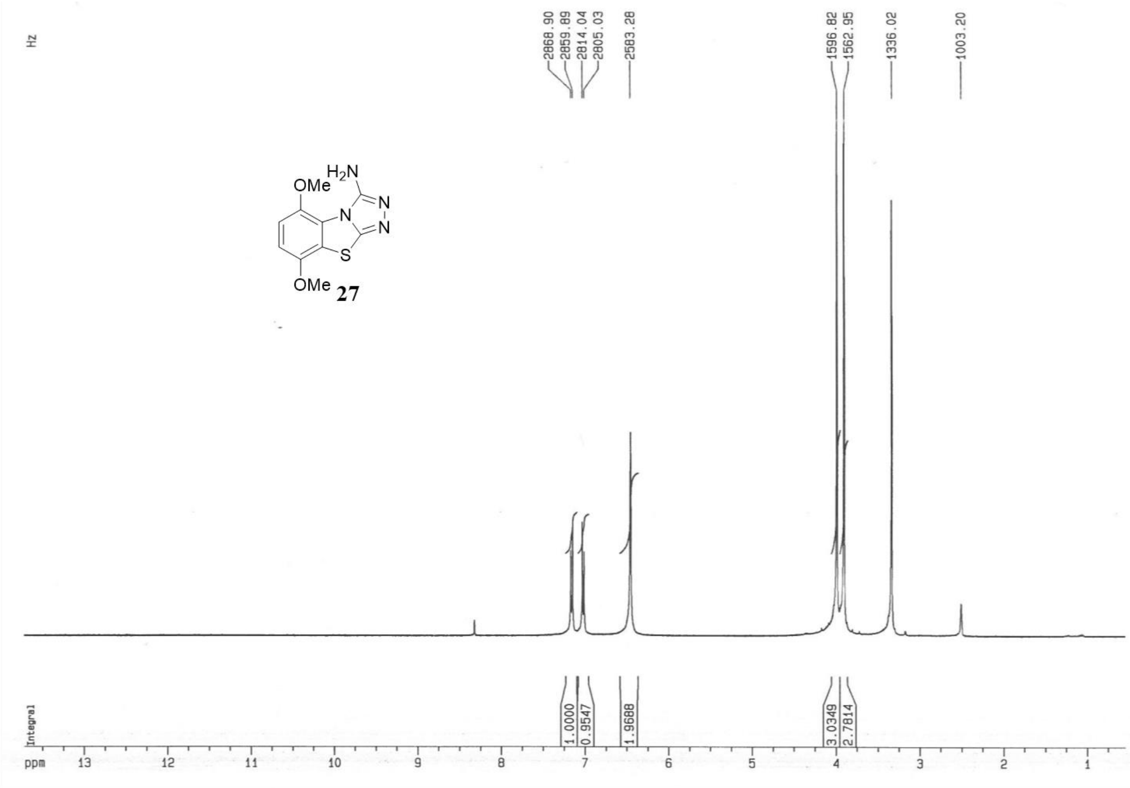
Hz
2868.90
2859.89
2814.04
2805.03
2583.28
1596.82
1562.95
1336.02
1003.20
OMe
H2N
N
N
N
S
OMe
27
Integral
1.0000
0.9547
1.9688
3.0349
2.7814
ppm 13 12 11 10 9 8 7 6 5 4 3 2 1

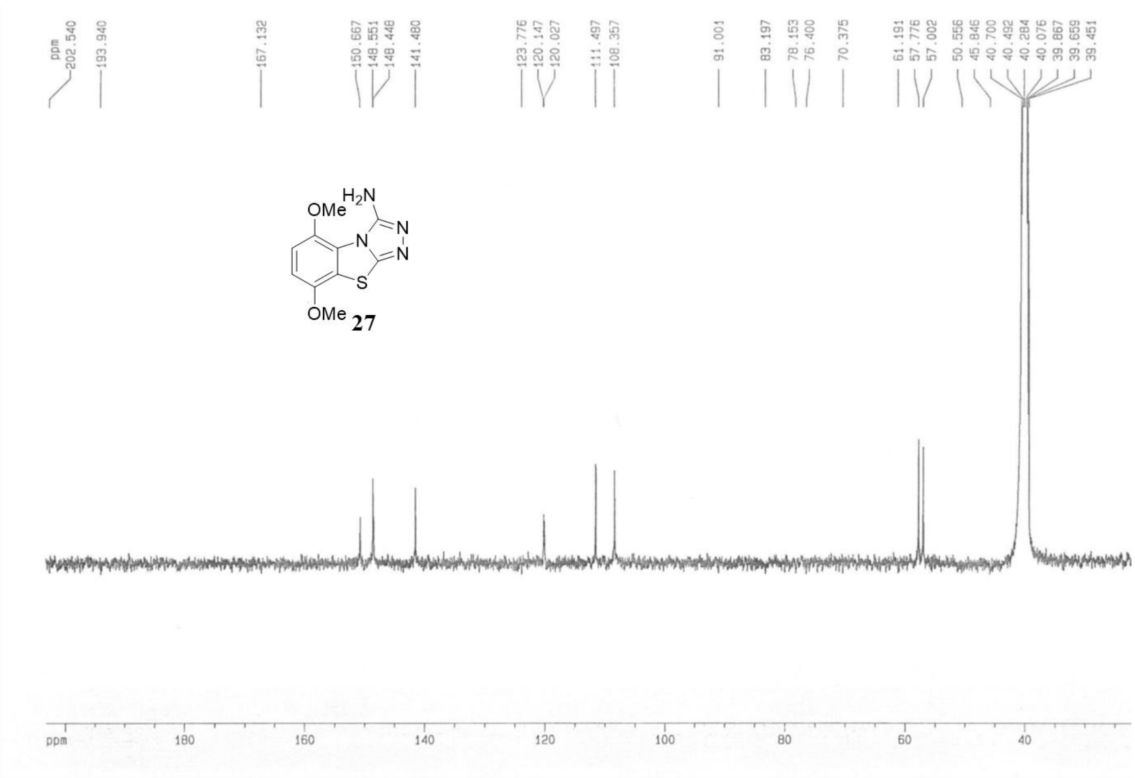
ppm
202.540
193.940
167.132
150.667
148.551
148.448
141.480
123.776
120.147
120.027
111.497
108.357
91.001
83.197
78.153
76.400
70.375
61.191
57.776
57.002
50.556
45.846
40.700
40.492
40.284
40.076
39.867
39.659
39.451
OMe
H2N
N
N
N
S
OMe
27
ppm 180 160 140 120 100 80 60 40

| **Table S2.** SMILES[1] **and biochemical data of compounds 1-31.** | | | | | |
|---|---|---|---|---|---|
| **Compd** | **SMILE** | **PARP2** | **TNKS2** | **PARP10** | **PARP15** |
| **1*** | CC1=CC=C2C(SC3=NN=CN32)=C1 | 4.0 µM | 5.3 µM | 3.2 µM | 3.2 µM |
| **2** | C1(S2)=CC=CC=C1N3C2=NN=C3 | 5.3 µM | 2.5 µM | >10 µM | >10 µM |
| **3** | CC1=CC=C(SC2=NN=CN23)C3=C1 | 1.5 µM | 0.95 µM (6.02 ± 0.13) | 10 µM | 6.6 µM |
| **4** | CC1=C(SC2=NN=CN23)C3=CC=C1 | 8.3 µM | 3.8 µM | 3.4 µM | 7.1 µM |
| **5** | CCC1=CC=C2C(SC3=NN=CN32)=C1 | 3.9 µM | 5.3 µM | 4.9 µM (5.31 ± 0.10) | 2.0 µM (5.70 ± 0.04) |
| **6** | CC(C)C1=CC=C2C(SC3=NN=CN32)=C1 | 4.5 µM | 9.5 µM | 5.5 µM (5.26 ± 0.08) | 3.9 µM (5.41 ± 0.13) |
| **7** | ClC1=CC=C2C(SC3=NN=CN32)=C1 | 2.4 µM | 10 µM | > 10 µM | 7.0 µM |
| **8#** | CC1=C2C(SC3=NN=CN32)=CC(C)=C1 | 1.3 µM | 1.2 µM | 1.7 µM | 0.78 µM (6.11 ± 0.10) |
| **9** | FC1=C2C(SC3=NN=CN32)=C(C)C=C1 | 6.9 µM | 5.4 µM | 5 µM | 6.9 µM |
| **10** | COC1=CC=C(SC2=NN=CN23)C3=C1 | 4.7 µM | 1.0 µM | 1.8 µM | 7.1 µM |
| **11** | COC1=CC=C2C(SC3=NN=CN32)=C1 | 8.8 µM | 5.5 µM | 6.5 µM | 4.0 µM |
| **12** | COC1=C(SC2=NN=CN23)C3=CC=C1 | >10 µM | 24 µM | 1.9 µM (5.71 ± 0.04) | 1.2 µM (5.93 ± 0.16) |
| **13** | COC1=C2C(SC3=NN=CN32)=CC=C1 | 2.1 µM | 1.0 µM | 5.1 µM (5.30 ± 0.15) | 2.1 µM (5.67 ± 0.03) |
| **14** | COC1=C2C(SC3=NN=CN32)=C(OC)C=C1 | 9.2 µM | 4.0 µM | 0.49 µM (6.31 ± 0.22) | 1.6 µM |
| **15** | COC1=C(SC2=NN=CN23)C3=CC(OC)=C1 | 20 µM | 7.6 µM | 1.1 µM | 1.6 µM |
| **16** | OC1=CC=C2C(SC3=NN=CN32)=C1 | 0.044 µM (7.44 ± 0.12) | 0.37 µM (6.43 ± 0.13) | > 10 µM | >10 µM |
| **17** | OC1=C(SC2=NN=CN23)C3=CC=C1 | 29 µM | 17 µM | >10 µM | 4.70 µM (5.33 ± 0.11) |
| **18** | OC1=C2C(SC3=NN=CN32)=C(O)C=C1 | 0.24 µM | 3.1 µM | 0.32 µM (6.50 ± 0.04) | 0.29 µM (6.54 ± 0.05) |
| **19** | OC1=NN=C2SC3=CC(C)=CC=C3N21 | >100 µM | >100 µM | 5.4 µM (5.27 ± 0.13) | >10 µM |
| **20** | SC1=NN=C2SC3=CC(C)=CC=C3N21 | >10 µM | >100 µM | 1.5 µM (5.81 ± 0.10) | >10 µM |
| **21** | NC1=NN=C2SC3=CC(C)=CC=C3N21 | 1.6 µM | 5.7 µM | 0.18 µM** (6.73 ± 0.12) | 0.30 µM (6.53 ± 0.14) |
| **22** | CC1=CC=C2C(SC3=NN=C(SC)N32)=C1 | 12 µM | >100 µM | 3.7 µM (5.45 ± 0.54) | >10 µM |
| **23** | CC1=CC=C2C(SC3=NN=C(SCC4=CC=C(Cl)C=C4)N32)=C1 | >100 µM | >100 µM | > 10 µM | >10 µM |
| **24** | CC1=CC=C2C(SC3=NN=C(NCC4=CC=C(Cl)C=C4)N32)=C1 | 70 µM | >100 µM | >> 10 µM | >10 µM |
| **25** | CC1=CC=C2C(SC3=NN=C(/N=C/C4=CC=C(Cl)C=C4)N32)=C1 | 2.9 µM | >100 µM | > 10 µM | 7.9 µM |
| **26** | SC1=NN=C2SC3=C(OC)C=CC(OC)=C3N21 | 19 µM | 66 µM | >> 10 µM | >>10 µM |
| **27** | NC1=NN=C2SC3=C(OC)C=CC(OC)=C3N21 | 10 µM | 10 µM | 0.12 µM** (6.93 ± 0.06) | 0.12 µM** (6.91 ± 0.03) |
| **28** | CSC1=NN=C2SC3=C(OC)C=CC(OC)=C3N21 | 8.4 µM | >100 µM | >> 10 µM | >>10 µM |

| | | | | | |
|---|---|---|---|---|---|
| **29** | O=C(C1=CC=C(Cl)C=C1)NC2=NN=C3SC4=C(OC)C=CC(OC)=C4N32 | 44 µM | 20 µM | >> 10 µM | 7.9 µM |
| **30** | CC(NC1=NN=C2SC3=C(OC)C=CC(OC)=C3N21)=O | 53 µM | >100 µM | >> 10 µM | >> 10 µM |
| **31** | O=C(C(C)(C)C)NC1=NN=C2SC3=C(OC)C=CC(OC)=C3N21 | 97 µM | >100 µM | >> 10 µM | >> 10 µM |
| *Purchased from NCI DTP repository #Purchased from Specs. >denotes less than 50% inhibition in the reported highest concentration and >> denotes no inhibition at the reported concentration. **Value limited by protein concentration used. | | | | | |
